# An evolutionary distinct Nipah G glycosylation site provides stability for receptor engagement

**DOI:** 10.1101/2025.03.31.646399

**Authors:** Tia É. Hawkins, Valeria Calvaresi, Sean A. Burnap, Liang Wu, Weston B. Struwe

## Abstract

Nipah virus is a deadly paramyxovirus with 40-75% mortality and >750 cases since 1998. Currently there are no clinically approved vaccines or therapeutics to target infection. Nipah is an enveloped virus with two surface glycoproteins, the trimeric fusion (F) and tetrameric attachment glycoprotein (G). G is responsible for cellular attachment via binding to ephrin B2/B3. Glycosylation of Nipah G and its effects on receptor engagement has not previously been studied but is important as glycosylation impacts immunogenicity, receptor binding and structural conformations for other enveloped virus glycoproteins. Our phylogenetic and mass spectrometry analysis of site-specific N-glycans of the Nipah G Malaysia strain revealed how N-glycosylation has evolved since the appearance of the virus in 1998. We discovered that the N481 N-glycosite is not conserved and although the glycan does not directly contribute to receptor binding, the threonine/serine in the glycosylation sequon is critical for maintaining long-range stability of individual G subunits that facilitates ephrin B2 binding affinity. Together, these data reveal plasticity of N-glycosylation sites across Nipah species and the presence of hydrogen bonding networks that contribute to G stability and host engagement, which is valuable information for understanding virus attachment/entry mechanisms as well as the rationale design of structure-based vaccines.

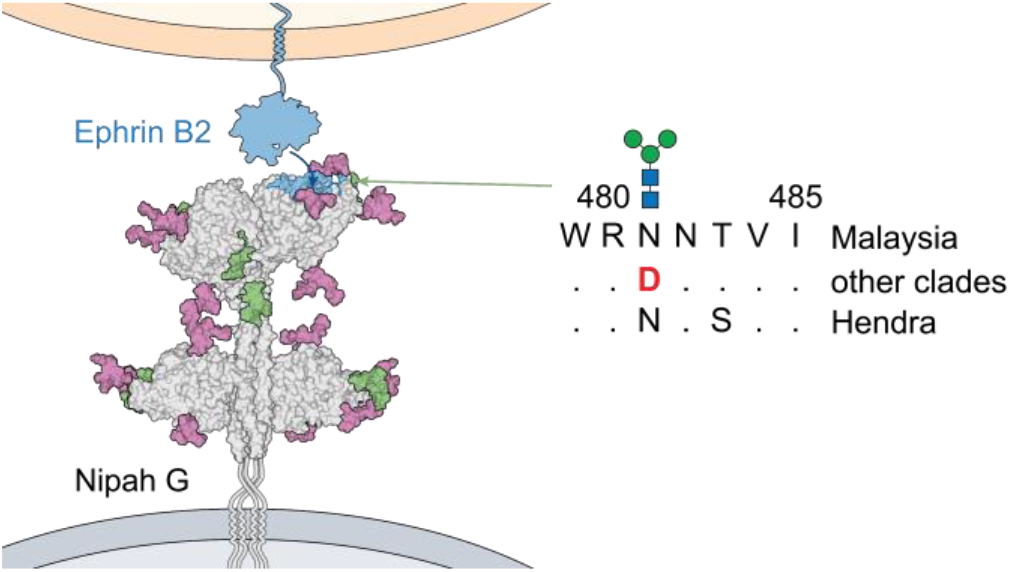

## INTRODUCTION

Nipah virus is a zoonotic paramyxovirus first reported in 1998 following outbreaks of severe encephalitis in Malaysia and Singapore^1-3^, with near-yearly outbreaks currently occurring in Bangladesh^4-6^. Nipah disease progression begins with fever, myalgia, nausea and coughing before rapid onset of encephalitis, seizures, coma and death^1,6^. The difficulty of Nipah diagnosis in the early stages of infection, coupled with a lack of approved therapeutics or vaccines, contributes to the high mortality rate (40-75%)^5^. Consequently, Nipah has been placed on the WHO R&D blueprint for emerging pathogens and epidemics^7^.

Nipah virus entry and infection are not well understood, and the lack of effective treatment regimens reflect this. The broad-spectrum antiviral ribavirin is the only drug to have been administered to infected individuals. While showing reductions in mortality, the small sample sizes and lack of control groups make it difficult to determine efficacy^8^. Currently, there are four candidate vaccines against Nipah in phase I trials, including a live attenuated (PHV02)^9^, mRNA (mRNA-1215)^10^, adenoviral vector (ChAdOx-1 NiV)^11^ and recombinant protein (HeV-G-sV)^12^ vaccines^13^. The monoclonal antibody m102.4, which targets Nipah G protein, is the only biotherapeutic to enter clinical trials^14^. Nipah and other paramyxoviruses are rapidly evolving and highly transmissible pathogens, and with a rise in vaccine hesitancy^15,16^, the long-term efficacy of these possible treatments is indeterminate. Defining conserved areas of vulnerability for Nipah will greatly aid therapeutic development against this family of viruses.

Nipah virus harbours two glycosylated surface proteins, the tetrameric attachment glycoprotein^17^ (G) that binds to the host receptor ephrin B2 or B3^18-20^ via its globular head domain and the trimeric fusion glycoprotein^21^ (F) that mediates viral fusion^22,23^ (**Fig. 1a**). While G does not display neuraminidase activity, its head domains adopt a classical neuraminidase structure, with a six-bladed beta-propeller fold^3,24,25^. F is a class I fusion protein, activated by cathepsin L and B cleavage in the endosome^26^. G binds ephrin B2 and B3 with nanomolar affinity for each head domain^18^, and picomolar affinity for the tetramer^27^ due to avidity effects. Recent data suggests the extension of a G protein head outward towards the F protein could induce the F conformational changes that lead to fusion pore formation^28^.

**Figure 1.**
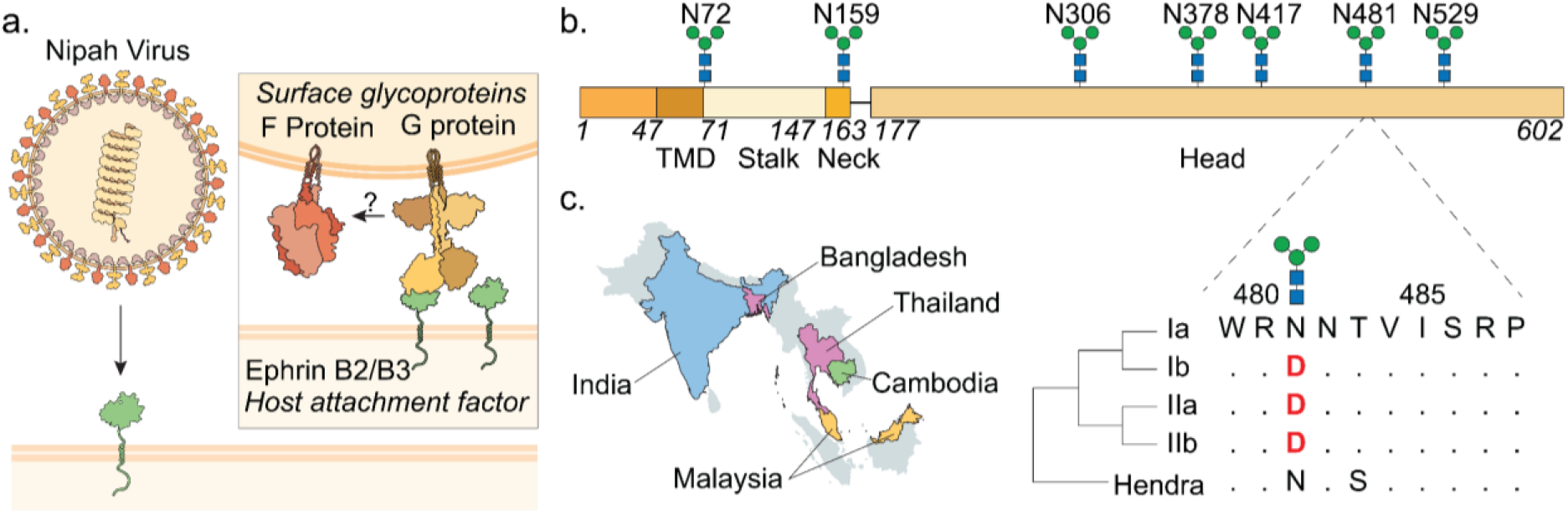
Nipah virus evolved a divergent N-glycan site. **a**) Schematic representation of Nipah virus binding host cells through ephrinB2/B3 interactions with the G protein, leading to fusion through F protein rearrangement. **b)** Nipah G_M_ predicted N-glycans sites with amino acid sequence shown. **c**) Left: Distribution of clades across location of isolates. Most prevalent clade shown for each country, identified through phylogenetic analysis (Fig S1). Clade Ia: yellow, Clade Ib: green, Clade IIa: purple, Clade IIb blue. Right: Sequence alignment of clades and Hendra outgroup for residues 479-489. Red amino acids denote a loss of N-glycosylation.

The Malaysia strain of Nipah G (G_M_) has 28 predicted N-glycan sites per tetramer, and F (F_M_) has 15 per trimer. N-glycan sites are genetically encoded through an N-X-S/T sequon and are subject to both positive and negative selection pressures^29-31^. Despite their occurrence, there is limited research into the roles of glycosylation for Nipah G^25,32^. Glycosylation of G has been suggested to be important for viral fusion^32,33^, but a site-specific characterisation of glycan structure and the associated effects on receptor engagement has not been performed. Here, we reveal the effects of Nipah G N-glycosylation on receptor binding using mass spectrometry (glycomics, glycoproteomics and hydrogen deuterium exchange (HDX)), mass photometry (MP) and biolayer interferometry (BLI) with phylogenetic analysis to uncover how a single glycan site governs structural dynamics and stability of a receptor binding complex.

## RESULTS

### Phylogenetic analysis of Nipah G N-glycosylation

The majority of research on the G protein has used the primary sequence isolated from the 1998 outbreak^3^, the so-called “Malaysia” strain. However multiple evolutionary analyses have shown that Nipah has diverged to a second strain, the Bangladesh strain^34,35^. To identify potential differences in N-glycosylation sites between strains, we aligned 81 complete sequences from the NCBI virus database using MEGA11 (Fig. S1, Supp File 3), including viruses isolated from four different host species, across five countries. In agreement with other studies, we identified distinct groupings corresponding to the Malaysia and Bangladesh strains (Clade I and II respectively in our analyses). These Clades were further classified into a and b subclades, with IIa consisting of the perennial Bangladesh outbreaks, and IIb the more recent outbreaks in Kerela, India. The location of sample collection shifted from Malaysia to Bangladesh and India over time (Tables S1-3), where most infections are now located. Notably, we found six of the seven predicted N-glycan sites were conserved between all clades (**Fig. 1b**), however, a specific mutation (N481D) resulted in the loss of an N-glycan site. This was either introduced during the split from Hendra virus or occurred independently in both Clade II and Clade Ib isolates (**Fig. 1c**). This mutation event was also independent of host and country of origin, as there were no differences in host and origin within clades themselves.

### Glycan structure and occupancy of N481

The role of the N481 glycan is unknown but it could contribute to receptor binding affinity, infectivity and/or immune detection/evasion. Critically, there is no comprehensive site-specific glycan analysis of Nipah G tetramers, which is a first step for exploring glycan structure-function relationships. Here, we used the soluble ectodomain of Nipah G_M_ (**Fig. 2**) for site-specific glycan characterisation, which was recombinantly expressed and purified from human embryonic kidney (HEK) Expi-293 cells to best mimic human glycosylation *in* vitro^36^. Purified SEC fractions of Nipah G_M_ tetramer and head domain ((Fig. S3a-b) were measured by MP, showing that soluble G_M_ formed a stable tetramer (Fig. S3c,d) as expected for Nipah G_M_ ^17^. Oligomeric assembly has been implicated in influencing glycosylation by affecting protein conformation and glycosite accessibility^37-39^. Using BLI (Fig. S4), we identified soluble G_M_ tetramer bound with picomolar affinity to recombinantly expressed soluble ephrin B2 binding domain (Fig. S3e). The Nipah monomeric head domain interacted with ephrin B2 with nanomolar affinity in a 1:1 binding model (Fig. S4b).

**Figure 2.**
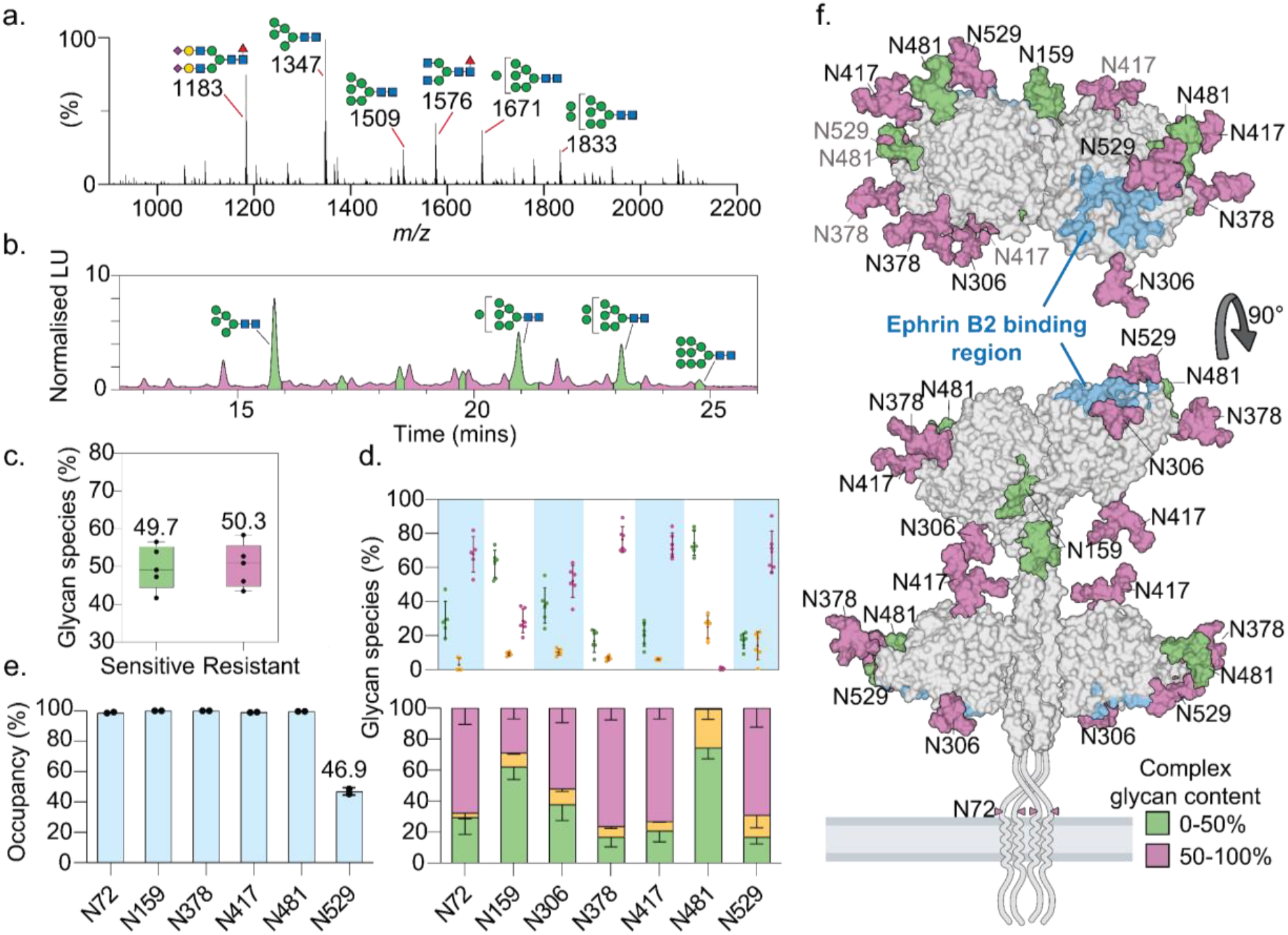
Nipah G_M_ ectodomain N-glycosylation. **a**) Total ion spectra of released N-glycans by IM-MS, scaled to most abundant ion, Man_5_GlcNAc_2_ (*m/z* 1347). **b**) Representative HPLC of 2-AA labelled N-glycans with peaks labelled as endo-H sensitive (i.e. oligomannose and hybrid) in green and endo-H resistant (i.e. complex) in purple. **c)** Mean abundance of glycan subtype (coloured as above). **d**) Glycan subtype specific for each site determined by label-free quantitative (LFQ) glycoproteomics, (n=6 and n=5 for N72). Glycans types are labelled green for oligomannose, yellow for hybrid and purple for complex. **e**) Occupancy of N-glycan sites measured by ^18^O labelled proteomics (n=2). **f**) Location of glycans on Nipah G tetramer (PDB: 7TXZ & 7TY0) modelled using GLYCAM (the most abundant glycan species per site is shown).

To identify all N-glycan structures present on G_M_, we performed ion-mobility tandem mass spectrometry (IM-MS/MS) from three biological replicates (**Fig. 2a**). A previous report identified a range of hybrid and complex type-glycans for the monomeric G_M_ head domain^25^. However, our data showed a shift towards unprocessed oligomannose structures with the tetrameric G_M_ soluble domain, with Man_5_-_8_GlcNAc_2_ (Man_5_-_8_) present as major ion species (**Fig. 2a**). Two biantennary complex type glycans were also detected as major ion species (*m/z* 1183 and 1576). Additional complex-type N-glycans, namely tri- and tetra-antennary species, were identified at low levels from ion-mobility extracted doubly and triply charged species (Fig. S5). The differences in glycan species we observed could be a result of the two stalk glycan sites or differences seen in glycan processing for the G_M_ tetramer vs monomeric head domain, as previously seen for HIV Env trimers and gp120 monomers^39^.

Next, HILIC-HPLC of 2-AA labelled N-glycans was used to quantify and confirm the relative abundance of G_M_ structures by type (**Fig. 2b**). Treatment of N-glycans with and without Endo-H identifies oligomannose and hybrid-type glycans (i.e. Endo-H sensitive) from complex-type structures (i.e. Endo-H resistant). Overall, IM-MS/MS and HPLC data agreed in that Endo-H sensitive glycans accounted for 50% of the total N-glycan pool (**Fig. 2c** and Fig. S6), with ∼75% present as Man_5_ (33.5 ± 6.2%) Man_7_ (24.5 ± 4.8%) and Man_8_ (15.1 ± 4.1%). Conversely, multiple peaks were identified as Endo-H resistant glycans across the chromatogram (i.e. pink peaks), which indicates complex-type glycan structures are diverse but at low levels compared to oligomannose structures.

Using glycoproteomics, we mapped the site-specific attachment and abundances of glycans on G_M_ from six biological replicates (**Figs. 2d**). Four of the seven N-glycan sites were predominantly complex-type, but with fewer processed oligomannose-type and hybrid structures. These sites are located at stalk site N72 (67.7 ± 9.4 %) and head sites N378 (76.5 ± 6.9 %), N417 (73.2 ± 6.4 %) and N529 (69.0 ± 11.2 %). N378 and N529 glycopeptides were primarily fucosylated (82.3 and 83.0% respectively) and, along with N72, contained the most sialyated N-glycans (37.2%, 43.0% and 35.1% respectively) (Fig. S7), consistent with the complex N-glycans identified by IM-MS/MS (Fig. S5). N417 was minimally fucosylated (4.5 ± 1.7%) and had notably lower levels of sialyation (19.2 ± 6.2%). N306 had a mixture of complex (51.9 ± 8.6%), oligomannose (37.7 % ± 9.6%) and hybrid-type (10.3 ± 1.8%) glycans. The N159 glycan, present in the neck of G_M_, contained mostly oligomannose glycans (62.1 ± 7.4%), while the evolutionary divergent site, N481, was oligomannose rich (74.4 ± 6.5%) and was the only site with negligible amounts of complex glycans (<1%). Overall, results between the three methods were in high agreement, with no significant difference between HPLC and glycoproteomics quantifications (Fig. S8).

To accurately assess N-glycosylation site occupancy on G_M_ and to support label-free quantitative (LFQ) glycoproteomics experiments, we employed differential glycopeptide Endo-H/PNGase-F digestion with ^18^O incorporation from three biological replicates (Fig. S9a)^40^. Due to using different proteolytic digest methods (in-gel digestion vs single-pot solid-phase enhanced (SP3)), we were unable to cover site N306 (see further description in the methods section, Supp File 1). Five of the six sites were ∼100% occupied, including N481 (*µ* = 99.6%, *σ* = 0.06), and only N529 was not fully occupied with 46% occupancy observed (*σ* = 2.5, *n* = 2) (**Figs. 2e,f**). For Hendra virus, this glycan is predicted to interact with ephrin B2^41^, however as N529 is not fully occupied, our data suggests it may not be the case for Nipah. To validate our glycoproteomics data, we compared the abundances of identical peptides from the same biological replicates (Table S4). There were no significant differences between the two methods (Fig. S9), suggesting ionisation efficiency between glycopeptides did not affect quantification among glycan types.

Previous research into Hendra virus showed the stalk domain of G stalk as being highly populated with O-glycans that are important for viral fusion, and some of these sites are conserved with Nipah^33^. However, Nipah’s O-glycosylation has not been investigated. Here, we identified four sites conserved between Nipah and Hendra (T114, S116, T117 and T119) were occupied with core-1 type O-glycans (Fig. S10a). In comparison to Hendra, Nipah G_M_ had fewer occupied O-glycan sites on its stalk, and these O-glycans were smaller, suggesting less overall processing of at these O-glycan sites. Three other O-glycosylation sites, which were not present on Hendra G, were identified at S380, S419 and S486, with S380 and S419 residing next to the N378 and N417 N-glycans in the sequon (Fig. S10a). S486 was O-glycosylated with a core-2 and core-3 O-glycan. T114, S116, T117 and T119 are located in the stalk region of Nipah G, at the oligomerisation interface of the two lower head domains (Fig. S10b, left), while S486 sits next to residues which interact with ephrin B2 (Fig. S10b, right). Therefore, there are conserved N- and O-glycans near the ephrin B2 binding site which could contribute to receptor binding.

### Receptor binding of Nipah G_M_ glycosite mutants

Glycans attached to viral glycoprotein glycans have been shown to influence receptor binding via direct interactions^42^ and by altering protein structural conformations^43-45^. Yet, for Nipah virus, the direct biophysical effects of G protein glycan loss on ephrin B2 binding has not been investigated, with previous studies focussing on viral fusion and cellular infectivity^32^. To explore this, we generated individual N-glycosite mutants of G_M_; T308A, S380A, S417A, T483A, T513A plus the evolutionary mutant N481D. Each N-glycan site was mutated via Ser/Thr to Ala in the sequon to prevent potential O-glycosylation of the serine or threonines^46^. The binding affinity between ephrin B2 and the head domain of each mutant was measured by BLI. The head domain was chosen over the tetramer in order to study the interaction in a 1:1 binding model (**Fig. 3** and Figs. S11,12). Furthermore, the picomolar affinity to the tetramer is at the concentration limit for BLI^47^ (Fig. S4). We also controlled for the effects of glycan structure on binding by expressing the G_M_ head in the presence of kifunensine to generate Man_8/9_ glycoforms^48^ that differ from WT glycosylation described above.

**Figure 3.**
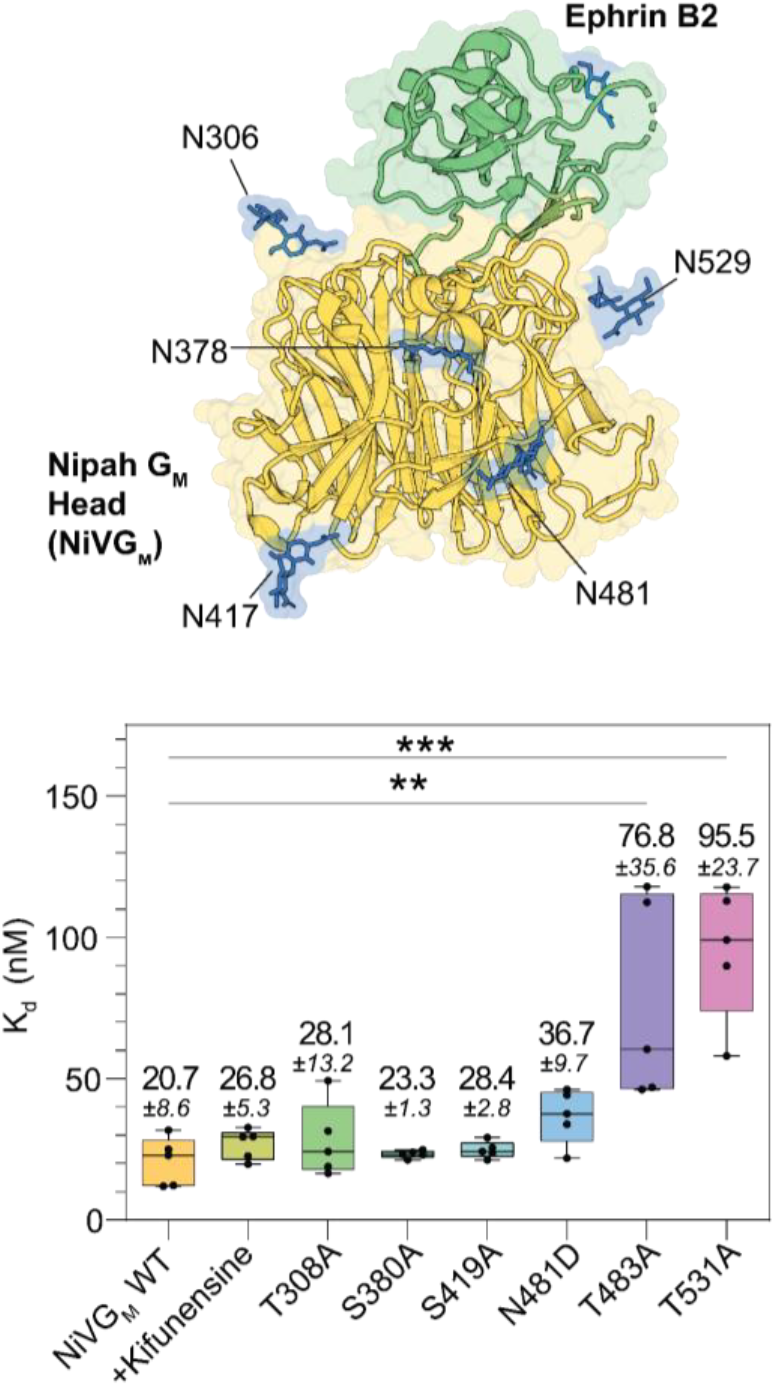
Effects of Nipah G_M_ glycan knockout mutants on ephrin B2 binding. Location of glycans on Nipah G_M_ in relation to the ephrin B2 binding interface (N-glycan chitobiose or singular GlcNAcs are shown in blue. G_M_ and ephrin B2 are yellow and green respectively). Image produced in PyMOL using PDB:2VSM, 7TY0. Mean steady state K_d_ of G_M_ mutants binding to ephrin B2 (error bars displaying standard deviation).

We did not detect any differences in K_d_ values between G_M_ WT and T308A, S380A, S419A, and N481D mutants, indicating that these N-glycans do not directly contribute to ephrin B2 binding. However, there was a 4.7-fold decrease in binding affinity with T531A (K_d_ = 95.5 ± 23.7 nM) compared to the WT (K_d_ = 20.7 ± 8.6 nM). The sidechain of T531 was previously suggested to interact through hydrophobic interactions with ephrin B2^9^, therefore, a reduction in affinity for this mutant cannot be solely attributed to N-glycan loss. T483A also exhibited a 3.7-fold reduction in binding affinity (K_d_ = 76.8 ± 35.6 nM) for ephrin B2 compared to WT. Both the evolutionary mutant N481D and T483A result in N-glycan loss at N481, but T483A elicited 2.1-fold weaker binding than N481D (K_d_ = 36.7 ± 9.7 nM). Affinity for ephrin B2 did not change with kifunensine treated G_M_, nor did the presence of an O-glycan at T483 have an effect (Fig. S13). This suggested the reduction of ephrin B2 affinity observed for T483A was not due to loss of the associated N481 glycan but was directly related to the amino acid change from threonine to alanine (**Fig. 3**).

### Structural dynamics of Nipah G_M_ and ephrin B2 complex

The loss of affinity implies a structural rearrangement of the binding site involving residues near the N481 glycan site. We used HDX MS to investigate structural dynamics associated with the N481D and T483A mutations compared to WT G_M_ head domains with and without ephrin B2 (HDX measurements were taken from 15 seconds to 24 hours). We followed the HDX of 139 peptides, covering 87% of G_M_ head domain sequence and 4 out of 5 glycosylation sites, including N481 (Figs. S14-16).

A significant decrease in HDX was measured across overlapping peptides in multiple regions for the WT G_M_ head-ephrinB2 complex (**Fig. 4a** and Fig. S14) mostly in agreement with the co-crystal structure^18^ and previously published HDX-MS data for G_M_ ectodomain-ephrin B2 complex^19^ Peptide regions that displayed significant HDX changes were localised to the ephrin B2 binding site (Fig. S17) and distributed across the oligomerisation interface, with amino acids 231-234, 455-459, 505-508 and 526-538 displaying greatest protection from HDX in the bound G_M_ form. Both 231-234 and 455-459 do not form direct contacts with ephrin B2 (Fig. S18), and this decrease in HDX is indicative of allosteric effects of rigidification in this region.

**Figure 4.**
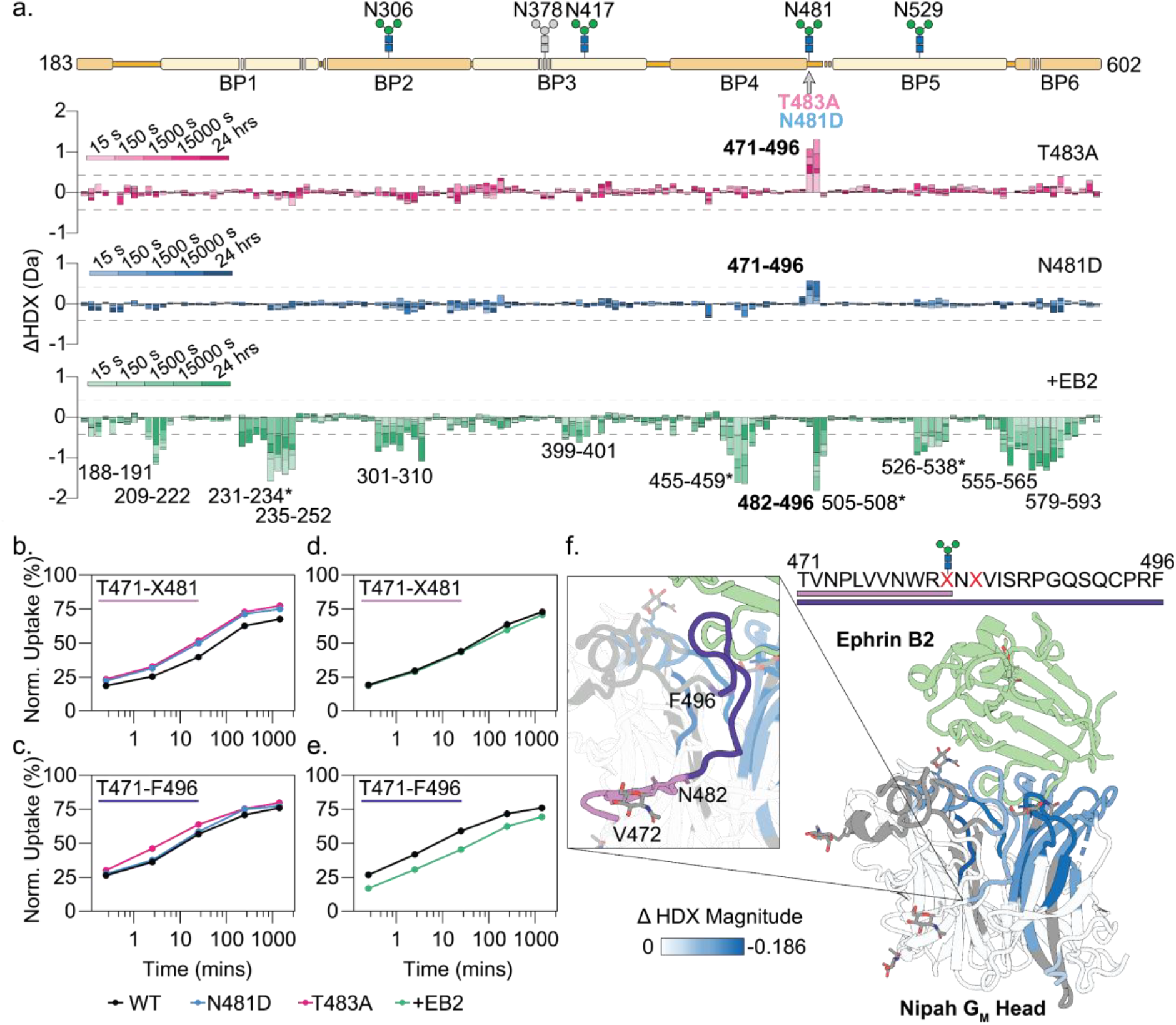
HDX-MS of Nipah G_M_ mutants +/- ephrin B2. a) Differential HDX plot of Nipah G_M_ head compared to N481D and T483A plus ephrin B2 (+EB2). Alignment of the Nipah G_M_ structure to the HDX plot (top), with the beta propellers (BP) highlighted. b) Normalised uptake plot for peptide T471-X481. c) Normalised uptake plot for peptide T471-F496. d) Normalised uptake plot for peptide T471-X481 for incubation with ephrin B2. e) Normalised uptake plot for peptide T471-F496 for incubation with ephrin B2. f) Magnitude of HDX change across the Nipah GM structure upon ephrin B2 binding is superimposed to PDB: 2VSM & 7TY0; peptides T471-X481 (light purple) and T471-F496 (indigo) are highlighted. Ephrin B2 in green, uncovered regions in grey.

Compared to the WT, both N481D and T483A mutants exhibited a significant increase in HDX localised to the mutation sites (**Fig. 4a** and Figs. S15-16), in the region comprising residues 471-481 (**Fig. 4b**). T483A also showed a clear increase in HDX across multiple time points in the longer peptide spanning 471-496, which was only seen in a single time point for N481D (**Fig. 4c**). This indicates the effect of deprotection for the T483A mutant is not only localised to the mutation site but is transmitted to the adjacent segment 482-496 (the region not covered by peptide 471-481). Interestingly, the receptor-bound complex did not display a significant change in HDX for peptide 471-481 (**Fig. 4d**), but did exhibit decreased HDX for 471-496 (**Fig. 4e**). This shows protection for residues 482-496 (**Fig. 4f**), which is the same region impacted by T483A mutant (and minimally by N481D mutant). Crucially, the residues 482-496 span a large loop in the ephrin B2 binding site (**Fig. 4f, left**). As the deuterium uptake of residues 471-496 increases over time with the unbound WT (**Fig. 4e**), its slower deuterium exchange indicates the 482-496 loop has a secondary structure. Ephrin B2 binding at this loop region results in HDX protection. HDX for the two mutants explain different binding affinities measured for ephrin B2. The T483A mutation has a greater impact on the dynamics of this loop compared to N481D, which is reflected in the greater amplitude of reduced binding affinity seen for this mutant (**Fig. 3**). Of the other N-glycan sites covered, peptides spanning the two adjacent glycosites, N306 and N529 displayed a lower HDX in the presence of ephrin B2. However, this is likely due to solvent occlusion or protein backbone interactions with the receptor.

## DISCUSSION

Here, we have identified a conserved area of vulnerability in Nipah G that impacts receptor engagement and involves an N-glycan sequon. During Nipah virus evolution, the N481 glycosylation site evolved to D481, but the following sixteen amino acids, including the T483 (S483 in Hendra), is conserved across all clades of Nipah and the primary strain for Hendra (Fig. S2). These residues form a structured loop that becomes stabilised upon ephrin B2 binding, with our HDX data highlighting that stabilisation is localised within residues 482-496. We have shown that the naturally occurring N481D mutation has minimal impact on the conformational dynamics of this loop and on receptor binding, which in principle does not impact infectivity potential. However, mutating threonine 483 to alanine significantly decreases receptor binding affinity by negatively impacting dynamics of this loop. The crystal structure of the NiVG_M_-ephrin B2 complex (PDB: 2VSM) shows the T483 hydroxyl engaging in a hydrogen bond with the carbonyl of the N543 sidechain (**Fig. 5**) and mutation of 483 to alanine would disengage this bonding, leading to conformational rearrangement of the loop and subsequent decrease in ephrin B2 binding affinity. Taken together, our data suggest that the residues spanning 483-496 are a good candidate for therapeutic targeting, as many of the other sites displaying increased protection upon receptor engagement are buried within the oligomerisation interface, and therefore potentially much less accessible to therapeutic monoclonal antibodies or antivirals.

**Figure 5.**
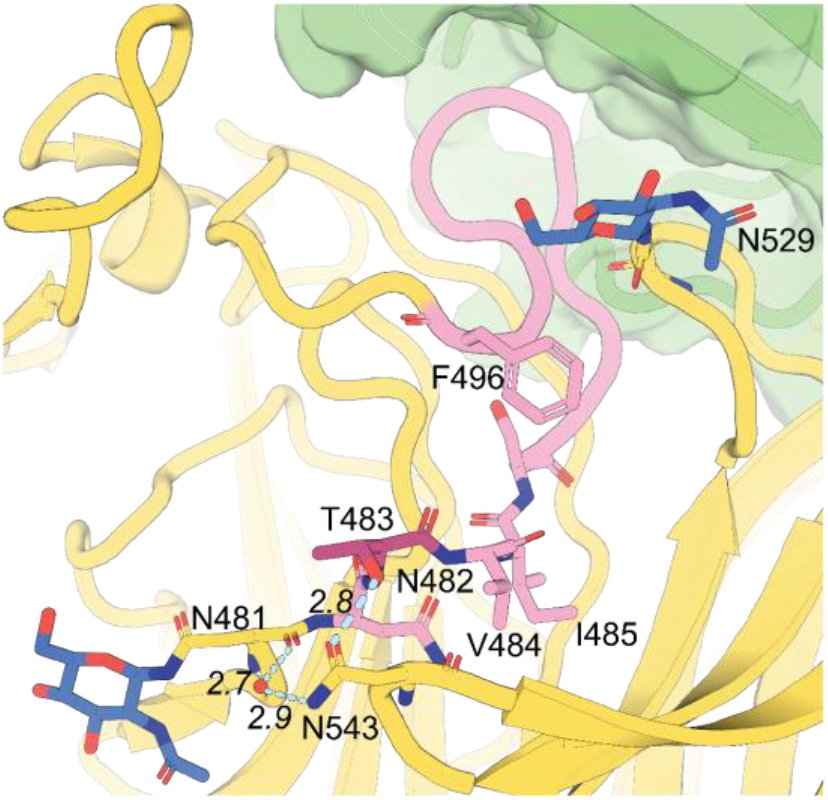
Hydrogen bonding network around N481. N482-F496 is shown in light pink, ephrin B2 is green, T483 is dark magenta. Image produced from PDB: 2VSM (distances are in Å).

To our knowledge, this is the first site-specific glycan analysis of Nipah G_M_. We identified that most sites are fully occupied with the exception of N529. However, we hypothesise that occupancy of N72 in a full-length G_M_ (i.e. on a Nipah virion) is likely low due to the close predicted proximity of this residue to the viral membrane preventing entry to the OST complex active site^49,50^. We found N159 and N481 N-glycans were oligomannose rich sites, and oligomannose-rich sites have been recently linked to immune escape for enveloped viruses, due to their ability to sample a large conformational space and bind to innate immune receptors DC-SIGN, L-SIGN, dectin-2 and Langerin^51,52^. The evolutionary variable N481 site had the greatest oligomannose occupancy, and so loss of this glycan could reduce immune pressure and clearance of Nipah. However, this does not correlate with the mortality rates seen for the two Nipah strains, where the mortality rate for the Bangladesh strain, which lacks this N-glycan, is considerably higher (∼70-90%) than the Malaysia strain (∼40%)^53^. Previous work suggests that loss of the 481 glycan through N481D increases fusogenicity in a pseudoviral model^32^. Therefore, an explanation for the non-conserved glycosite could be in response to increasing viral fusion, suggesting this region could be involved in F-protein engagement.

## CONCLUSION

Our analyses showed a single N-glycan sites on Nipah G, N481, was not conserved across known isolates. Through performing the first photogenic and glycan site-specific analyses of Nipah G glycosylation, we determined that this site has an abundance of oligomannose structures. Finally, we discovered that a conserved loop region proximal to N481 is required for ephrin B2 binding and suggest it as an ideal target for therapeutic development and/or targeting for Nipah virus.

## Supporting information

Supporting File 2. HDX-MS uptake plots

Supporting File 3. HDX-MS reporting form (Mutants)

Supporting File 4. HDX-MS reporting form (Ephrin B2 complex)

Supporting File 5. HDX-MS individual deuterium uptake values

## Data availability

Glycoproteomics and hydrogen deuterium exchange data are deposited to ProteomeXchange Consortium (http://proteomecentral.proteomexchange.org) via the PRIDE partner repository^54^ with the dataset identifier PXD062381. In line with hydrogen deuterium mass spectrometry reporting guidelines^55^, HDX-MS uptake plots, uptake values and reporting forms are attached as Supplementary files (Supp Files 2-5). All other data available upon request.

## Acknowledgements

We would like to thank Joseph Thrush and Raymond Owens for the kind gifts of NiVG 71-602, NiVG 183-602, ephrin B2 28-165 and BirA DNA constructs. We would also like Shabaz Mohammed and Yana Demanyenko for useful discussion on SP3 digestion, O^18^ labelling and proteomic analysis. This work was supported by funding from the UK Research and Innovation Future Leaders Fellowship (MR/V02213*X*/1 to WBS) and the Wellcome Trust Sir Henry Dale Fellowship (218579/Z/19/Z to LW). The Rosalind Franklin Institute is funded by UK Research and Innovation through the Engineering and Physical Sciences Research Council (EPSRC).

## Author contributions

TÉH, LW and WBS conceptualised experiments. TÉH and SAB conducted and analysed glycan HPLC. SAB conducted and analysed IM-MS/MS of glycans with input from WBS. TÉH and VC conducted and analysed HDX-MS experiments. TÉH conducted and analysed all other experiments. TÉH wrote the manuscript with input from all authors.

## Supporting Information

Supporting File 1. Methods and supporting figures and tables

Supporting File 2. HDX-MS uptake plots

Supporting File 3. HDX-MS reporting form (mutants)

Supporting File 4. HDX-MS reporting form (ephrin B2 complex)

Supporting File 5. HDX-MS individual deuterium uptake values

## Competing interests

WBS is a shareholder and consultant to Refeyn Ltd.

## Notes

### Competing Interest Statement

The authors have declared no competing interest.

## REFERENCES

(1) Chua, K. B.; Goh, K. J.; Wong, K. T.; Kamarulzaman, A.; Tan, P. S.; Ksiazek, T. G.; Zaki, S. R.; Paul, G.; Lam, S. K.; Tan, C. T. Fatal encephalitis due to Nipah virus among pig-farmers in Malaysia. Lancet 1999, 354 (9186), 1257–1259. DOI: 10.1016/S0140-6736(99)04299-3.

(2) Harcourt, B. H.; Tamin, A.; Ksiazek, T. G.; Rollin, P. E.; Anderson, L. J.; Bellini, W. J.; Rota, P. Molecular characterization of Nipah virus, a newly emergent paramyxovirus. Virology 2000, 271 (2), 334–349. DOI: 10.1006/viro.2000.0340.

(3) Chua, K. B.; Bellini, W. J.; Rota, P. A.; Harcourt, H.; Tamin, A.; Lam, S. K.; Ksiazek, T. G.; Rollin, P. E.; Zaki, S. R.; Shieh, W.; et al. Nipah virus: a recently emergent deadly paramyxovirus. Science 2000, 288 (5470), 1432–1435. DOI: 10.1126/science.288.5470.1432.

(4) Lo, M. K.; Lowe, L.; Hummel, K. B.; Sazzad, H. M. S.; Gurley, E. S.; Hossain, M. J.; Luby, S. P.; Miller, D. M.; Comer, J. A.; Rollin, P. E.; et al. Characterization of Nipah virus from outbreaks in Bangladesh, 2008-2010. Emerg Infect Dis 2012, 18 (2), 248–255. DOI: 10.3201/eid1802.111492.

(5) Nazmunnahar; Ahmed, I.; Roknuzzaman, A. S. M.; Islam, M. R. The recent Nipah virus outbreak in Bangladesh could be a threat for global public health: A brief report. Health Science Reports 2023, 6 (7), e1423. DOI: 10.1002/hsr2.1423

(6) Hsu, V. P.; Hossain, M. J.; Parashar, U. D.; Ali, M. M.; Ksiazek, T. G.; Kuzmin, I.; Niezgoda, M.; Rupprecht, C.; Bresee, J.; Breiman, R. F. Nipah virus encephalitis reemergence, Bangladesh. Emerg Infect Dis 2004, 10 (12), 2082–2087. DOI: 10.3201/eid1012.040701.

(7) Sweileh, W. M. Global research trends of World Health Organization’s top eight emerging pathogens. Global Health 2017, 13 (1), 9. DOI: 10.1186/s12992-017-0233-9.

(8) Banerjee, S.; Niyas, V. K. M.; Soneja, M.; Shibeesh, A. P.; Basheer, M.; Sadanandan, R.; Wig, N.; Biswas, A. First experience of ribavirin postexposure prophylaxis for Nipah virus, tried during the 2018 outbreak in Kerala, India. Journal of Infection 2019, 78 (6). DOI: 10.1016/j.jinf.2019.03.005.

(9) Monath, T. P.; Nichols, R.; Feldmann, F.; Griffin, A.; Haddock, E.; Callison, J.; Meade-White, K.; Okumura, A.; Lovaglio, J.; Hanley, P. W.; et al. Immunological correlates of protection afforded by PHV02 live, attenuated recombinant vesicular stomatitis virus vector vaccine against Nipah virus disease. Front Immunol 2023, 14. DOI: 10.3389/fimmu.2023.1216225.

(10) Loomis, R. J.; DiPiazza, A. T.; Falcone, S.; Ruckwardt, T. J.; Morabito, K. M.; Abiona, O. M.; Chang, L. A.; Caringal, R. T.; Presnyak, V.; Narayanan, E.; et al. Chimeric Fusion (F) and Attachment (G) Glycoprotein Antigen Delivery by mRNA as a Candidate Nipah Vaccine. Front Immunol 2021, 12. DOI: 10.3389/fimmu.2021.772864.

(11) Doremalen, N. v.; Avanzato, V. A.; Goldin, K.; Feldmann, F.; Schulz, J. E.; Haddock, E.; Okumura, A.; Lovaglio, J.; Hanley, P. W.; Cordova, K.; et al. ChAdOx1 NiV vaccination protects against lethal Nipah Bangladesh virus infection in African green monkeys. NPJ Vaccines 2022, 7 (1). DOI: 10.1038/s41541-022-00592-9.

(12) Eldridge, J. H.; Egan, M. A.; Matassov, D.; Hamm, S.; Hermida, L.; Chen, T.; Tremblay, M.; Sciotto-Brown, S.; Xu, R.; Dimitrov, A.; et al. A Brighton Collaboration standardized template with key considerations for a benefit/risk assessment for a soluble glycoprotein vaccine to prevent disease caused by Nipah or Hendra viruses. Vaccine 2021, 39 (38). DOI: 10.1016/j.vaccine.2021.07.030.

(13) Rodrigue, V.; Gravagna, K.; Yao, J.; Nafade, V.; Basta, N. E. Current progress towards prevention of Nipah and Hendra disease in humans: A scoping review of vaccine and monoclonal antibody candidates being evaluated in clinical trials. Tropical Medicine & International Health 2024, 29 (5). DOI: 10.1111/tmi.13979.

(14) EG, P.; T, M.; SM, M.; S, E.; M, G.; KL, H.; ML, J.; P, G.; KD, L.; H, C.; et al. Safety, tolerability, pharmacokinetics, and immunogenicity of a human monoclonal antibody targeting the G glycoprotein of henipaviruses in healthy adults: a first-in-human, randomised, controlled, phase 1 study - PubMed. The Lancet. Infectious diseases 2020, 20 (4). DOI: 10.1016/S1473-3099(19)30634-6.

(15) Nowak, G. J.; Cacciatore, M. A. State of Vaccine Hesitancy in the United States. Pediatric Clinics of North America 2023, 70 (2). DOI: 10.1016/j.pcl.2022.11.001.

(16) He, K.; Mack, W. J.; Neely, M.; Lewis, L.; Anand, V.; He, K.; Mack, W. J.; Neely, M.; Lewis, L.; Anand, V. Parental Perspectives on Immunizations: Impact of the COVID-19 Pandemic on Childhood Vaccine Hesitancy. Journal of Community Health 2021 47:1 2021, 47 (1). DOI: 10.1007/s10900-021-01017-9.

(17) Wang, Z.; Amaya, M.; Addetia, A.; Dang, H. V.; Reggiano, G.; Yan, L.; Hickey, A. C.; DiMaio, F.; Broder, C. C.; Veesler, D. Architecture and antigenicity of the Nipah virus attachment glycoprotein. Science 2022, 375 (6587), 1373–1378. DOI: 10.1126/science.abm5561.

(18) Bowden, T. A.; Aricescu, A. R.; Gilbert, R. J. C.; Grimes, J. M.; Jones, E. Y.; Stuart, D. I. Structural basis of Nipah and Hendra virus attachment to their cell-surface receptor ephrin-B2. Nat Struct Mol Biol 2008, 15 (6), 567–572. DOI: 10.1038/nsmb.1435.

(19) Wong, J. J. W.; Young, T. A.; Zhang, J.; Liu, S.; Leser, G. P.; Komives, E. A.; Lamb, R. A.; Zhou, Z. H.; Salafsky, J.; Jardetzky, T. S. Monomeric ephrinB2 binding induces allosteric changes in Nipah virus G that precede its full activation. Nat Commun 2017, 8 (1), 781. DOI: 10.1038/s41467-017-00863-3.

(20) Bonaparte, M. I.; Dimitrov, A. S.; Bossart, K. N.; Crameri, G.; Mungall, B. A.; Bishop, K. A.; Choudhry, V.; Dimitrov, D. S.; Wang, L.-F.; Eaton, B. T.; et al. Ephrin-B2 ligand is a functional receptor for Hendra virus and Nipah virus. Proceedings of the National Academy of Sciences 2005, 102 (30), 10652–10657. DOI: 10.1073/pnas.0504887102

(21) Xu, K.; Chan, Y.-P.; Bradel-Tretheway, B.; Akyol-Ataman, Z.; Zhu, Y.; Dutta, S.; Yan, L.; Feng, Y.; Wang, L.-F.; Skiniotis, G.; et al. Crystal Structure of the Pre-fusion Nipah Virus Fusion Glycoprotein Reveals a Novel Hexamer-of-Trimers Assembly. PLOS Pathogens 2015, 11 (12), e1005322. DOI: 10.1371/journal.ppat.1005322

(22) Aguilar, H. C.; Aspericueta, V.; Robinson, L. R.; Aanensen, K. E.; Lee, B. A Quantitative and Kinetic Fusion Protein-Triggering Assay Can Discern Distinct Steps in the Nipah Virus Membrane Fusion Cascade. Journal of Virology 2010, 84 (16), 8033–8041. DOI: 10.1128/jvi.00469-10

(23) Liu, Q.; Stone, J. A.; Bradel-Tretheway, B.; Dabundo, J.; Benavides Montano, J. A.; Santos-Montanez, J.; Biering, S. B.; Nicola, A. V.; Iorio, R. M.; Lu, X.; et al. Unraveling a three-step spatiotemporal mechanism of triggering of receptor-induced Nipah virus fusion and cell entry. PLOS Pathogens 2013, 9 (11), e1003770. DOI: 10.1371/journal.ppat.1003770.

(24) K, M.; P, S.; P, H.; A, H.; A, G.; L, G.; H, W.; L, H.; L, S.; B, R. A morbillivirus that caused fatal disease in horses and humans - PubMed. Science 1995, 268 (5207). DOI: 10.1126/science.7701348.

(25) Bowden, T. A.; Crispin, M.; Harvey, D. J.; Aricescu, A. R.; Grimes, J. M.; Jones, E. Y.; Stuart, D. I. Crystal structure and carbohydrate analysis of Nipah virus attachment glycoprotein: a template for antiviral and vaccine design. Journal of Virology 2008, 82 (23), 11628–11636. DOI: 10.1128/JVI.01344-08.

(26) Wang, Q.; Liu, J.; Luo, Y.; Kliemke, V.; Matta, G. L.; Wang, J.; Liu, Q. The nanoscale organization of the Nipah virus fusion protein informs new membrane fusion mechanisms. eLife 2024, 13. DOI: 10.7554/eLife.97017.2.

(27) Negrete, O. A.; Chu, D.; Aguilar, H. C.; Lee, B. Single Amino Acid Changes in the Nipah and Hendra Virus Attachment Glycoproteins Distinguish EphrinB2 from EphrinB3 Usage. Journal of Virology 2007, 81 (19), 10804–10814. DOI: 10.1128/JVI.00999-07

(28) Fan, P.; Sun, M.; Zhang, X.; Zhang, H.; Liu, Y.; Yao, Y.; Li, M.; Fang, T.; Sun, B.; Chen, Z.; et al. A potent Henipavirus cross-neutralizing antibody reveals a dynamic fusion-triggering pattern of the G-tetramer. Nat Commun 2024, 15 (1), 4330. DOI: 10.1038/s41467-024-48601-w

(29) Alves, I.; Fernandes, Â.; Santos-Pereira, B.; Azevedo, C. M.; Pinho, S. S. Glycans as a key factor in self and nonself discrimination: impact on the breach of immune tolerance. FEBS Lett 2022, 596 (12), 1485–1502. DOI: 10.1002/1873-3468.14347

(30) Miller, N. L.; Clark, T.; Raman, R.; Sasisekharan, R. Glycans in Virus-Host Interactions: A Structural Perspective. Frontiers in Molecular Biosciences 2021, 8, 666756. DOI: 10.3389/fmolb.2021.666756

(31) Kobayashi, Y.; Suzuki, Y. Evidence for N-Glycan Shielding of Antigenic Sites during Evolution of Human Influenza A Virus Hemagglutinin. Journal of Virology 2012, 86 (7), 3446–3451. DOI: 10.1128/jvi.06147-11

(32) Biering, S. B.; Huang, A.; Vu, A. T.; Robinson, L. R.; Bradel-Tretheway, B.; Choi, E.; Lee, B.; Aguilar, H. C. N-Glycans on the Nipah Virus Attachment Glycoprotein Modulate Fusion and Viral Entry as They Protect against Antibody Neutralization. Journal of Virology 2012, 86 (22), 11991. DOI: 10.1128/JVI.01304-12

(33) Stone, J. A.; Nicola, A. V.; Baum, L. G.; Aguilar, H. C. Multiple Novel Functions of Henipavirus O-glycans: The First O-glycan Functions Identified in the Paramyxovirus Family. PLOS Pathogens 2016, 12 (2). DOI: 10.1371/journal.ppat.1005445.

(34) de Campos, G. M.; Cella, E.; Kashima, S.; Alcântara, L. C. J.; Sampaio, S. C.; Elias, M. C.; Giovanetti, M.; Slavov, S. N. Updated Insights into the Phylogenetics, Phylodynamics, and Genetic Diversity of Nipah Virus (NiV). Viruses 2024, 16 (2), 171. DOI: 10.3390/v16020171.

(35) Shi, J.; Sun, J.; Hu, N.; Hu, Y. Phylogenetic and genetic analyses of the emerging Nipah virus from bats to humans. Infect Genet Evol 2020, 85, 104442. DOI: 10.1016/j.meegid.2020.104442.

(36) Goh, J. B.; Ng, S. K. Impact of host cell line choice on glycan profile. Critical Reviews in Biotechnology 2018, 38 (6). DOI: 10.1080/07388551.2017.1416577.

(37) Rudd, P. M.; Dwek, R. A. Glycosylation: Heterogeneity and the 3D Structure of Proteins. Critical Reviews in Biochemistry and Molecular Biology 1997, 32 (1). DOI: 10.3109/10409239709085144.

(38) Fercher, C.; Zacchi, L. F. Resolving the TorsinA Oligomerization Conundrum: The Glycan Hypothesis. Frontiers in Molecular Biosciences 2020, 7. DOI: 10.3389/fmolb.2020.585643.

(39) Struwe, W. B.; Chertova, E.; Allen, J. D.; Seabright, G. E.; Watanabe, Y.; Harvey, D. J.; Medina-Ramirez, M.; Roser, J. D.; Smith, R.; Westcott, D.; et al. Site-Specific Glycosylation of Virion-Derived HIV-1 Env Is Mimicked by a Soluble Trimeric Immunogen. Cell Rep 2018, 24 (8), 1958-1966.e1955. DOI: 10.1016/j.celrep.2018.07.080.

(40) Cao, L.; Diedrich, J. K.; Ma, Y.; Wang, N.; Pauthner, M.; Park, S.-K. R.; Delahunty, C. M.; McLellan, J. S.; Burton, D. R.; Yates, J. R.; et al. Global site-specific analysis of glycoprotein N-glycan processing. Nat Protoc 2018, 13 (6), 1196–1212. DOI: 10.1038/nprot.2018.024.

(41) Colgrave, M. L.; Snelling, H. J.; Shiell, B. J.; Feng, Y.-R.; Chan, Y.-P.; Bossart, K. N.; Xu, K.; Nikolov, D. B.; Broder, C. C.; Michalski, W. P. Site occupancy and glycan compositional analysis of two soluble recombinant forms of the attachment glycoprotein of Hendra virus. Glycobiology 2012, 22 (4). DOI: 10.1093/glycob/cwr180.

(42) St-Pierre, C.; Manya, H.; Ouellet, M.; Clark, G. F.; Endo, T.; Tremblay, M. J.; Sato, S. Host-Soluble Galectin-1 Promotes HIV-1 Replication through a Direct Interaction with Glycans of Viral gp120 and Host CD4. Journal of Virology 2011, 85 (22). DOI: 10.1128/jvi.05351-11.

(43) Casalino, L.; Gaieb, Z.; Goldsmith, J. A.; Hjorth, C. K.; Dommer, A. C.; Harbison, A. M.; Fogarty, C. A.; Barros, E. P.; Taylor, B. C.; McLellan, J. S.; et al. Beyond Shielding: The Roles of Glycans in the SARS-CoV-2 Spike Protein. ACS Cent Sci 2020, 6 (10), 1722–1734. DOI: 10.1021/acscentsci.0c01056.

(44) Li, Y.; Luo, L.; Rasool, N.; Kang, C. Y. Glycosylation is necessary for the correct folding of human immunodeficiency virus gp120 in CD4 binding. Journal of Virology 1993, 67 (1). DOI: 10.1128/jvi.67.1.584-588.1993.

(45) Mathys, L.; François, K. O.; Quandte, M.; Braakman, I.; Balzarini, J. Deletion of the Highly Conserved N-Glycan at Asn260 of HIV-1 gp120 Affects Folding and Lysosomal Degradation of gp120, and Results in Loss of Viral Infectivity. PLoS ONE 2014, 9 (6). DOI: 10.1371/journal.pone.0101181.

(46) Trinidad, J. C.; Schoepfer, R.; Burlingame, A. L.; Medzihradszky, K. F. N-and O-Glycosylation in the Murine Synaptosome. Molecular & Cellular Proteomics 2013, 12 (12). DOI: 10.1074/mcp.M113.030007.

(47) Kamat, V.; Rafique, A. Designing binding kinetic assay on the bio-layer interferometry (BLI) biosensor to characterize antibody-antigen interactions. Analytical Biochemistry 2017, 536. DOI: 10.1016/j.ab.2017.08.002.

(48) Elbein, A. D.; Tropea, J. E.; Mitchell, M.; Kaushal, G. P. Kifunensine, a potent inhibitor of the glycoprotein processing mannosidase I. The Journal of Biological Chemistry 1990, 265 (26), 15599–15605.

(49) Alcorlo, M.; Dik, D. A.; De Benedetti, S.; Mahasenan, K. V.; Lee, M.; Domínguez-Gil, T.; Hesek, D.; Lastochkin, E.; López, D.; Boggess, B.; et al. Structural basis of denuded glycan recognition by SPOR domains in bacterial cell division. Nat Commun 2019, 10 (1), 5567. DOI: 10.1038/s41467-019-13354-4.

(50) Bañó-Polo, M.; Baldin, F.; Tamborero, S.; Marti-Renom, M. A.; Mingarro, I. N-glycosylation efficiency is determined by the distance to the C-terminus and the amino acid preceding an Asn-Ser-Thr sequon. Protein Science : A Publication of the Protein Society 2010, 20 (1). DOI: 10.1002/pro.551.

(51) Gao, C.; Stavenhagen, K.; Eckmair, B.; McKitrick, T. R.; Mehta, A. Y.; Matsumoto, Y.; McQuillan, A. M.; Hanes, M. S.; Eris, D.; Baker, K. J.; et al. Differential recognition of oligomannose isomers by glycan-binding proteins involved in innate and adaptive immunity. Science Advances 2021, 7 (24). DOI: 10.1126/sciadv.abf6834.

(52) Fogarty, C. A.; Fadda, E. Oligomannose N-Glycans 3D Architecture and Its Response to the FcγRIIIa Structural Landscape. The Journal of Physical Chemistry B March 4, 2021, 125 (10). DOI: 10.1021/acs.jpcb.1c00304.

(53) FH, T.; A, S.; N, I.; KC, O.; JP, S.; CT, T.; SH, T.; KT, W.; LP, W.; KK, T.; et al. A systematic review on Nipah virus: global molecular epidemiology and medical countermeasures development - PubMed. Virus evolution 2024, 10 (1). DOI: 10.1093/ve/veae048.

(54) Perez-Riverol, Y.; Bandla, C.; Kundu, Deepti J.; Kamatchinathan, S.; Bai, J.; Hewapathirana, S.; John, Nithu S.; Prakash, A.; Walzer, M.; Wang, S.; et al. The PRIDE database at 20 years: 2025 update. Nucleic Acids Res 2025, 53 (D1). DOI: 10.1093/nar/gkae1011.

(55) Masson, G. R.; Burke, J. E.; Ahn, N. G.; Anand, G. S.; Borchers, C.; Brier, S.; Bou-Assaf, G. M.; Engen, J. R.; Englander, S. W.; Faber, J.; et al. Recommendations for performing, interpreting and reporting hydrogen deuterium exchange mass spectrometry (HDX-MS) experiments. Nature Methods 2019 16:7 2019, 16 (7). DOI: 10.1038/s41592-019-0459-y.

