## Supporting File 1. Methods Supplementary Figures and Tables for "An evolutionary distinct Nipah G glycosylation site provides stability for receptor engagement"

**Authors**

Tia É. Hawkins[a,b,c], Valeria Calvaresi[b,d], Sean A. Burnap[b,d], Liang Wu[a,e*], Weston B. Struwe[b,d*]

**Addresses**

a. The Rosalind Franklin Institute, Harwell Science & Innovation Campus, Harwell OX11 0FA, U.K.

b. Kavli Institute for Nanoscience Discovery, University of Oxford, OX1 3QU, U.K.

c. Department of Chemistry, University of Oxford, Oxford OX1 3TA, U.K.

d. Department of Biochemistry, University of Oxford, Oxford, OX1 3QU, U.K.

e. Division of Structural Biology, Nuffield Department of Medicine, University of Oxford, Oxford, OX3 7BN, U.K.

**Correspondents**

*, *

### Methods

#### Phylogenetic analysis of Nipah G isolates

81 Nipah G isolates and the Hendra G RefSeq (AAC83193, NP_047112) were exported as fasta sequences from NCBI Virus database (https://www.ncbi.nlm.nih.gov/labs/virus/vssi/#/, accessed 31/03/2025) (Tables S1-S3). Evolutionary analyses were conducted in MEGA11[1]. The evolutionary history was inferred by using the Maximum Likelihood method and JTT matrix-based model[2]. Initial tree(s) for the heuristic search were obtained automatically by applying Neighbour-Join and BioNJ algorithms to a matrix of pairwise distances estimated using the JTT model, and then selecting the topology with superior log likelihood value. There were a total of 604 positions in the final dataset[1].

#### DNA production and purification

NiVG 71-602, NiVG 183-602, ephrin B2 28-165 and *E.coli* biotin ligase (BirA) constructs in pOPIN vectors for mammalian expression (Table S5). DNA was transformed into DH5a NEB competent cells (New England Bioscience) for purification. 2 µL DNA was added to 50 µL of cells, and rested on ice for 45 mins, before heat shocking (45 seconds 42°C). Cells recovered on ice for a further 2 mins before addition of 400 µL of SOC outgrowth media (New England Bioscience), and incubation at 37°C for 2 hrs. 100 µL of cells were then plated onto LB agar plates containing ampicillin (Sigma), and grown overnight at 37°C. 10 mL LB media (with ampicillin) was the inoculated with a streak of colonies and grown for 8 hrs at 37°C before adding to 1L of LB media (with ampicillin) and incubating at 37°C overnight. The following day, the Qiagen Giga Prep protocol, including isopropanol and ethanol clean up steps, were followed to yield DNA for transfection.

**Site directed mutagenesis**

Polymerase chain reaction (PCR) was used for site directed mutagenisis. Primers were designed so that codon mutations lay in the FWD primer only, generating linear DNA with non-overlapping primers (Table S6). PCR was carried out by the manufacturer's instructions, where Q5 Polymerase, DNTPs, Q5 buffer (New England Bioscience) and primers (Integrated DNA Technologies) were mixed with 25 ng of DNA with $ddH_2O$. PCR mix was then added to KLD reaction buffer, KLD enzyme mix and $ddH_2O$ and incubated for 5 minutes before adding 10 µl of DNA to chemically competent DH5a cells (New England Bioscience). DNA was produced and purified as previously described, and sanger sequencing was using to verify correct site mutagenesis

**Protein Expression and Purification**

Expi293F™(Gibco) cells cultured in Expi293™ Expression Medium (Gibco) were seeded to 1 x$10^6$ cells per mL and placed at 37°C, 0.05% $CO_2$, 24 hours prior to transfection. On the day of transfection, a solution of OptiMem (10% v/v) (Gibco), DNA (1 µg/mL) and PEI 40kDa (5.4 µg/mL) (Polysciences) was added to the culture. 18 hours post transfection, valproic acid (5.9 mM), sodium propionate (6.8 mM) and glucose (50 mM) were added (biotin (840 µM) was added to cultures containing BirA) before incubating for a further 6 days. Conditioned media was harvested (6000 *g,* 30 mins, 4°C) and supernatant loaded onto a Histrap Excel 1 mL (Cytiva). Proteins were washed in PBS, pH7.4, 20 mM imidazole before eluting with 1M imidazole with a gradient elution of 0-100% over 15 minutes. Ephrin B2 purification was supplemented with a high salt wash of 300 mM NaCl prior to elution. Peak fractions were pooled and concentrated in 10 kDa (ephrin B2), 30 kDa (NiVG head constructs) or 50 kDa MWCO PES concentrators (Vivaspin) to 600 µL (4000 g, 4°C). Concentrate was injected onto a Superdex 200 increase 10/300 gl (Cytiva) and ran in PBS pH 7.4 for 1.4 CV at 0.75 mL/min. Desired peak fractions were ran on SDS-PAGE (Invitrogen) before pooling. Pooled peak fractions were concentrated as described above and snap froze in 10 µL aliquots before storing at -80°C.

**Release and uHPLC of N-glycans**

10 µg of 5 biological replicates of Nipah $G_M$ ectodomain were ran on 4-12% Bis-Tris SDS PAGE gel (40 mins 200V) (Invitrogen). Gel bands of interest were extracted and de-stained in

50 mM ammonium bicarbonate (AmBic) 50% acetonitrile (MeCN). Incubated at 37 °C for 30 mins and washed in 50 mM AmBic 50% MeCN. Dehydrated gel bands in 100% MeCN then covered gel with 700 µL containing PNGaseF at 12.5 ng/µL (in house) and incubated at 4 °C for 1 hr, before leaving for 18 hrs 37 °C.

Reaction mix and three washes of gel bands with 100 µL of HPLC grade water (incubation 10 mins 37 °C) was pooled. Glycans were dried and resuspended in 30 µL of HPLC grade water. 80 µL of freshly prepared 2-AA reaction mix (30 mg/mL 2-anthracillic acid, 45 mg/mL $NaBH_3CN$, 4% w/v $CH_3COONa{\cdot}3H_2O$, 2% w/v $H_3BO_3$; in MeOH) was added to glycans and incubated at 80 °C for 1hr. Samples were cooled before clean-up with Spe-ed amide 2 cartridges. Labelled glycans were eluted with two washes with 750 µL HPLC grade water and lyophilised for 18 hrs.

Labelled glycans were resuspended in 20 µL HPLC grade water and treated 10 µL with endoglycosidase: 500 NEB units of endoglycosidase H (Endo-H) (New England Bioscience) was added to 10 µL of labelled glycans and supplemented with GlycoBuffer 3 (New England Bioscience) and incubated for 22 hrs 37 °C. Endo-H reaction mix was cleaned up with 10 kDa MWCO cartridges (Omega), before drying and resuspending in HPLC grade water.

MeCN was added to labelled glycans to a final concentration of 70% MeCN. 5 µL of glycans were injected onto an ACQUITY UHPLC BEH Amide column (1.7 µM, 2.1x150 mm) using an Agilent Technologies UHPLC (Table S7). Dextran, $Man_5$ and $Man_9$ glycan standards (Ludger) were ran beforehand as controls.

**Release and IM-MS/MS of N-glycans**

40 µg of 3 biological replicates of Nipah G were ran on 4-12% Bis-Tris SDS PAGE gel (40 mins 200V) (Invitrogen). Briefly, gel was stained with coomassie before extracting gel bands of interest. Destained gel pieces in 50 mM AmBic 50% MeCN, incubating at 37 ºC for 30 mins, before washing in 50 mM AmBic 50% MeCN. Dehydrated the gel pieces in 100% MeCN then covered gel with 700 µL containing PNGaseF (in house) in a 1:10 ratio of enzyme to glycans, incubated at 4 °C for 1 hr, before leaving for 18 hrs 37 °C. Released N-glycans were desalted using C18 zip tips (Merck) overlayed with porous graphite carbon (in house), then dried down and resuspended in 50:50 MeOH:$H_2O$.

Approximately 2 µL of N-glycan sample material was directly ionized into a Synapt XS (Waters) by nano-electrospray ionization (nano-ESI) (Table S8), using gold-coated borosilicate glass capillaries (prepared in-house). Data acquisition and processing were carried out using Waters Driftscope (version 2.8) software and MassLynx™ (version 4.1) (Tables S9-11).

**Glycoproteomics**

3 µg of 6 biological replicates of Nipah $G_M$ ectodomain were ran on 4-12% Bis-Tris SDS PAGE gel (40 mins 200V) (Invitrogen). Gel bands of interest were extracted and de-stained in 50 mM AmBic 50% MeCN and incubated at 37 °C for 30 mins, before washing in 50 mM AmBic 50% MeCN. Dehydrated gel bands in 100% MeCN then reduced proteins with 10 mM dithiothreitol in 100 mM AmBic and incubated for 30 min at 56°C. Dehydrated gel bands in 80% MeCN for 10 mins at room temperature (RTP). Replaced solution with 50 mM iodoacetamide in 100 mM AmBic and incubated in the dark for 30 mins. Removed liquid and washed gel bands sequentially with 100% MeCN and 50 mM AmBic. Dehydrated gel pieces 100% MeCN then added 12.5 ng/µL protease (trypsin (Promega), chymotrypsin (Promega) or alpha-lytic (New England Bioscience)) in 50 mM AmBic, before incubating at 37°C for 1 hr. Solution was replaced with fresh 50 mM AmBic, incubating bands at 37°C 18 hrs. Pooled peptide solution and sequential washes of 5% (v/v) formic acid (FA) in $H_2O$ and MeCN. Samples were dried and reconstituted in 2% MeCN, 0.05% trifluoroacetic acid and analysed by LC-MS/MS.

Samples were analysed using an Ultimate 3000 UHPLC coupled to an Orbitrap Q Exactive mass spectrometer (Thermo Fisher Scientific). Peptides were loaded onto a 75 µm × 2 cm pre-column and separated on a 75 µm × 15 cm Pepmap C18 analytical column (Thermo Fisher Scientific). Buffer A was 0.1% FA in $H_2O$ and buffer B was 0.1% FA in 80% MeCN with 20% $H_2O$. A 100-minute linear gradient (Table S12) and universal HCD identification method (Table S13) method was used.

Data were analysed in Byos (Protein Metrics). Digestion was set to RK, FYWML and TASV for trypsin, chymotrypsin and alpha-lytic protease digests respectively with maximum of two missed cleavages. Fixed modifications were carbamidomethylation (57.02 Da). And variable modifications were methionine oxidation (15.99 Da) and deamidation (N or Q) (0.98 Da). Byos in-built common human N-linked (132 glycans) and O-linked glycan (6 glycans) glycan libraries were used. FDR was set to 0.01. For analysis, a minimum Byos threshold score of 300 was used for glycopeptide filtering, with the exception of peptides covering site N72 for which a score of 100 was used. Glycopeptides were manually validated, and one peptide sequence per glycosite was chosen as representative for that site (Table S4). For quantification, the extracted ion chromatogram intensities for each glycopeptide of that sequence were summed and plotted relative to the total intensity for each glycosite.

**Differential Endoglycosidase treatment with $^{18}O$ labelling**

The method by Cao *et al*[3] was adapted with the following modifications; 5 μg of Nipah $G_M$ ectodomain was denatured in 4M Urea, 1% SDS for 1hr at RTP. Protein was reduced in 10 mM TCEP (pH 7.5) for 30 mins RTP and labelled with 50 mM iodoacetamide in 100 mM AmBic for 30 mins in the dark, RTP. Protein solution was added to 250 μg of 50:50 mix of Speed Bead Magnetic Carboxylate Modified Particles (Cytiva: 65152105050250 and 45152105050250). MeCN was added to a final concentration of ~90% and incubated for 30 mins, 37C, 1000 rpm. A magnetic plate was used to aspirate the supernatant before and during sequential washes of 80% EtOH (x2) and 100% MeCN. Replaced washes with 0.2 μg of trypsin/LysC (Promega) in 50 mM $(NH_4)HCO_3$ in $H_2O$ and incubated overnight at 37 °C. Peptide solution was kept and 5% DMSO $H_2O$ was added to beads for 2 mins, before pooling solution with peptides. Half of the peptides were further treated with 0.2 μg of GluC (NEB) in 50 mM AmBic for 5.5 hrs at 37 °C, after which all peptides were dried.

Peptides were resuspended in 50 mM ammonium acetate (pH 5.5) (1000 rpm, 10 mins, 37 °C) before addition of 313 NEB units of Endo-H (New England Bioscience) per sample. Samples were incubated for 1 hr at 37 °C before drying and sealing. PNGase F (New England Bioscience) was lyophilised and resuspended in 50 mM AmBic in $H_2{}^{18}O$ before adding to peptides, with a 160 NEB units per sample. Samples were incubated for 30 mins 37 °C before heat inactivation (10 min, 100 °C) and drying. Labelled peptides were reconstituted in 2% MeCN 0.05% TFA and analysed by LC-MS/MS using the same method as for glycoproteomics.

Data were analysed in Proteome Discoverer (Thermo Fisher Scientific). Digestion was set to RK, and RKED for trypsin or trypsin/gluC protease digests respectively with maximum two missed cleavages. Static modifications were carbamidomethylation (57.02 Da) and dynamic modifications were methionine oxidation (15.99 Da) and $^{18}O$ deamidation (N only) (2.98 Da). Strict FDR was set to 0.01. The library searched consisted of the Nipah $G_M$ sequence lacking a signal sequence, supplemented with the CRAPome[4]. The same peptide sequences were used for quantification (Table S4). For quantification, the extracted ion chromatogram intensities for each peptide of that sequence were summed and plotted relative to the total intensity for each glycosite.

**Biolayer Interferometry of NiVG glycan mutants**

Octet® streptavidin biosensors (Sartorius) were soaked in 200 μL PBS for 30 mins prior to runs. NiVG mutants and ephrin B2 were diluted in PBS with 0.01% Tween 20. Blocking buffer was PBS with 0.01% Tween. Runs were carried out on an Octet® R8 system (Table S14).

Concentrations of mutants were titrated in a 2-fold dilution series starting at 200, 100 or 44 nM. Timings and concentrations were determined prior such that the association phase would reach steady state and the dilution series covered both above and below the determined $K_D$. Analysis was carried out in Octet® Analysis Software, where baselines were subtracted prior to fitting data to a 1:1 binding model. The $R_{max}$ of each curve was fit to a steady state model to give the $K_D$ values. Students unpaired, two-tailed t-test was used to calculate statistical significance for $K_D$ values.

**HDX-MS analysis of WT, T483A, N481D and WT-EB2 complex**

NiVG WT head, T483A and N481D were diluted to 5.9 µM, and deuterated PBS (pD 7.4) was added to a deuterium fraction of 88% to initiate the exchange reaction. At selected time intervals (15 s, 150 s , 1500 s, 15000 s, 24 hrs), a fraction of reaction mixture containing the equivalent of 23 pmol of protein sample was withdrawn and quenched with an ice-cold phosphate buffer containing 2M urea and 40 mM TCEP, which dropped the pH/D to 2.3 and the deuterium fraction to 50%. Quenched samples were kept for 30 s on ice then flash frozen in LN2, and stored at -80 °C before LC-MS analysis. For WT-EB2 complexing, NiVG WT head was incubated in a 1:1.2 molar ratio with ephrin B2 (28-165), or an equivalent volume of PBS. Deuterium labelling was carried out as specified above, and the equivalent of 46 pmol of protein samples were quenched. Maximally labelled samples (MaxD) were produced by labelling NiVG WT head, T483A and N481D at 37 °C in the presence of 4 M deuterated Urea in $D_2O$ and 2.4 mM TCEP, keeping the deuterium fraction at 88%. The maximally labelled samples were quenched after 18 hrs with ice-cold phosphate buffer (1.4 % FA) in a similar manner as for the other labelled samples and stored in the same fashion. Timepoints were repeated in triplicate, but duplicates were recorded for WT (mutant analysis) 15 s, WT (ephrin B2 analysis) 1500 s, N481D 150 s, T483A 150 s.

For LC-MS analysis, samples were quickly thawed and injected into an Acquity UPLC M-Class System with HDX Technology (Waters), on-line digested at 20 °C into a hand packed pepsin column, prior to online deglycosylation with a PNGAse Rc column[5] (Affipro), which removed N-glycans through enzymatic deamidation of N-glycosylated asparagines into aspartates. Peptides were trapped/desalted with solvent A (0.23% formic acid in water) for 4 min at 120 µL/min and at 1 °C through an Acquity BEH C18 VanGuard pre-column (1.7 µm, 2.1 mm × 5 mm, Waters). Peptides were eluted from the trap column into an Acquity UPLC BEH C18 analytical column (1.7 µm, 1 mm × 100 mm, Waters) with a 7 min-linear gradient raising from 8 to 35% of solvent B (0.23% formic acid in acetonitrile) at a flow rate of 40 µL/min and at 1 °C (Table S15). Eluted peptides went through electrospray ionization in positive mode and underwent MS analysis onto a Synapt-G2 Si, TWIMS-TOF (Waters) with ion mobility separation enabled (Table S16). For peptide identification, non-deuterated protein samples were ran with the same method as for deuterated samples, but underwent fragmentation using an $MS^E$ method, applying collision energy ramping from 20 to 30 kV (Table S17)

$MS^E$ runs were analysed with ProteinLynx Global Server (PLGS) 3.0 (Waters) and peptides identified in 3 out of 4 replicates, with at least 0.2 fragments per amino acid (at least 2 fragments in total), with mass error below 10 ppm were selected in DynamX 3.0 (Waters). Using both retention time and drift time, spectra of deuterated peptides were matched to the peptide map, and manually annotated to calculate peptide deuterium incorporation.

#### Statistics and Graphing

Students unpaired, two-tailed t-test was used to test for statistical significance between complex glycan containing glycopeptides and $^{18}O$ labelled peptides. Statistics and Graphing were conducted in GraphPad Prism v10.2.3.

Students unpaired, two-tailed t-test was used to test for statistical significance between the $K_D$ of Nipah $G_M$ Head and glycosite mutants, and between Nipah $G_M$ head ± kifunensine. Statistics and Graphing were conducted in GraphPad Prism v10.2.3.

The threshold for a statistically significant difference in HDX between two states was established based on an approach described earlier[6]. Briefly, a confidence interval (CI) was calculated based on the average standard deviation (SD) of peptide deuterium uptake values for time points performed in triplicates, according to equation 1:

Equation 1. $$SD_{state} = \frac{\sum SD_i{}^2}{N}$$

where N is the number of peptides considered, multiplied by the number of time points performed in triplicate. The CI at the significance level of 99% between two states was calculated based on a zero-centered average difference in deuterium content, considering a two-tailed distribution with two degrees of freedom (n = 3) (equation 2)

Equation 2. $$CI = 9.925 \times \frac{\sqrt{SD_{stateA}+SD_{stateB}}}{\sqrt{n}}$$

The deuterium uptake of peptides carrying T483A were compared to the correspondent WT peptides by normalization with their respective MaxD uptake values (equation 3)[7]:

Equation 3. $$\Delta HDX = D_{max}\ WT\ peptide \left(\frac{Uptake\ T483A\ peptide}{Uptake\ T483A\ D_{max}\ peptide}\right) - Uptake\ WT\ peptide$$

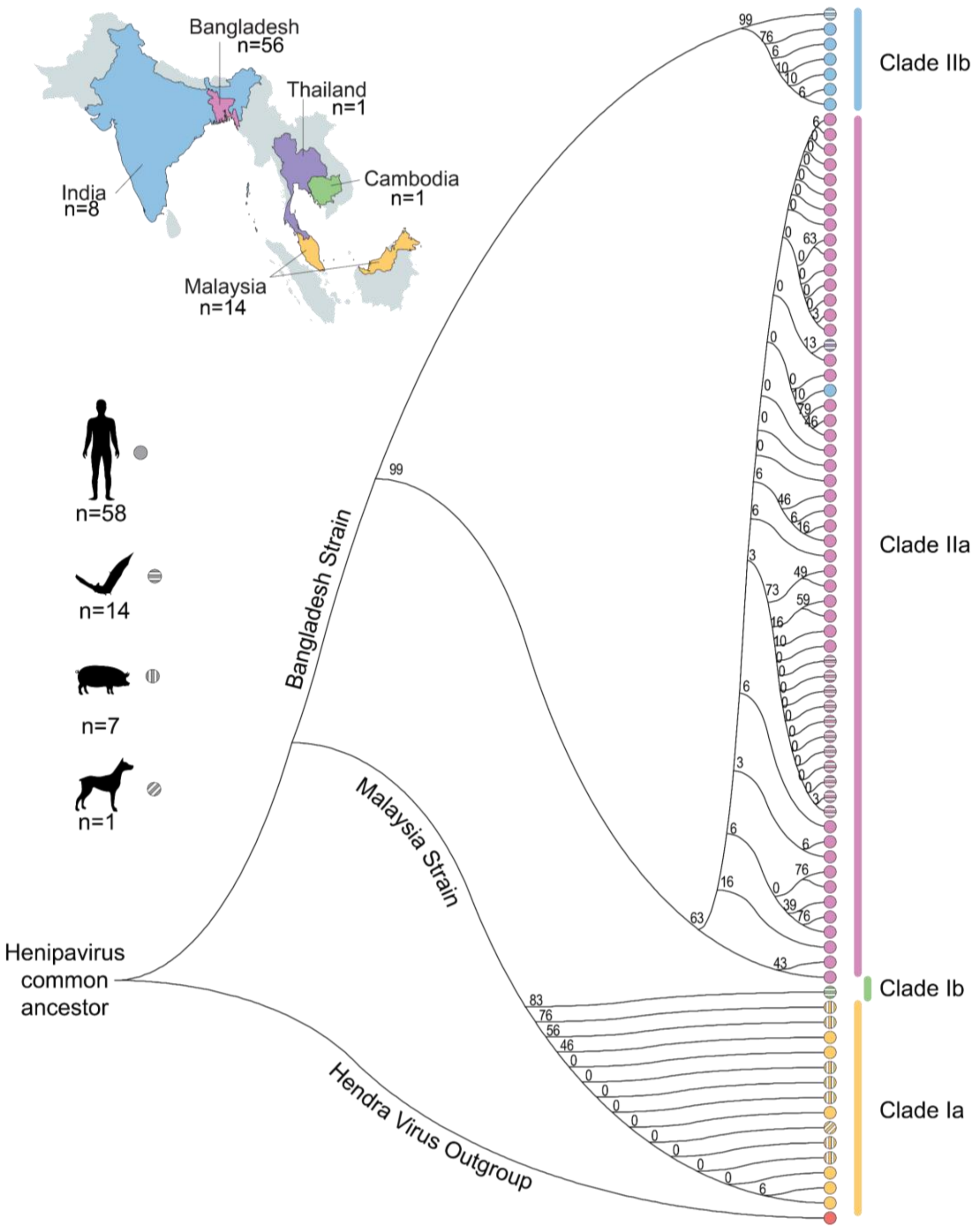


**Figure S1. Evolution of Nipah G from 1998-2023.** Top left: Map of locations and number of isolates per country (with countries colour-coded in the phylogenetic tree below). Middle left: Isolates identified for each host. Right: Maximum likelihood tree of 81 complete Nipah G sequences: The percentage of trees in which the associated taxa clustered together is shown next to the branches. The Hendra virus G protein is shown in orange.

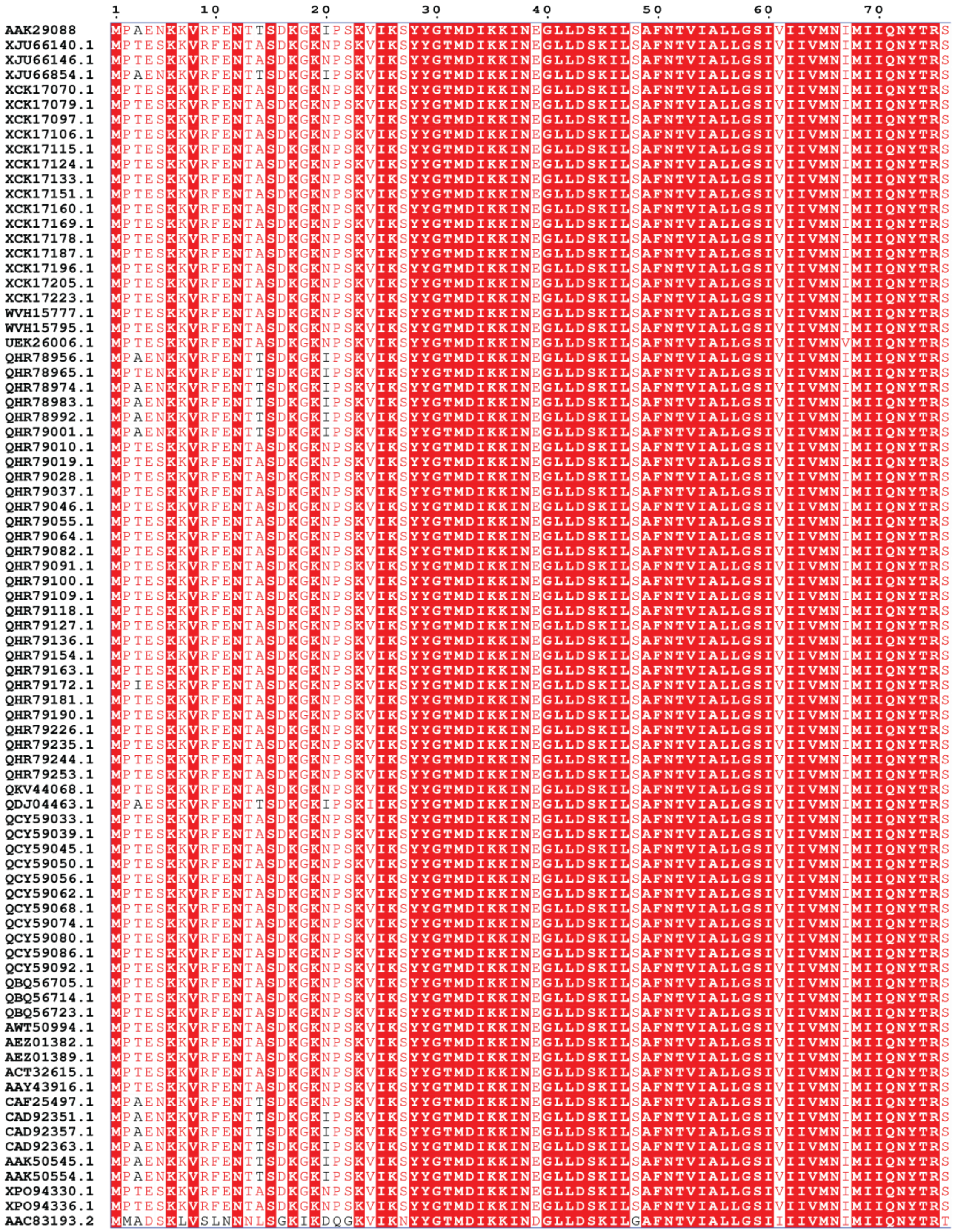

**Figure S2. Multiple Sequence Alignment of 81 Nipah G isolates and 1 Hendra G reference sequence.** Sequences were aligned in MEGA11 and displayed using ESPrit 3.0 (https://espript.ibcp.fr/ESPript/cgi-bin/ESPript.cgi). Conserved amino acids are coloured white on a red background, similar amino acids coloured red, non-conserved in black, and gaps are denoted with dots. Information about sequence accessions can be found in tables S1-S3.

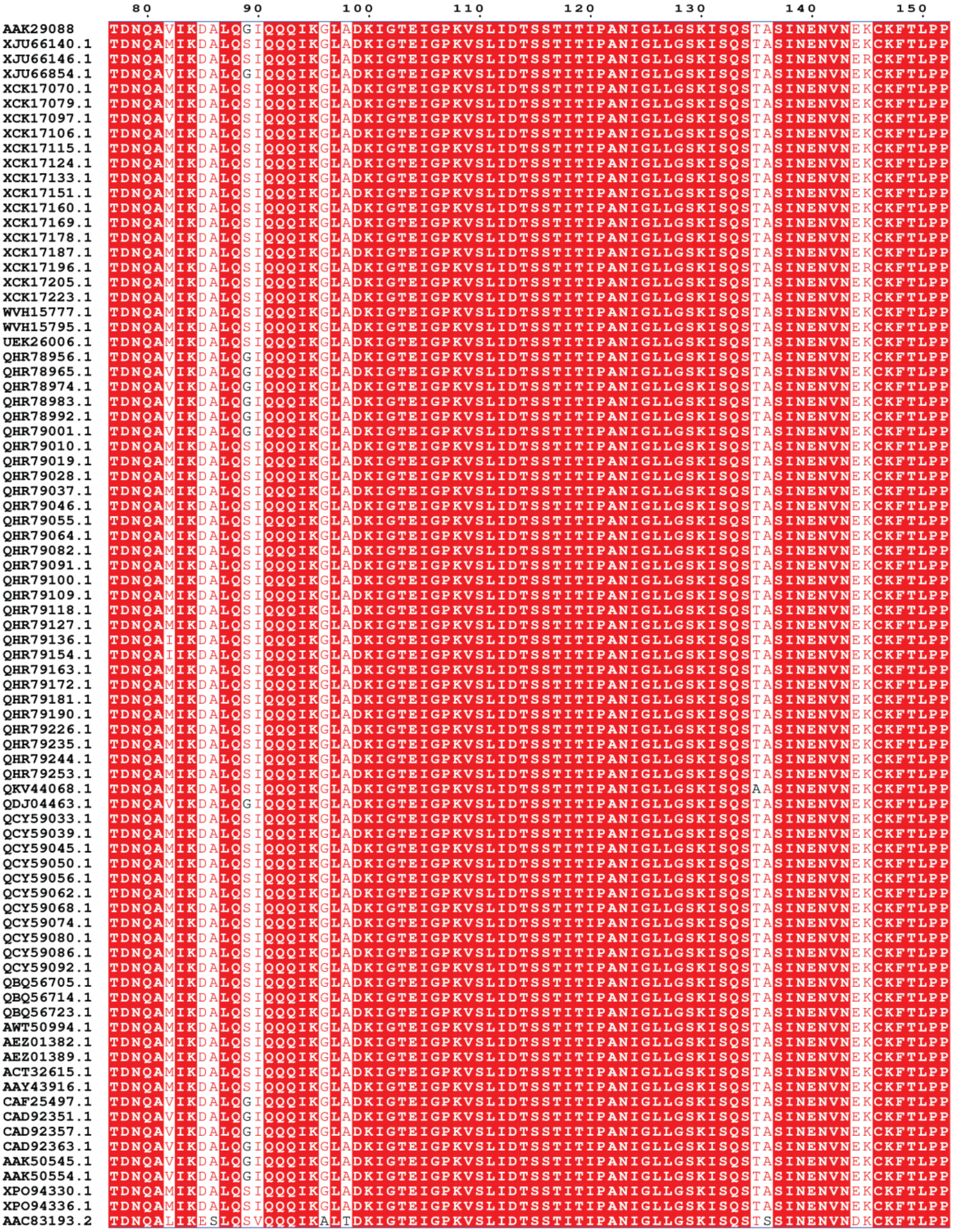


**Figure S2 continued. Multiple Sequence Alignment of 81 Nipah G isolates and 1 Hendra G reference sequence.** Sequences were aligned in MEGA11 and displayed using ESPrit 3.0 (https://espript.ibcp.fr/ESPript/cgi-bin/ESPript.cgi). Conserved amino acids are coloured white on a red background, similar amino acids coloured red, non-conserved in black, and gaps are denoted with dots. Information about sequence accessions can be found in tables S1-S3.

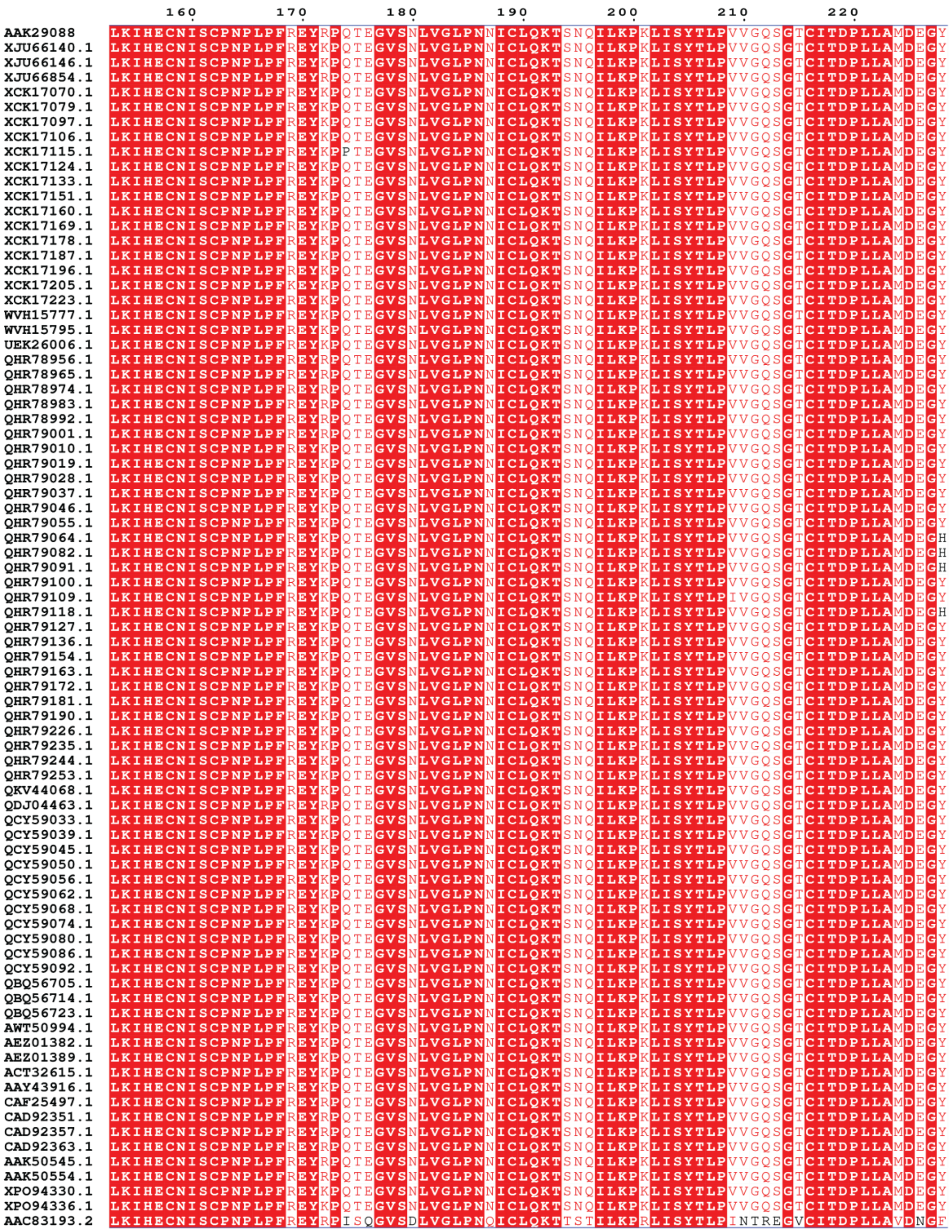


**Figure S2 continued. Multiple Sequence Alignment of 81 Nipah G isolates and 1 Hendra G reference sequence.** Sequences were aligned in MEGA11 and displayed using ESPrit 3.0 (https://espript.ibcp.fr/ESPript/cgi-bin/ESPript.cgi). Conserved amino acids are coloured white on a red background, similar amino acids coloured red, non-conserved in black, and gaps are denoted with dots. Information about sequence accessions can be found in tables S1-S3.

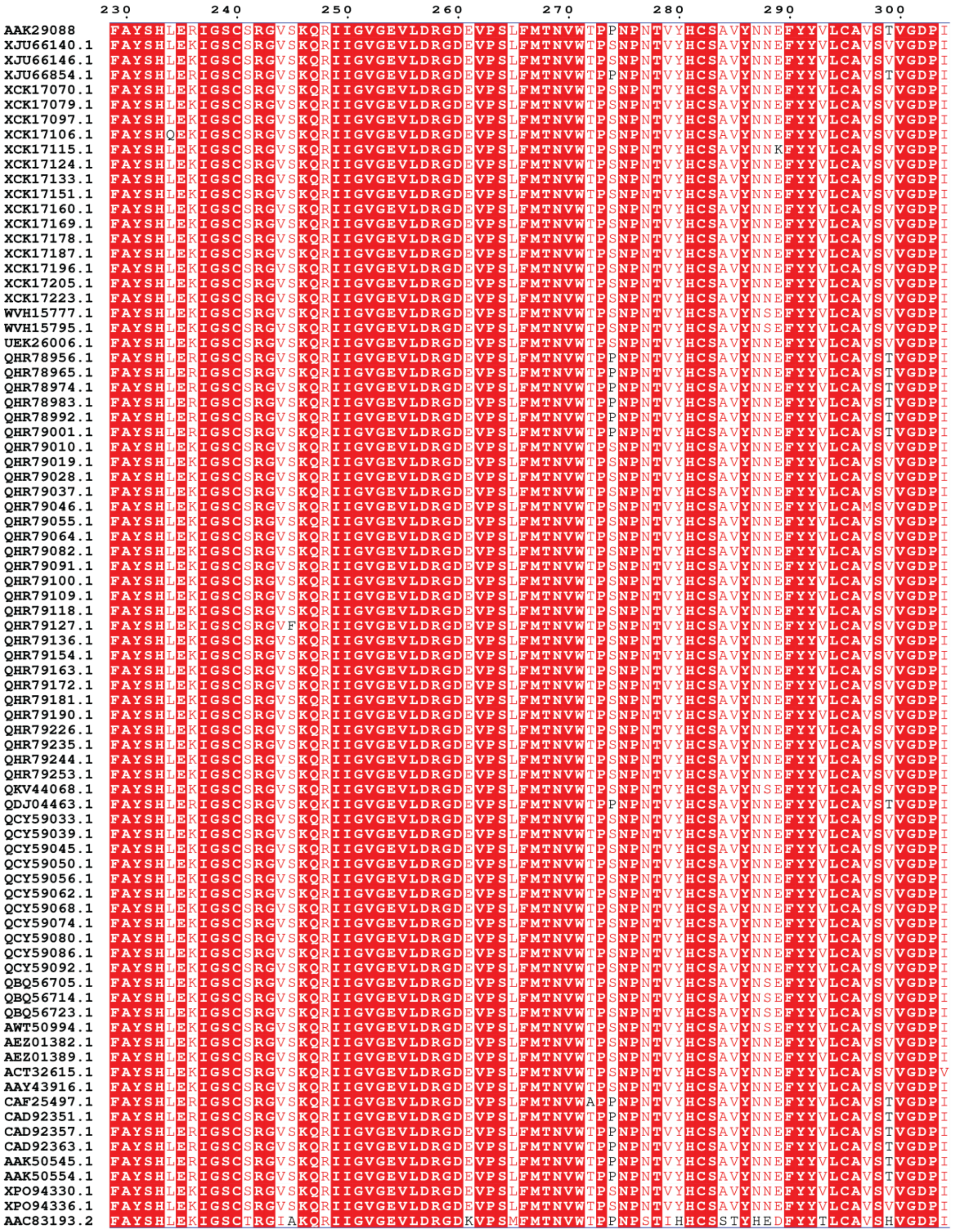


**Figure S2 continued. Multiple Sequence Alignment of 81 Nipah G isolates and 1 Hendra G reference sequence.** Sequences were aligned in MEGA11 and displayed using ESPrit 3.0 (https://espript.ibcp.fr/ESPript/cgi-bin/ESPript.cgi). Conserved amino acids are coloured white on a red background, similar amino acids coloured red, non-conserved in black, and gaps are denoted with dots. Information about sequence accessions can be found in tables S1-S3.

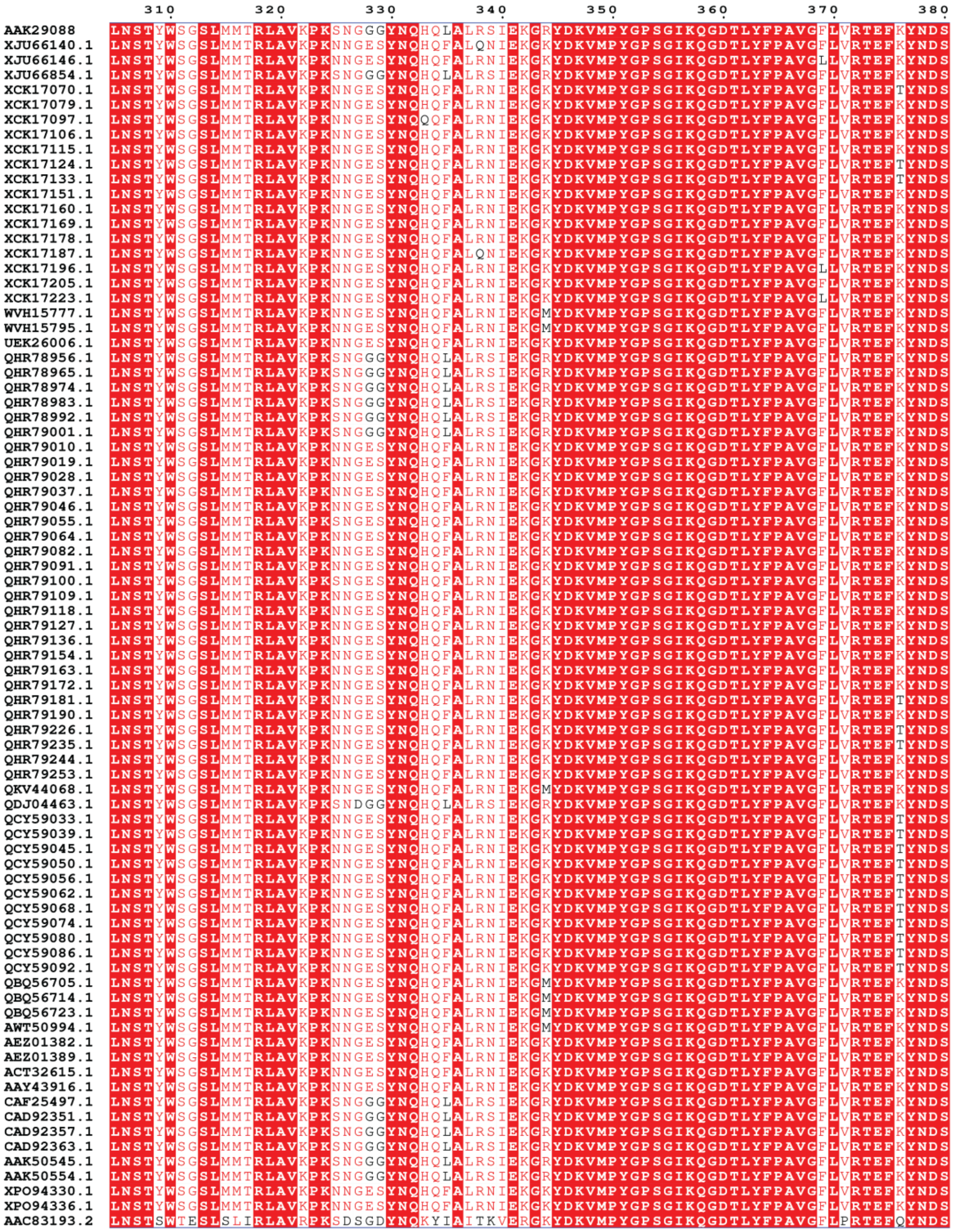


**Figure S2 continued. Multiple Sequence Alignment of 81 Nipah G isolates and 1 Hendra G reference sequence.** Sequences were aligned in MEGA11 and displayed using ESPrit 3.0 (https://espript.ibcp.fr/ESPript/cgi-bin/ESPript.cgi). Conserved amino acids are coloured white on a red background, similar amino acids coloured red, non-conserved in black, and gaps are denoted with dots. Information about sequence accessions can be found in tables S1-S3.

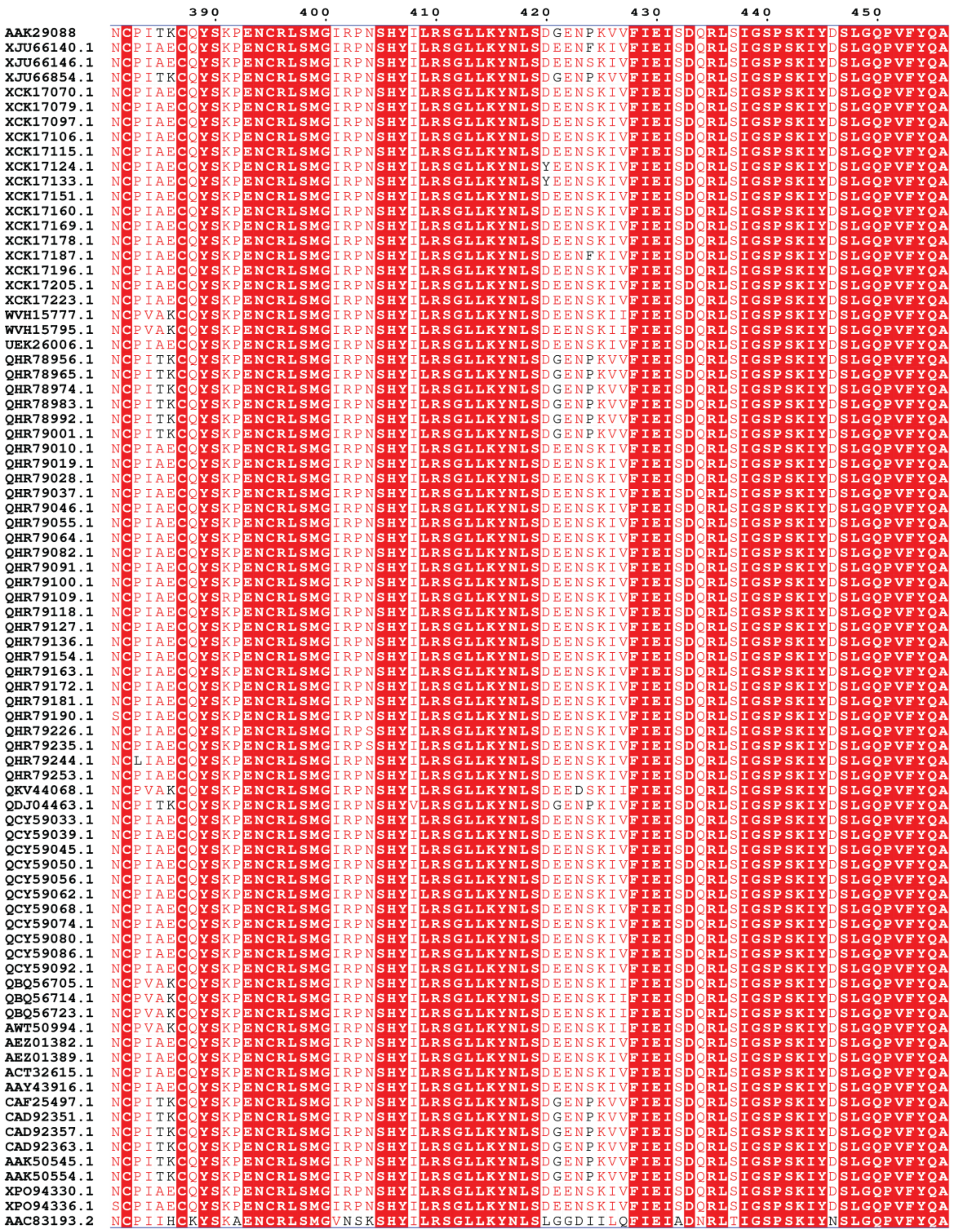


**Figure S2 continued. Multiple Sequence Alignment of 81 Nipah G isolates and 1 Hendra G reference sequence.** Sequences were aligned in MEGA11 and displayed using ESPrit 3.0 (https://espript.ibcp.fr/ESPript/cgi-bin/ESPript.cgi). Conserved amino acids are coloured white on a red background, similar amino acids coloured red, non-conserved in black, and gaps are denoted with dots. Information about sequence accessions can be found in tables S1-S3.

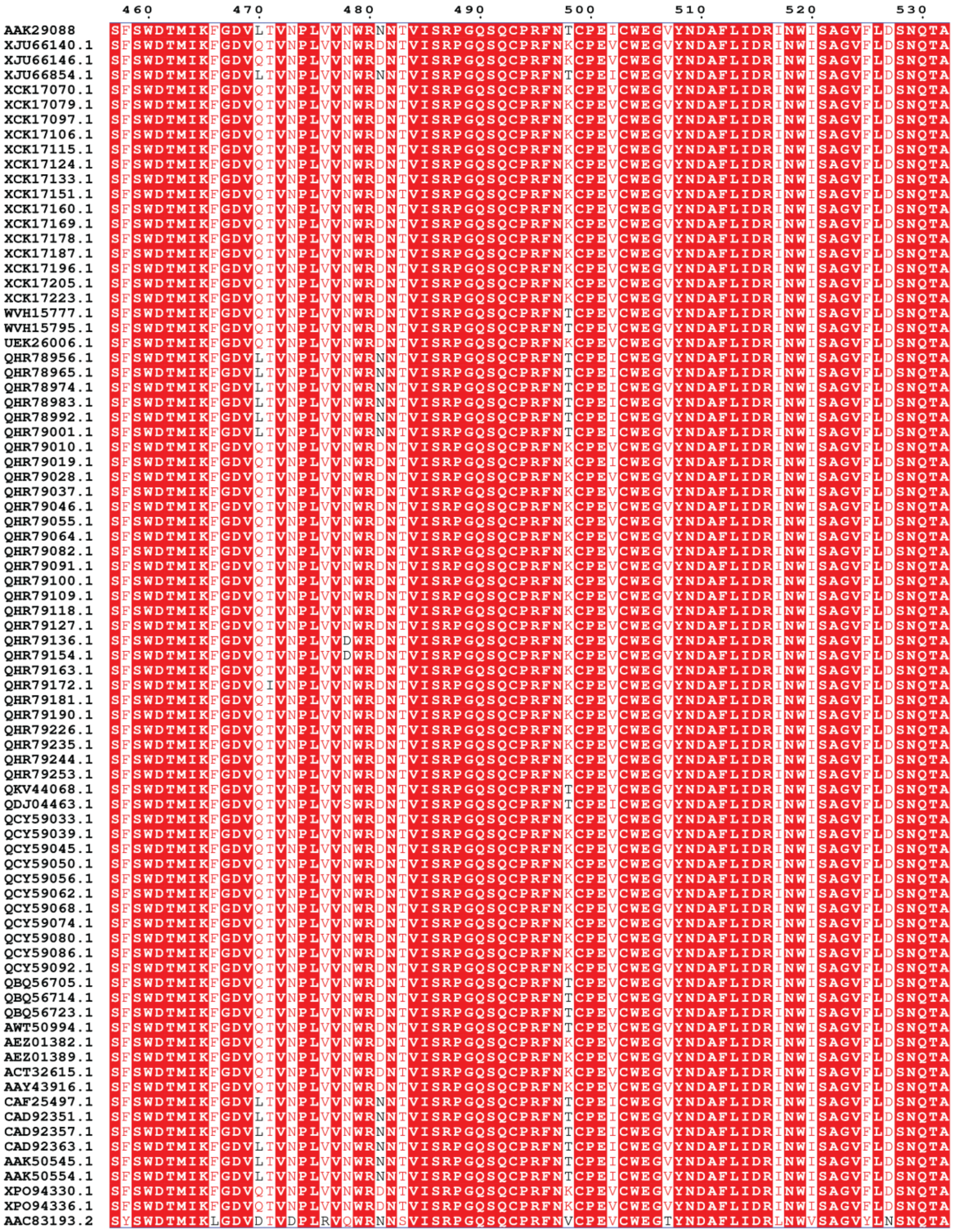


**Figure S2 continued. Multiple Sequence Alignment of 81 Nipah G isolates and 1 Hendra G reference sequence.** Sequences were aligned in MEGA11 and displayed using ESPrit 3.0 (https://espript.ibcp.fr/ESPript/cgi-bin/ESPript.cgi). Conserved amino acids are coloured white on a red background, similar amino acids coloured red, non-conserved in black, and gaps are denoted with dots. Information about sequence accessions can be found in tables S1-S3.

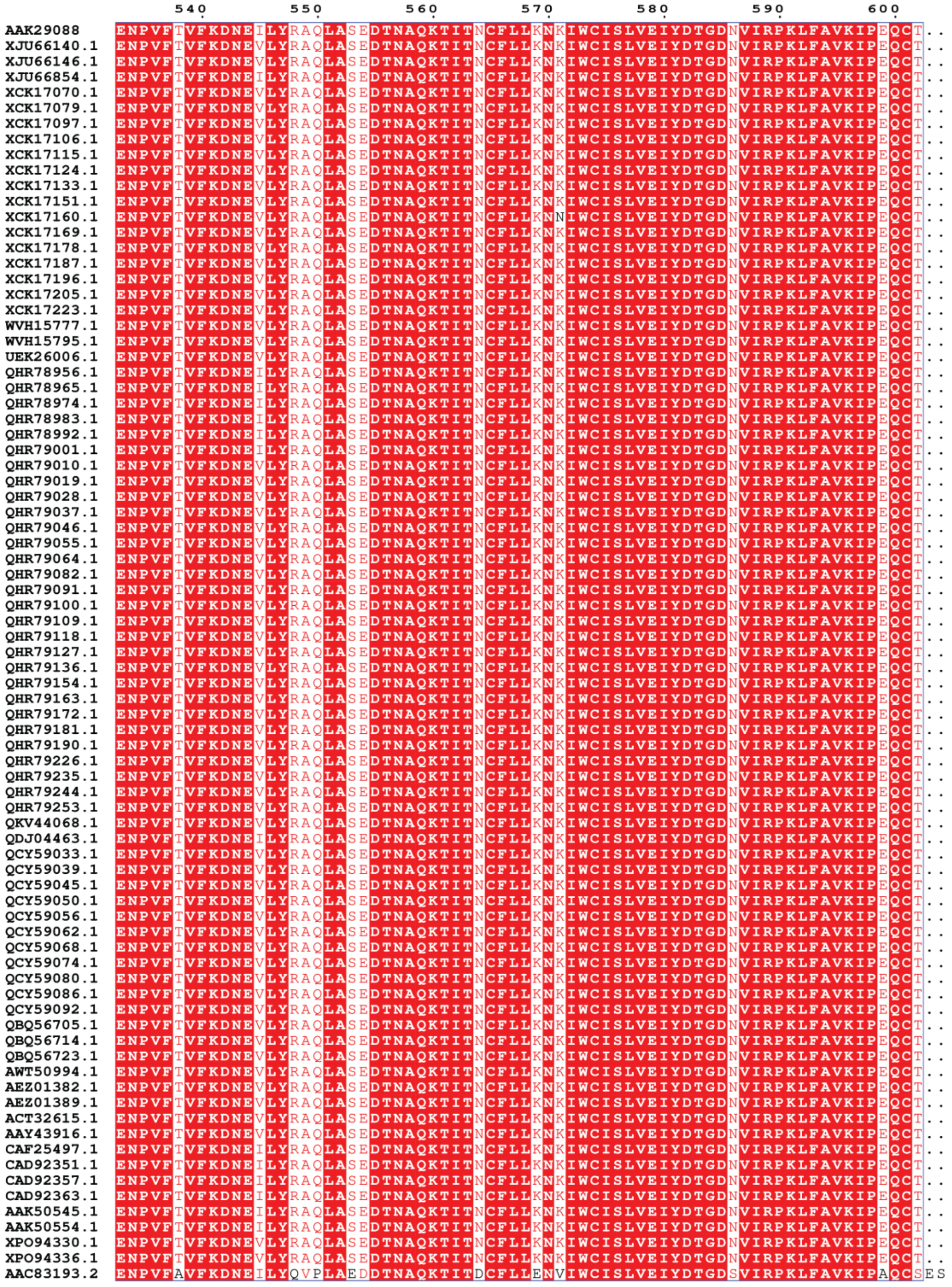


**Figure S2 continued. Multiple Sequence Alignment of 81 Nipah G isolates and 1 Hendra G reference sequence.** Sequences were aligned in MEGA11 and displayed using ESPrit 3.0 (https://espript.ibcp.fr/ESPript/cgi-bin/ESPript.cgi). Conserved amino acids are coloured white on a red background, similar amino acids coloured red, non-conserved in black, and gaps are denoted with dots. Information about sequence accessions can be found in tables S1-S3.

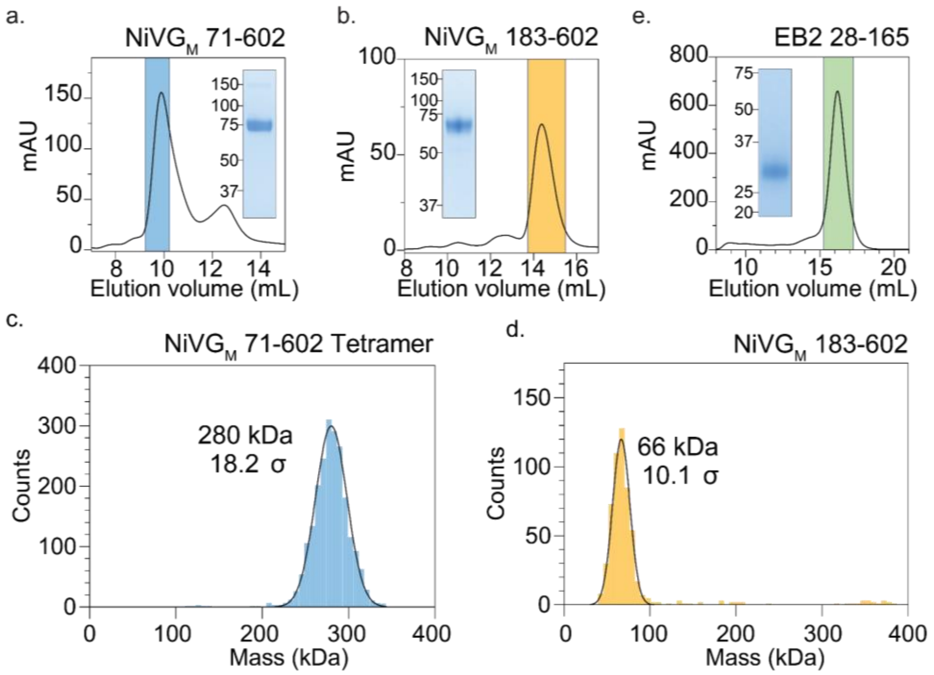


**Figure S3. Purification and oligomeric distribution of Nipah $G_M$ ectodomain and ephrin B2 RBD.** Size exclusion chromatogram of Nipah $G_M$ ectodomain (a) (pooled fractions in blue) and Nipah $G_M$ head domain (b) (pooled fractions in orange), with representative bands from Coomassie stained SDS-PAGE gels. c) Mass photometry of Nipah $G_M$ ectodomain. The theoretical monomeric mass is 60.5 kDa, whilst the theoretical tetrameric mass is 242 kDa. The measured mass was 280 kDa, indicative of a glycosylated tetramer. d) Mass photometry of Nipah $G_M$ head domain. The theoretical monomeric mass is 51 kDa, and the measured mass is 66 kDa, indicative of a glycosylated monomer. e) Size exclusion chromatogram of ephrin B2 RBD (pooled fractions in greens) with representative band from Coomassie stained SDS-PAGE gel. Ephrin B2 RBD runs considerably higher than its theoretical mass of 18.8 kDa (band at ~30 kDa under denatured conditions) due to glycosylation.

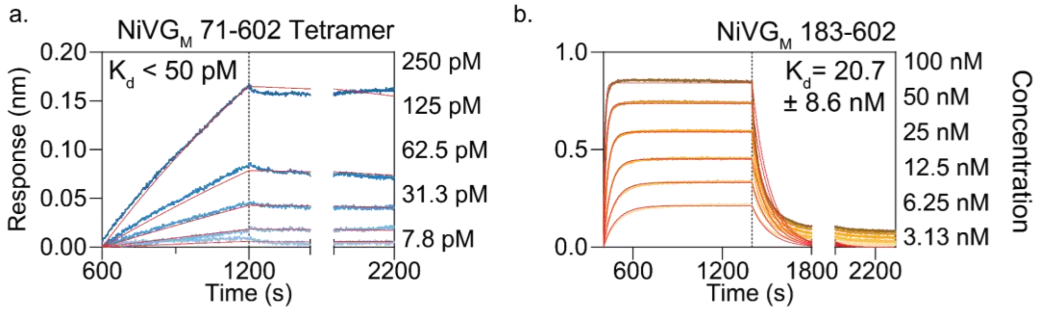


**Figure S4. Biolayer interferometry of Nipah $G_M$ to ephrin B2 RBD.** Nipah $G_M$ ectodomain (a) and Nipah $G_M$ head domain (b) were incubated with immobilised ephrin B2 RBD. Fits are shown in red and used to calculate upper $K_d$ limit for (a). The $K_d$ for (b) was calculated using steady state approximations. Nipah $G_M$ concentration shown to the right of the sensorgrams.

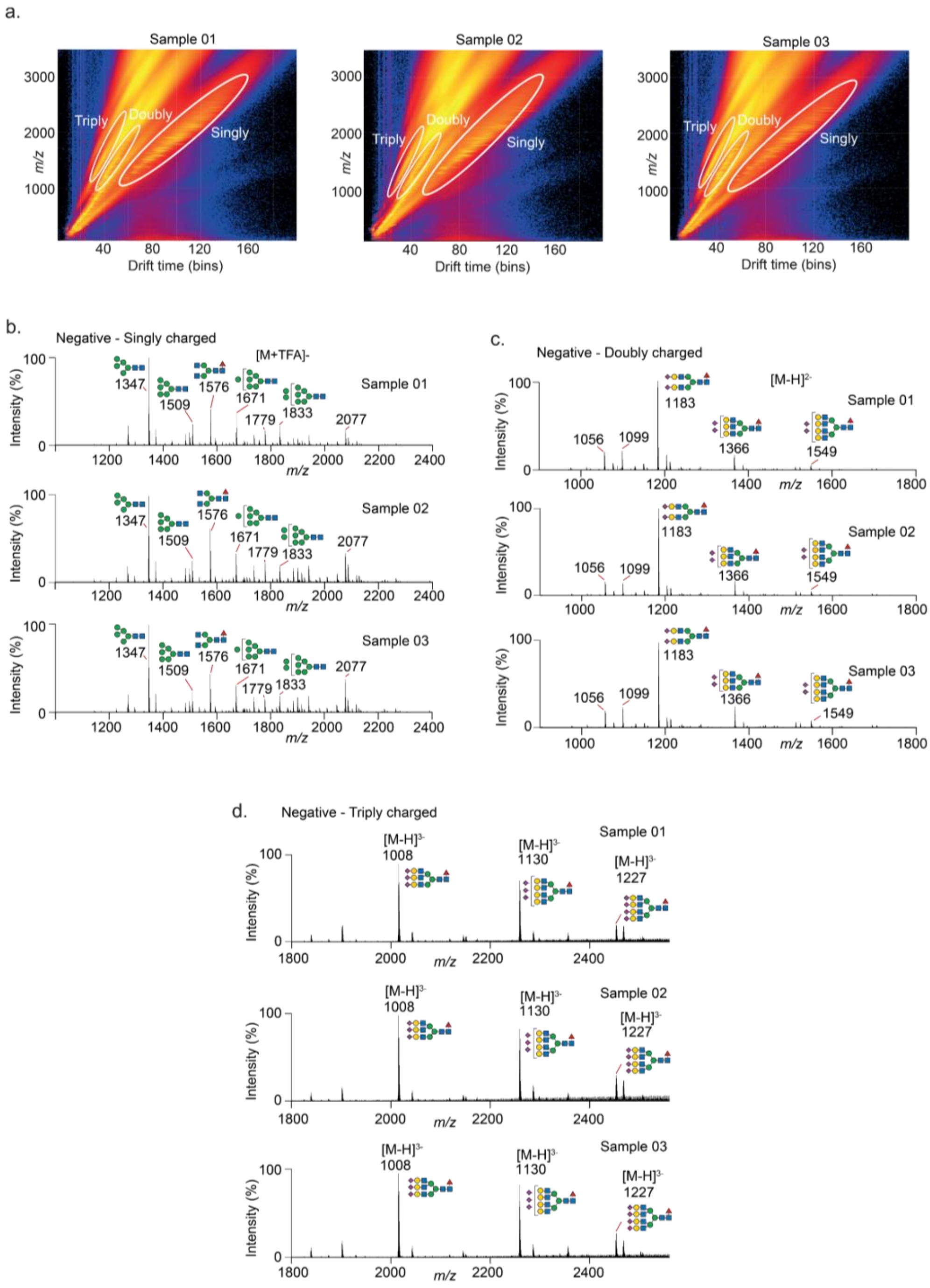


**Figure S5. IM-MS of released N-glycans from Nipah $G_M$ tetramer.** a) Ion mobility drift plots for three biological replicates, with IM-extracted charge states indicated in white circles with their associated MS spectra for b) singly charged, c) doubly charged and d) triply negatively charged species. Three biological replicates are shown. Ions were detected as either deprotonated ions ($[M-H]^-$) or TFA adduct ions.

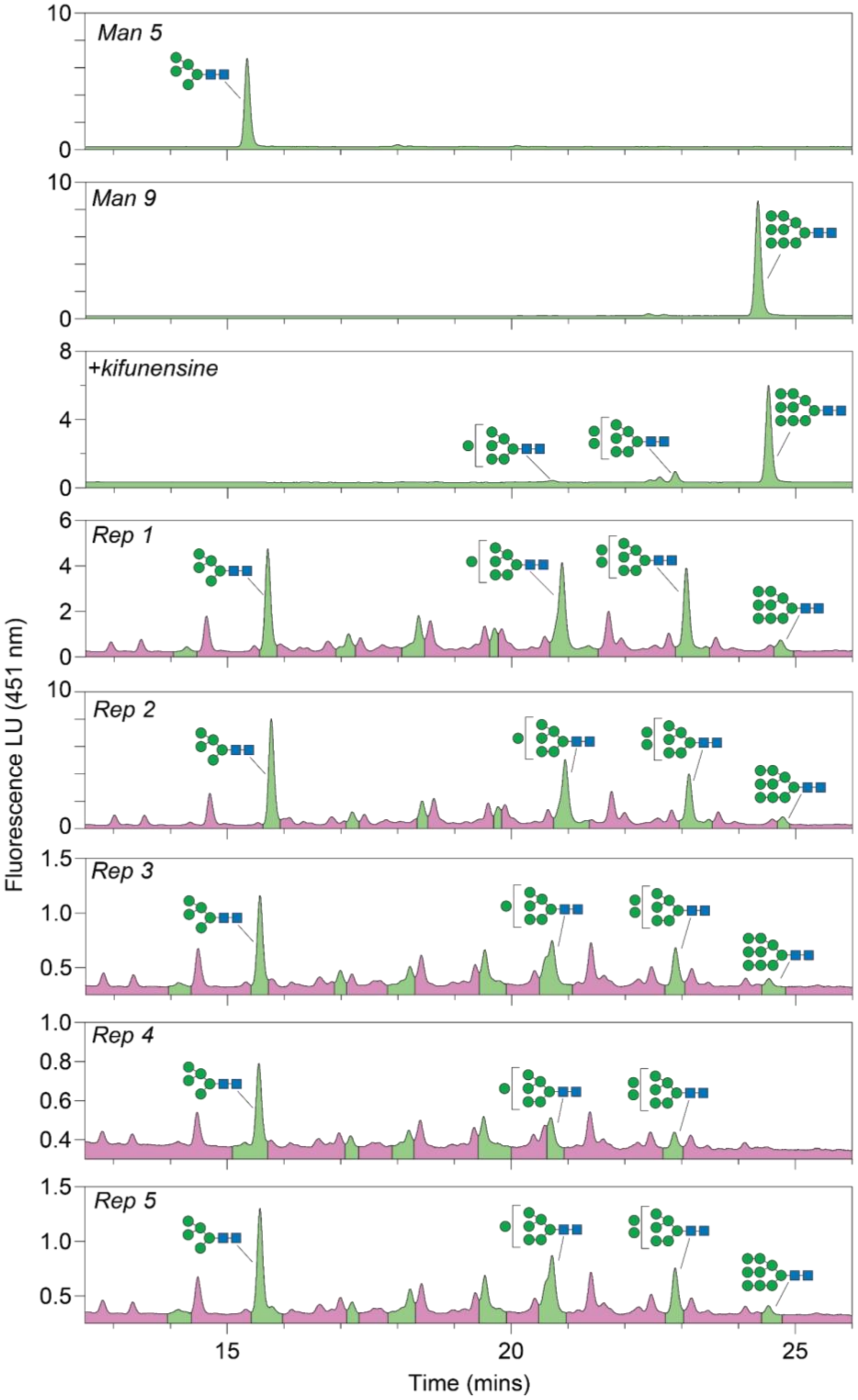


**Figure S6. Endo-H resistant and Endo-H sensitive glycan identification with UHPLC of 2-AA labelled N-glycans.** N-glycans that were endo-H sensitive are marked in green, those which were resistant in purple. The top two chromatograms are a $GlcNAc_2Man_5$ (Man5) and a $GlcNAc_2Man_9$ (Man9) standard. The profile from the top are N-glycans from a NiVG 71 tetramer expressed the presence of kifunensine producing $GlcNAc_2Man_{7\text{-}9}$ N-glycans. These chromatograms were used to identify the $GlcNAc_2Man_{5,7\text{-}9}$ species.

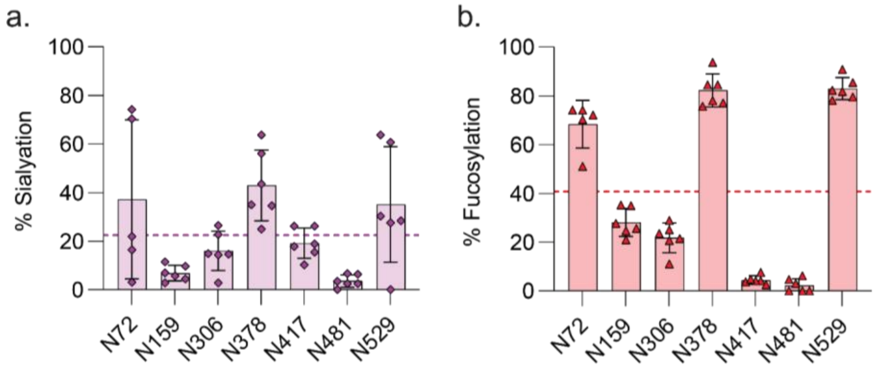


**Figure S7. Site-specific sialyation and fucosylation of Nipah $G_M$ tetramer. $G_M$.** Percentage of glycopeptide species containing sialic acid (a) and fucose (b) at each site for Nipah G. Dotted line represents the mean sialyation or fucosylation of the tetramer.

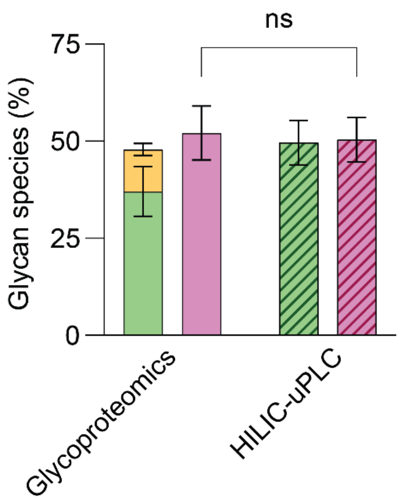


**Figure S8. Comparison of glycan species identified using glycoproteomics vs UHPLC.** Oligomannose glycan containing species in solid green, hybrid in yellow and complex in purple. Endo-H sensitive glycan species in striped green, endo-H resistant glycan species in striped purple.

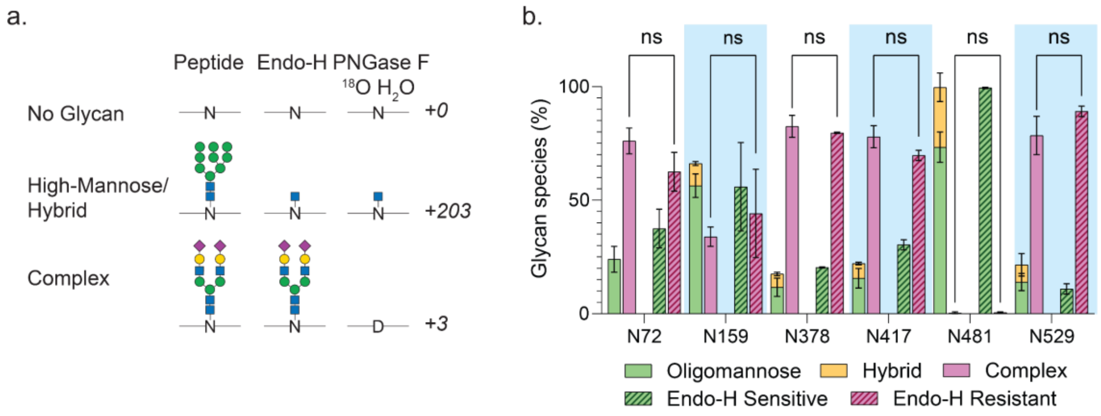


**Figure S9. Comparison on LFQ and $^{18}O$ labelling for glycoproteomic quantification.** a) Schematic of differential enzymatic treatment of glycopeptides with Endo-H and PNGaseF labelled with $^{18}O$. b) Relative quantification of glycan species present in glycopeptides, with comparisons between the complex assignments, ns = ($p<0.05$), n = 3 for N72, n = 2 for LFQ.

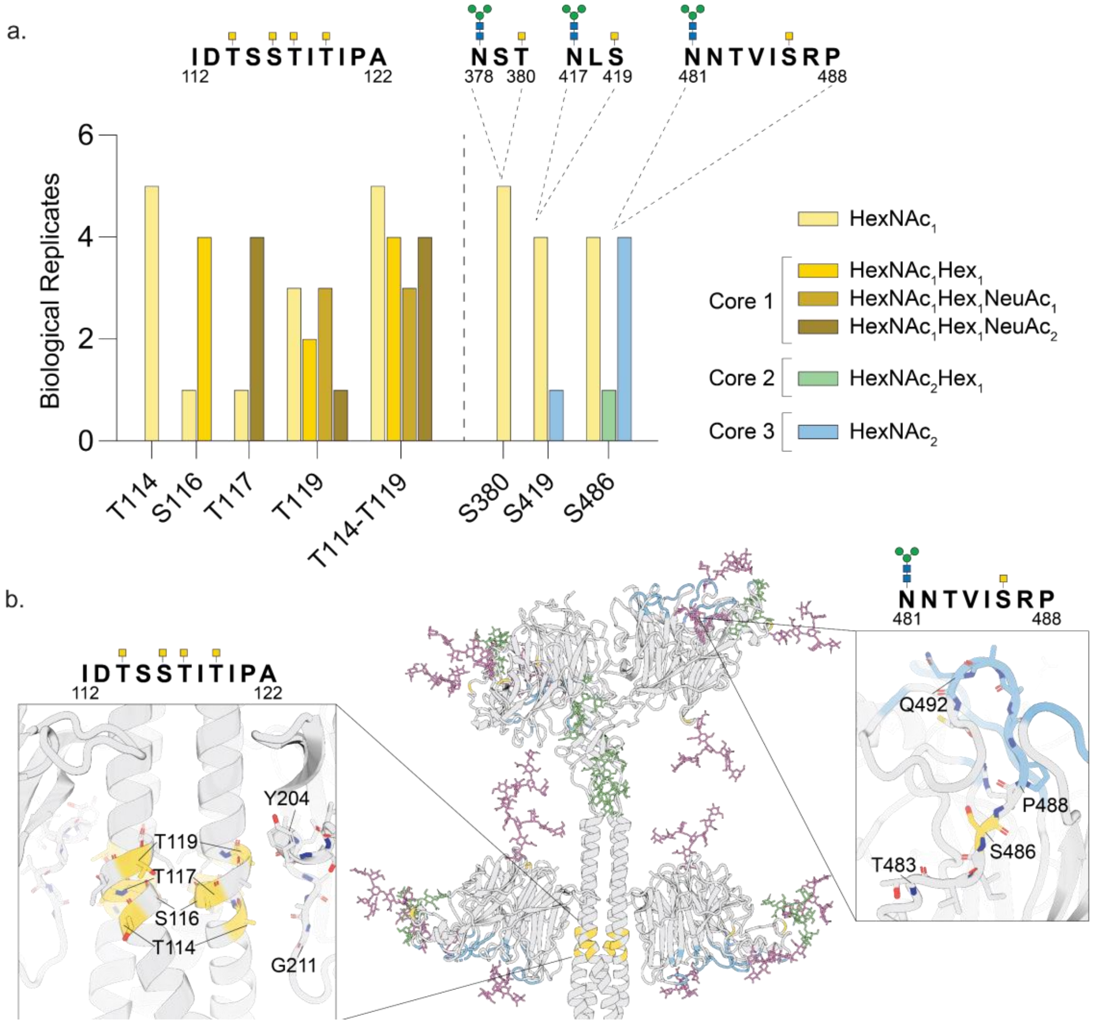


**Figure S10. O-glycosylation of the Nipah $G_M$ tetramer**. a) Positions of O-glycans sites identified using glycoproteomics revealed compositions corresponding to core 1-3 structures as shown in the figure. O-glycan sites are highlighted above by GalNAc residues (yellow square) and N-glycan sites are highlighted by $Man_3GlcNAc_2$. Only sites that were identified in ≥4 biological replicates out of 6 were included. b) Location of O-glycans on the stalk and near the ephrin B2 binding region (blue). N-glycans are coloured similarly to main text Figure 2.

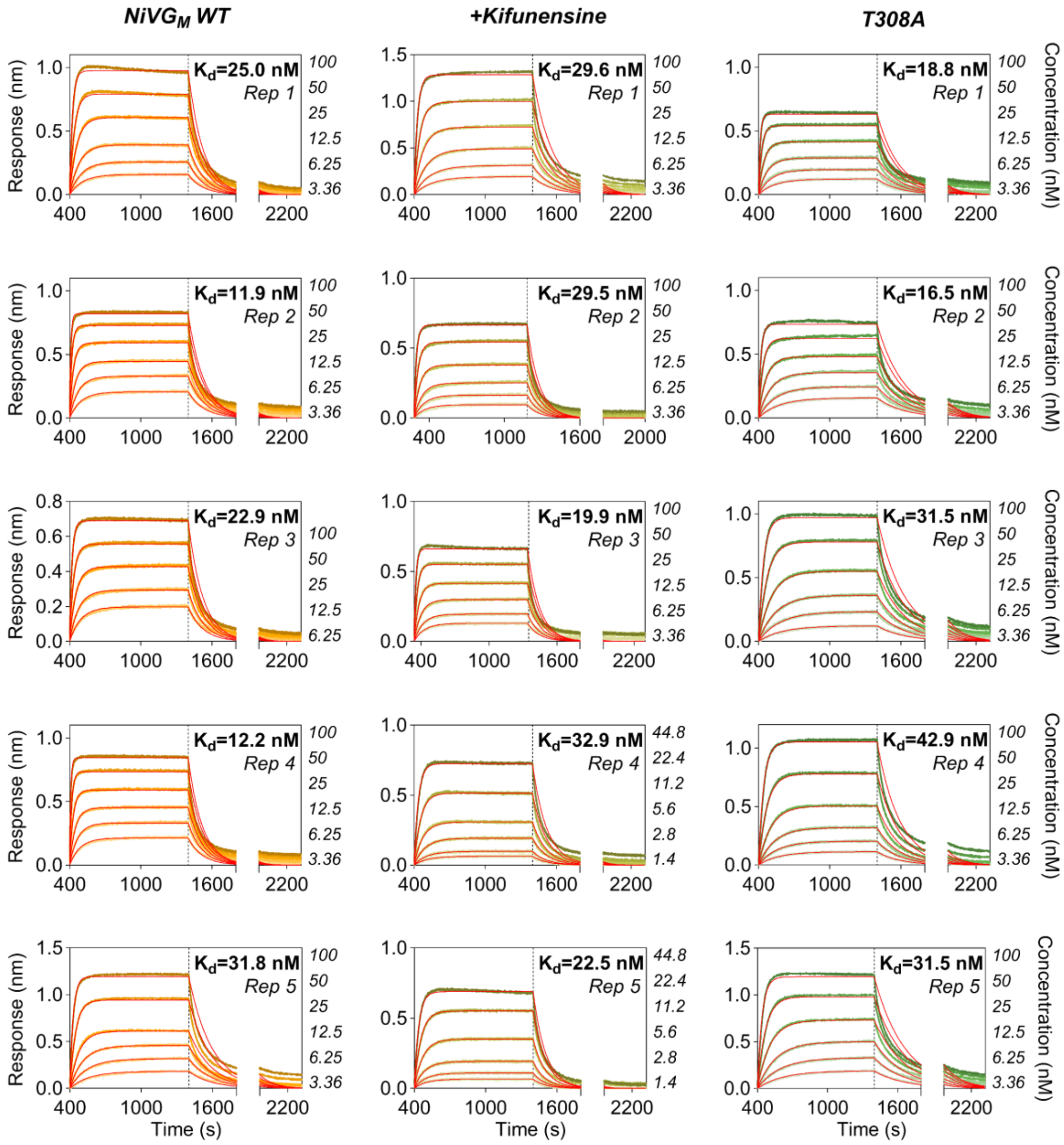


**Figure S11. Biolayer interferometry of binding between ephrin B2 RBD and Nipah $G_M$ head domain WT, +kifunensine and T308A, S378A, S419A, N481D, T483A and T531A mutants.** Fits are coloured red and steady-state approximation was used to determine $K_d$ values. The concentration of Nipah $G_M$ is shown to the right of each sensorgram.

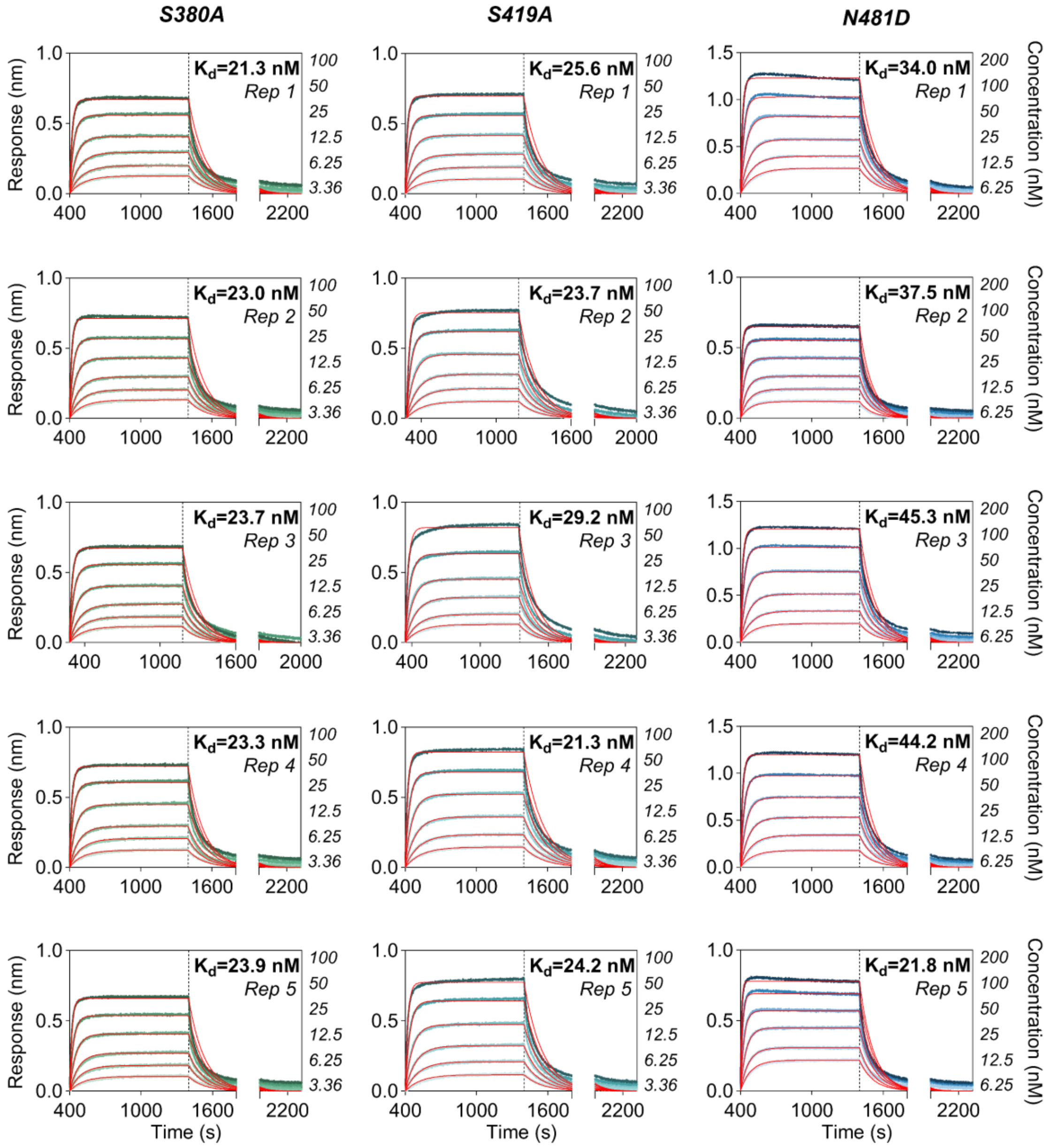


**Figure S11 continued. Biolayer interferometry of binding between ephrin B2 RBD and Nipah $G_M$ head domain WT, +kifunensine and T308A, S378A, S419A, N481D, T483A and T531A mutants.** Fits are coloured red and steady-state approximation was used to determine $K_d$ values. The concentration of Nipah $G_M$ is shown to the right of each sensorgram.

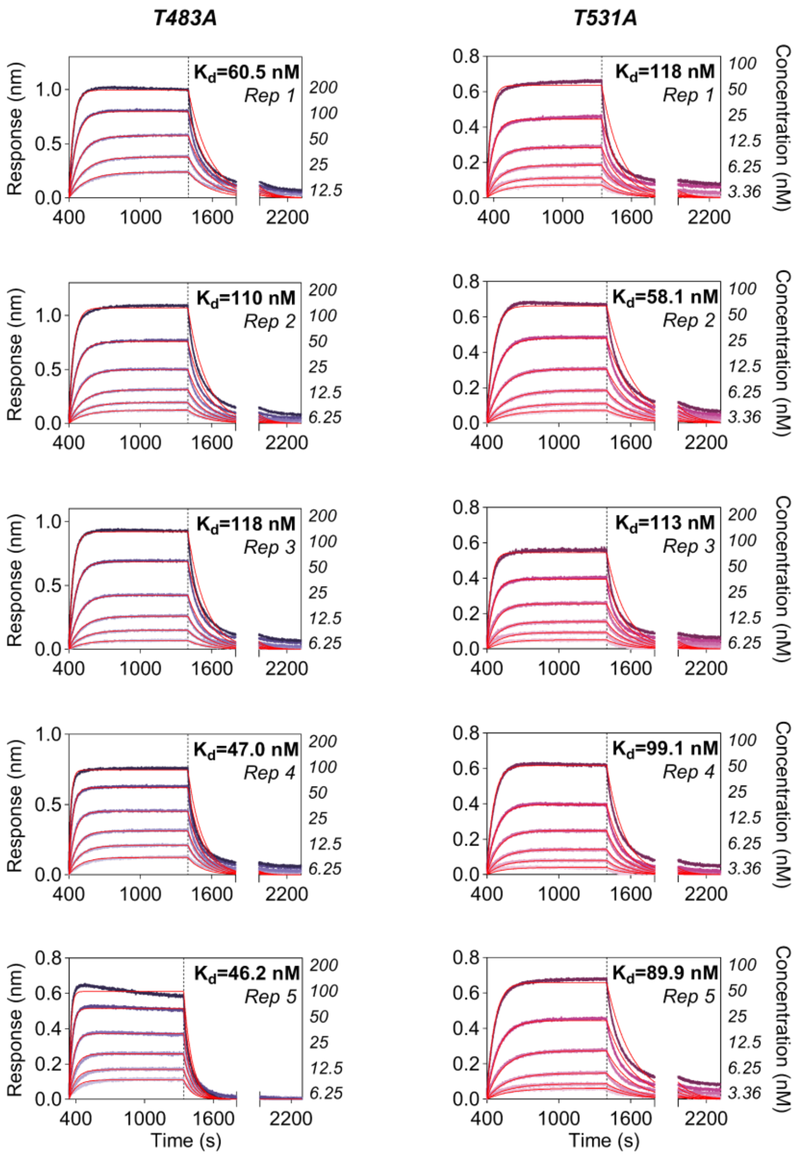


**Figure S11 continued. Biolayer interferometry of binding between ephrin B2 RBD and Nipah $G_M$ head domain WT, +kifunensine and T308A, S378A, S419A, N481D, T483A and T531A mutants.** Fits are coloured red and steady-state approximation was used to determine $K_d$ values. The concentration of Nipah $G_M$ is shown to the right of each sensorgram.

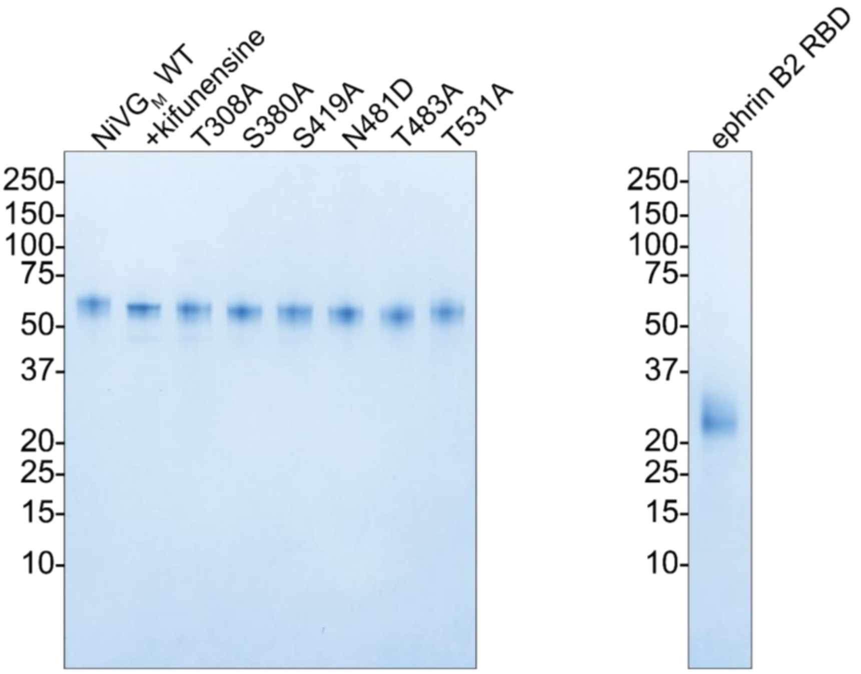


**Figure S12. Coomassie stained SDS-PAGE gel of Nipah G$_M$ head mutants and ephrin B2 RBD.** Expected molecular weights (based on peptide mass) are ~51 kDa for Nipah G$_M$ and ~18.8 kDa for ephrin B2 RBD.

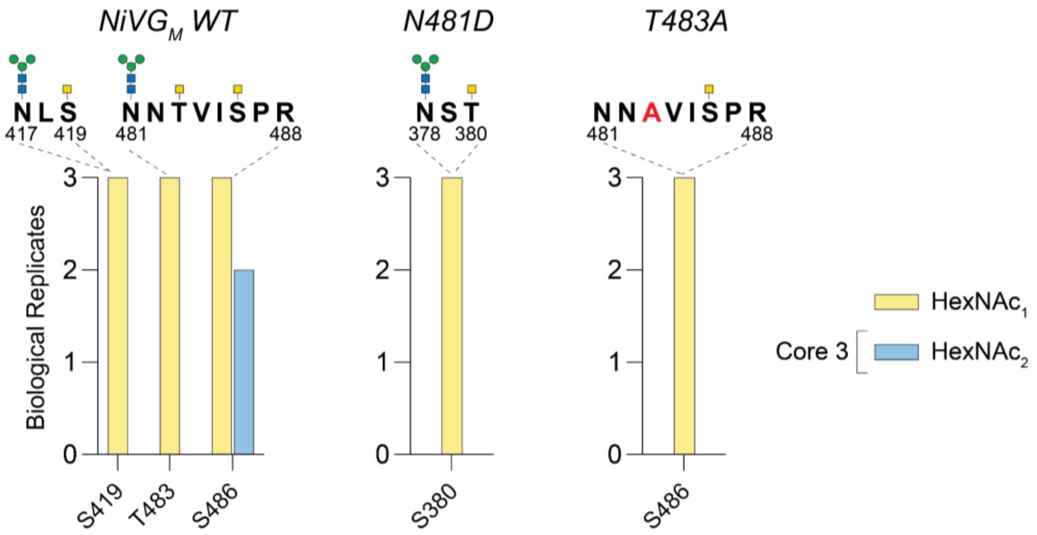


**Figure S13. O-glycosylation of Nipah $G_M$ head, N481D and T483A mutants.** Positions of O-glycans sites identified using glycoproteomics revealed compositions corresponding to a single $HexNAc_1$ and core-3 structures as shown in the figure. O-glycan sites are highlighted above by GalNAc residues (yellow square) and N-glycan sites are highlighted by $Man_3GlcNAc_2$. Only sites identified in ≥2 biological replicates out of 3 are shown. Mutation site shown in red.

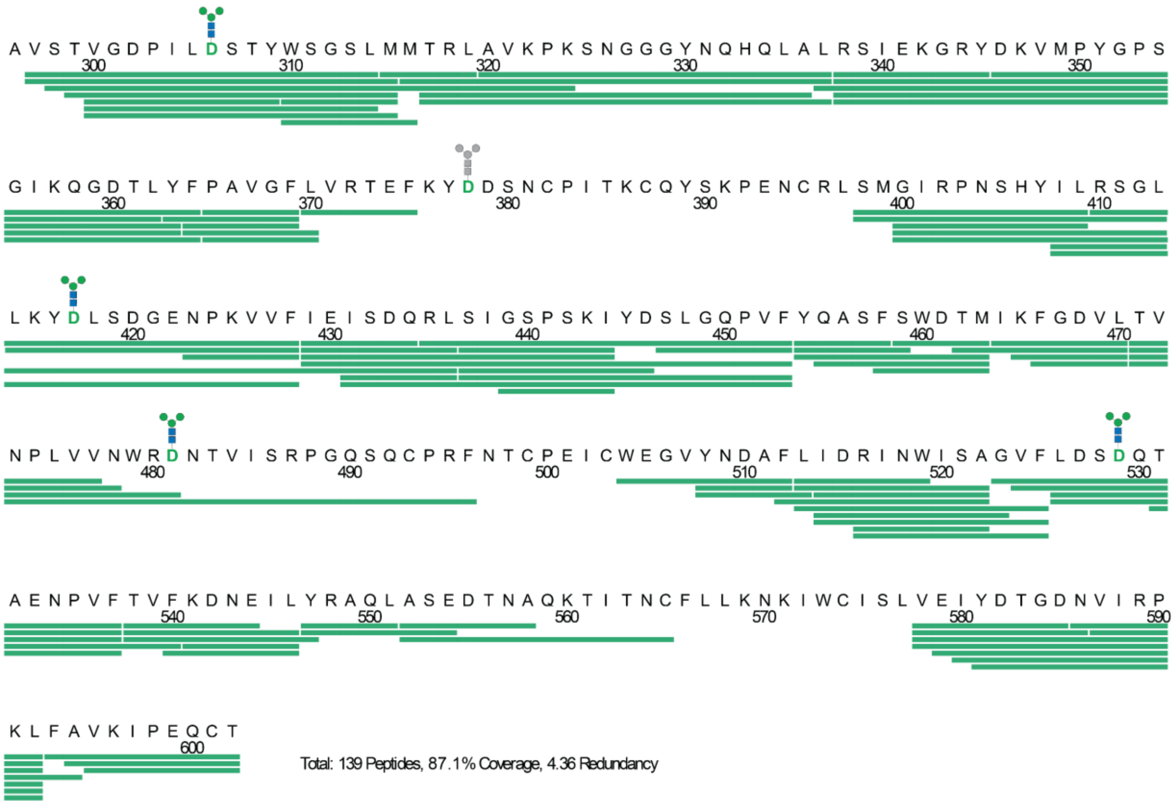


**Figure S14. Effective sequence coverage of NiVG$_M$ WT for HDX-MS.** N-glycans shown with Man$_3$GlcNAc$_2$ and at the green letter D, as N-glycans were enzymatically deamidated using PNGase Rc [5]. Grey Man$_3$GlcNAc$_2$ glycan cartoon is a site that is not covered.

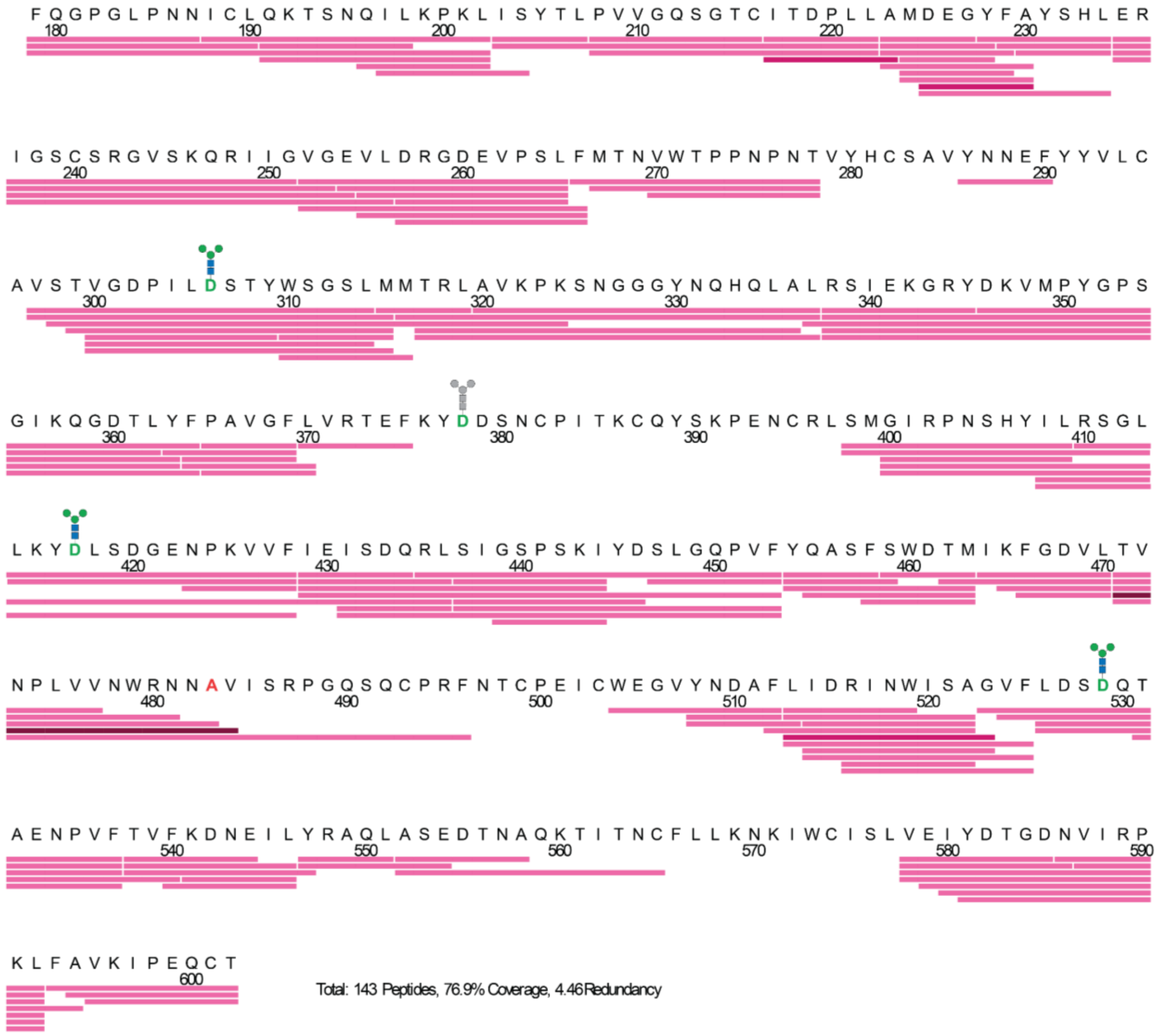

**Figure S15. Effective sequence coverage of NiVG$_M$ T483A HDX-MS.** N-glycans shown with $Man_3GlcNAc_2$ and at the green letter D, as N-glycans were enzymatically deamidated using PNGase Rc [5]. Grey $Man_3GlcNAc_2$ glycan cartoon is a site that is not covered. Dark pink peptides are unique to the mutant dataset and dark maroon peptides are unique to the T483A dataset.

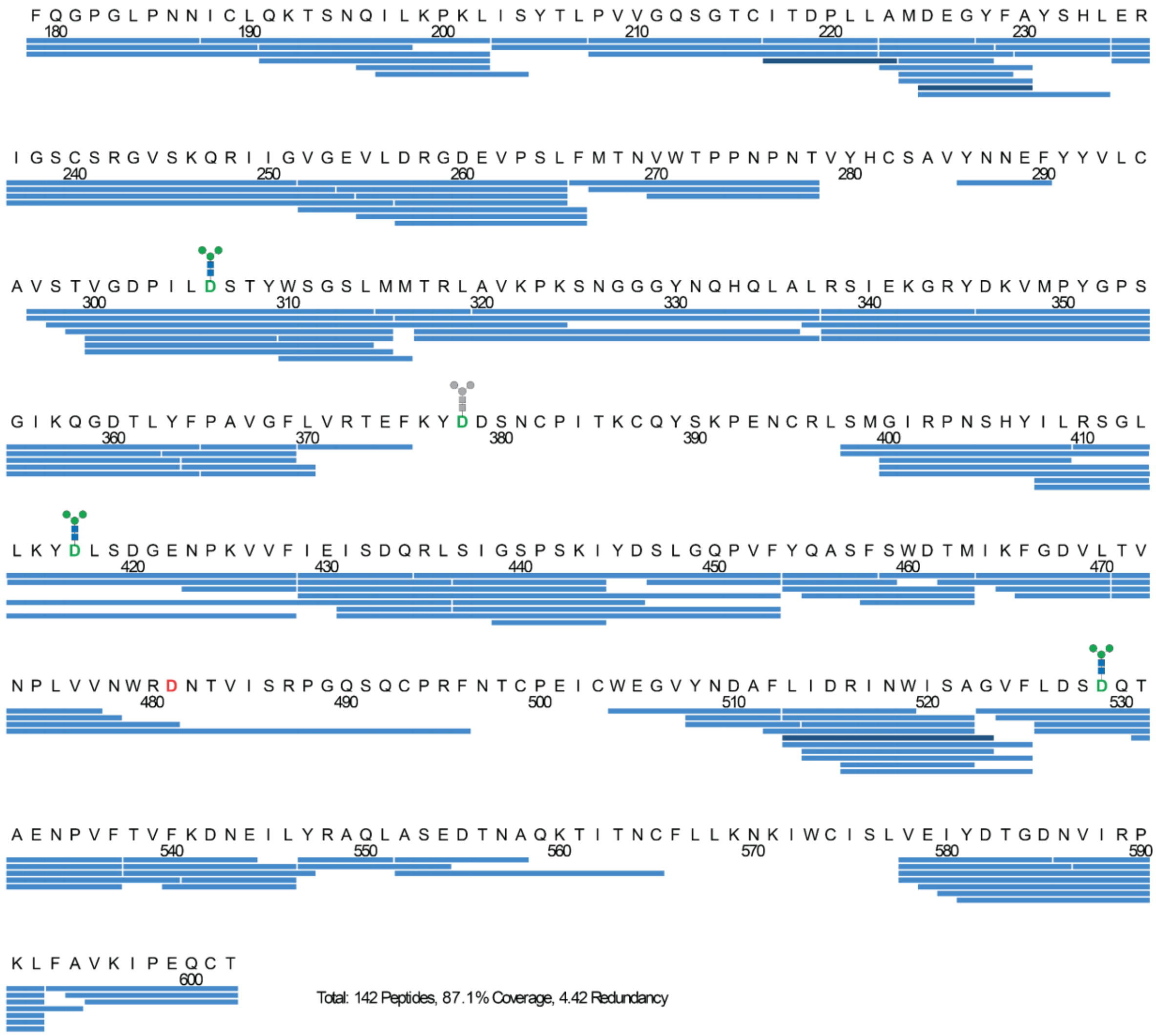

**Figure S16. Effective sequence coverage of NiVG$_M$ N481D HDX-MS.** N-glycans shown with $Man_3GlcNAc_2$ and at the green letter D, as N-glycans were enzymatically deamidated using PNGase Rc [5]. Grey $Man_3GlcNAc_2$ glycan cartoon is a site that is not covered. Dark blue peptides are unique to the mutant dataset.

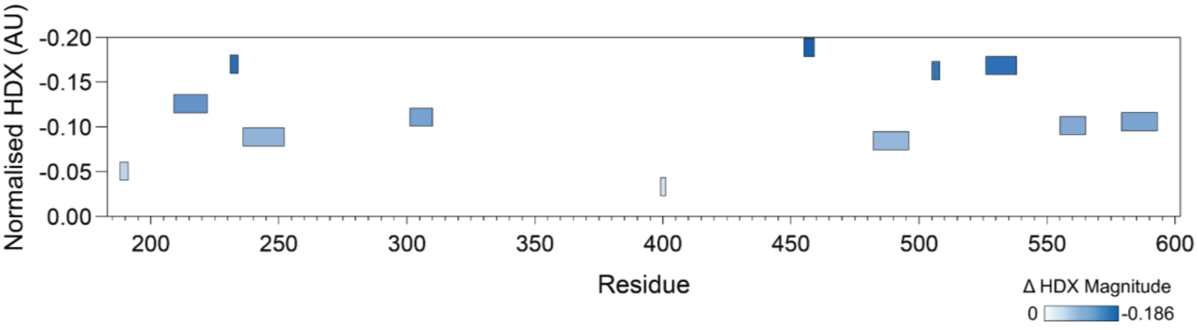


**Figure S17. Woods plot of statistically significant HDX peptides**. Peptide length, position in sequence, and HDX response, normalised to the length of the peptide, is displayed in this plot. Intensity of blue correlates to the HDX magnitude displayed by the peptides.

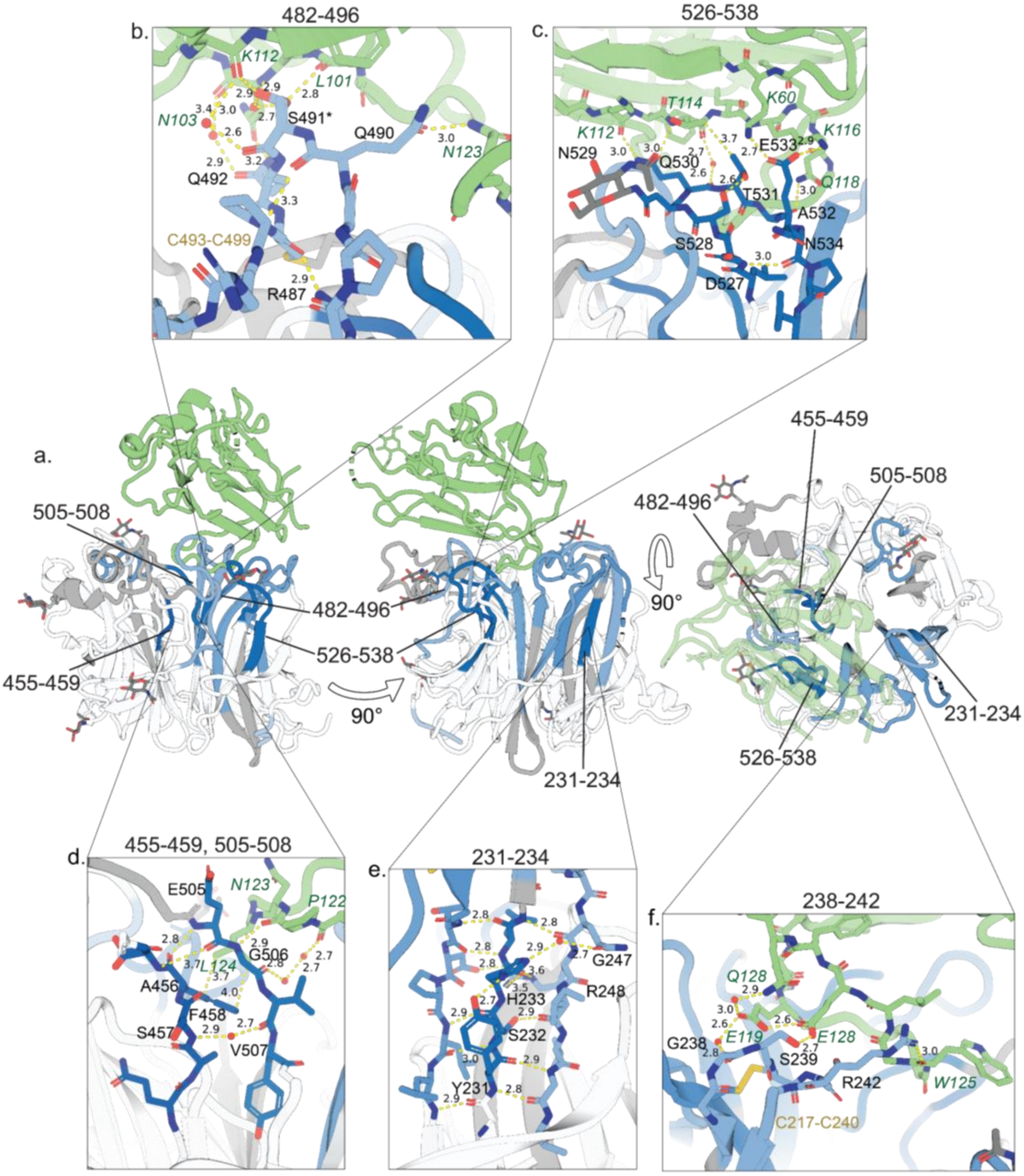


**Figure S18. Overview of HDX changes upon ephrin B2 binding to Nipah G$_M$ Head.** a) Three-way view of Nipah G$_M$ head and ephrin B2 complex (PDB:2VSM). b) Intrapeptide and peptide-ephrin B2 interactions for 482-496. c) Intrapeptide and peptide-ephrin B2 interactions for 526-538. d) Intrapeptide and peptide-ephrin B2 interactions for 455-459 and 505-508. e) Intrapeptide interactions for 231-234. Only amino acids involved in interactions other than the backbone beta-sheet interaction are labelled. f) Intrapeptide and peptide-ephrin B2 interactions for 238-242. Response normalised to the length of the peptide, and coloured based on response intensity. Distances shown by dashed yellow lines, labelled in Å. Head amino acid labels in black, receptor in green and disulfide bonds in yellow.

| Accession | Species | Year | Country | Host |
|---|---|---|---|---|
| AAC83193*† | Henipavirus hendraense | 1994 | Australia | Equus ferus callabus |
| XJU66854 | Henipavirus nipahense | 1998 | Malaysia | Homo sapiens |
| QHR78992 | Henipavirus nipahense | 1999 | Malaysia | Homo sapiens |
| QHR78983 | Henipavirus nipahense | 1999 | Malaysia | Sus scrofa domesticus |
| QHR78956 | Henipavirus nipahense | 1999 | Malaysia | Sus scrofa domesticus |
| QHR78965 | Henipavirus nipahense | 1999 | Malaysia | Sus scrofa domesticus |
| QHR78974 | Henipavirus nipahense | 1999 | Malaysia | Sus scrofa domesticus |
| QHR79001 | Henipavirus nipahense | 1999 | Malaysia | Canis lupus familiaris |
| AAK50545 | Henipavirus nipahense | 1999 | Malaysia | Homo sapiens |
| AAK50554 | Henipavirus nipahense | 1999 | Malaysia | Homo sapiens |
| CAD92363 | Henipavirus nipahense | 1999 | Malaysia | Homo sapiens |
| AAK29088* | Henipavirus nipahense | 1999 | Malaysia | Homo sapiens |
| CAF25497 | Henipavirus nipahense | 1999 | Malaysia | Sus scrofa |
| CAD92351 | Henipavirus nipahense | 1999 | Malaysia | Sus scrofa |
| CAD92357 | Henipavirus nipahense | 1999 | Malaysia | Sus scrofa |
| QDJ04463 | Henipavirus nipahense | 2003 | Cambodia | Pteropus lylei |
| AAY43916 | Henipavirus nipahense | 2004 | Bangladesh | Homo sapiens |
| QHR79010 | Henipavirus nipahense | 2004 | Bangladesh | Homo sapiens |
| QHR79019 | Henipavirus nipahense | 2004 | Bangladesh | Homo sapiens |
| QHR79028 | Henipavirus nipahense | 2004 | Bangladesh | Homo sapiens |
| QHR79037 | Henipavirus nipahense | 2004 | Bangladesh | Homo sapiens |
| ACT32615 | Henipavirus nipahense | 2007 | India | Homo sapiens |
| AEZ01382 | Henipavirus nipahense | 2008 | Bangladesh | Homo sapiens |
| AEZ01389 | Henipavirus nipahense | 2008 | Bangladesh | Homo sapiens |
| QHR79046 | Henipavirus nipahense | 2008 | Bangladesh | Homo sapiens |
| QHR79109 | Henipavirus nipahense | 2011 | Bangladesh | Homo sapiens |
| QHR79064 | Henipavirus nipahense | 2011 | Bangladesh | Homo sapiens |
| QHR79082 | Henipavirus nipahense | 2011 | Bangladesh | Homo sapiens |
| QHR79118 | Henipavirus nipahense | 2011 | Bangladesh | Homo sapiens |
| QHR79091 | Henipavirus nipahense | 2011 | Bangladesh | Homo sapiens |
| QHR79100 | Henipavirus nipahense | 2011 | Bangladesh | Homo sapiens |
| QHR79055 | Henipavirus nipahense | 2011 | Bangladesh | Homo sapiens |
| QHR79127 | Henipavirus nipahense | 2011 | Bangladesh | Homo sapiens |
| QHR79136 | Henipavirus nipahense | 2012 | Bangladesh | Homo sapiens |
| QHR79154 | Henipavirus nipahense | 2012 | Bangladesh | Homo sapiens |
| QHR79163 | Henipavirus nipahense | 2012 | Bangladesh | Homo sapiens |

**Table S1. Nipah G protein accessions from isolates collected from 1998-2012 used in phylogenetic analysis.** *Reference sequence. † Hendra virus outgroup.

| Accession | Species | Year | Country | Host |
|---|---|---|---|---|
| QCY59092 | Henipavirus nipahense | 2013 | Bangladesh | Pteropus giganteus |
| QHR79172 | Henipavirus nipahense | 2013 | Bangladesh | Homo sapiens |
| QCY59033 | Henipavirus nipahense | 2013 | Bangladesh | Pteropus giganteus |
| QCY59039 | Henipavirus nipahense | 2013 | Bangladesh | Pteropus giganteus |
| QCY59045 | Henipavirus nipahense | 2013 | Bangladesh | Pteropus giganteus |
| QCY59050 | Henipavirus nipahense | 2013 | Bangladesh | Pteropus giganteus |
| QCY59056 | Henipavirus nipahense | 2013 | Bangladesh | Pteropus giganteus |
| QCY59062 | Henipavirus nipahense | 2013 | Bangladesh | Pteropus giganteus |
| QCY59068 | Henipavirus nipahense | 2013 | Bangladesh | Pteropus giganteus |
| QCY59074 | Henipavirus nipahense | 2013 | Bangladesh | Pteropus giganteus |
| QCY59080 | Henipavirus nipahense | 2013 | Bangladesh | Pteropus giganteus |
| QCY59086 | Henipavirus nipahense | 2013 | Bangladesh | Pteropus giganteus |
| QHR79253 | Henipavirus nipahense | 2014 | Bangladesh | Homo sapiens |
| QHR79244 | Henipavirus nipahense | 2014 | Bangladesh | Homo sapiens |
| QHR79226 | Henipavirus nipahense | 2014 | Bangladesh | Homo sapiens |
| QHR79235 | Henipavirus nipahense | 2014 | Bangladesh | Homo sapiens |
| QHR79181 | Henipavirus nipahense | 2015 | Bangladesh | Homo sapiens |
| QHR79190 | Henipavirus nipahense | 2015 | Bangladesh | Homo sapiens |
| XCK17070 | Henipavirus nipahense | 2017 | Bangladesh | Homo sapiens |
| XCK17079 | Henipavirus nipahense | 2017 | Bangladesh | Homo sapiens |
| UEK26006 | Henipavirus nipahense | 2017 | Thailand | Pteropus lylei |
| QBQ56705 | Henipavirus nipahense | 2018 | India | Homo sapiens |
| QBQ56723 | Henipavirus nipahense | 2018 | India | Homo sapiens |
| AWT50994 | Henipavirus nipahense | 2018 | India | Homo sapiens |
| QBQ56714 | Henipavirus nipahense | 2018 | India | Homo sapiens |
| XCK17097 | Henipavirus nipahense | 2019 | Bangladesh | Homo sapiens |
| QKV44068 | Henipavirus nipahense | 2019 | India | Pteropus giganteus |
| XCK17115 | Henipavirus nipahense | 2020 | Bangladesh | Homo sapiens |
| XCK17124 | Henipavirus nipahense | 2020 | Bangladesh | Homo sapiens |
| XCK17133 | Henipavirus nipahense | 2020 | Bangladesh | Homo sapiens |

**Table S2. Nipah G protein accessions from isolates collected from 2013-2020 used in phylogenetic analysis.**

| Accession | Species | Year | Country | Host |
|---|---|---|---|---|
| XCK17151 | Henipavirus nipahense | 2022 | Bangladesh | Homo sapiens |
| XCK17160 | Henipavirus nipahense | 2022 | Bangladesh | Homo sapiens |
| XCK17169 | Henipavirus nipahense | 2022 | Bangladesh | Homo sapiens |
| XCK17106 | Henipavirus nipahense | 2023 | Bangladesh | Homo sapiens |
| XCK17178 | Henipavirus nipahense | 2023 | Bangladesh | Homo sapiens |
| XCK17187 | Henipavirus nipahense | 2023 | Bangladesh | Homo sapiens |
| XCK17196 | Henipavirus nipahense | 2023 | Bangladesh | Homo sapiens |
| XCK17205 | Henipavirus nipahense | 2023 | Bangladesh | Homo sapiens |
| XJU66140 | Henipavirus nipahense | 2023 | Bangladesh | Homo sapiens |
| XCK17223 | Henipavirus nipahense | 2023 | Bangladesh | Homo sapiens |
| XJU66146 | Henipavirus nipahense | 2023 | Bangladesh | Homo sapiens |
| WVH15777 | Henipavirus nipahense | 2023 | India | Homo sapiens |
| WVH15795 | Henipavirus nipahense | 2023 | India | Homo sapiens |
| XPO94330 | Henipavirus nipahense | 2024 | Bangladesh | Homo sapiens |
| XPO94336 | Henipavirus nipahense | 2024 | Bangladesh | Homo sapiens |

**Table S3. Nipah G protein accessions from isolates collected from 2022-2024 used in phylogenetic analysis.**

| Peptide | Site | Experiment |
|---|---|---|
| -.ETGHHHHHHGQ**n**YTR.S | N72 | Both |
| K.IHEC**n**ISCPNPLPFR.E | N159 | Both |
| V.GDPIL**n**ST.Y | N306 | LFQ only |
| R.TEFKY**n**DSNCPITK.C | N378 | Both |
| K.Y**n**LSDGENPK.V | N417 | Both |
| R.**n**NTVISRPGQSQCPR.F | N481 | Both |
| R.INWISAGVFLDS**n**QTAENPVFTVFK.D | N529 | LFQ only |
| D.S**n**QTAENPVFTVFK.D | N529 | $^{18}$O only |

**Table S4. Glycopeptides used in for abundance analysis and comparison.**

| Construct | Sequence |
|---|---|
| **Nipah $G_M$ ectodomain (71-602)** | atggggatccttcccagccctgggatgcctgcgctgctctccctcgtgagccttctctccgtgctgctgatgggttgcgt<br>agctgaaaccggccatcaccatcaccatcacggccaaaattacacaagatcaacagacaatcaggccgtgatc<br>aaagatgcgttgcagggtatccaacagcagatcaaagggcttgctgacaaaatcggcacagagatagggccca<br>aagtatcactgattgacacatccagtaccattactatcccagctaacattgggctgttaggttcaaagatcagccagt<br>cgactgcaagtataaatgagaatgtgaatgaaaaatgcaaattcacactgcctcccttgaaaatccacgaatgtaa<br>catttcttgtcctaacccactcccttttagagagtataggccacagacagaaggggtgagcaatctagtaggattacc<br>taataatatttgcctgcaaaagacatctaatcagatattgaagccaaagctgatttcatacactttacccgtagtcggtc<br>aaagtggtacctgtatcacagacccattgctggctatggacgagggctattttgcatatagccacctggaaagaatc<br>ggatcatgttcaagaggggtctccaaacaaagaataataggagttggagaggtactagacagaggtgatgaagtt<br>ccttctttatttatgaccaatgtctggaccccaccaaatccaaacaccgtttaccactgtagtgctgtatacaacaatga<br>attctattatgtactttgtgcagtgtcaactgttggagaccctattctgaatagcacctactggtccggatctctaatgatg<br>acccgtctagctgtgaaacccaagagtaatggtgggggttacaatcaacatcaacttgccctacgaagtatcgaga<br>aagggaggtatgataaagttatgccgtatggaccttcaggcatcaaacagggtgacaccctgtattttcctgctgtag<br>gatttttggtcaggacagagtttaaatacaatgattcaaattgtcccatcacgaagtgtcaatacagtaaacctgaaa<br>attgcaggctatctatggggattagaccaaacagccattatatccttcgatctggactattaaaatacaatctatcagat<br>ggggagaaccccaaagttgtattcattgaaatatctgatcaaagattatctattggatctcctagcaaaatctatgattc<br>tttgggtcaacctgttttctaccaagcgtcattttcatgggatactatgattaaatttggagatgttctaacagtcaaccctc<br>tggttgtcaattggcgtaataacacggtaatatcaagacccgggcaatcacaatgccctagattcaatacatgtcca<br>gagatctgctgggaaggagtttataatgatgcattcctaattgacagaatcaattggataagcgcgggtgtattccttg<br>acagcaatcagaccgcagaaaatcctgtttttactgtattcaaagataatgaaatactttatagggcacaactggcttc<br>tgaggacaccaatgcacaaaaaacaataactaattgttttctcttgaagaataagatttggtgcatatcactggtaga<br>aatttacgatacaggagataacgtgatccggcctaaactgtttgcggtcaaaatccctgaacaatgcacgtaa |
| **Nipah $G_M$ Head (183-602)** | atggggatccttcccagccctgggatgcctgcgctgctctccctcgtgagccttctctccgtgctgctgatgggttgcgt<br>agctgaaaccggccatcaccatcaccatcacggcctgaacgatatttttgaagcgcagaaaattgaatggcatga<br>aggtctggaagttctgtttcagggtccgggattacctaataatatttgcctgcaaaagacatctaatcagatattgaag<br>ccaaagctgatttcatacactttacccgtagtcggtcaaagtggtacctgtatcacagacccattgctggctatggacg<br>agggctattttgcatatagccacctggaaagaatcggatcatgttcaagaggggtctccaaacaaagaataatagg<br>agttggagaggtactagacagaggtgatgaagttccttctttatttatgaccaatgtctggaccccaccaaatccaaa<br>caccgtttaccactgtagtgctgtatacaacaatgaattctattatgtactttgtgcagtgtcaactgttggagaccctatt<br>ctgaatagcacctactggtccggatctctaatgatgacccgtctagctgtgaaacccaagagtaatggtgggggtta<br>caatcaacatcaacttgccctacgaagtatcgagaaagggaggtatgataaagttatgccgtatggaccttcaggc<br>atcaaacagggtgacaccctgtattttcctgctgtaggatttttggtcaggacagagtttaaatacaatgattcaaattgt<br>cccatcacgaagtgtcaatacagtaaacctgaaaattgcaggctatctatggggattagaccaaacagccattatat<br>ccttcgatctggactattaaaatacaatctatcagatggggagaaccccaaagttgtattcattgaaatatctgatcaa<br>agattatctattggatctcctagcaaaatctatgattctttgggtcaacctgttttctaccaagcgtcattttcatgggatact<br>atgattaaatttggagatgttctaacagtcaaccctctggttgtcaattggcgtaataacacggtaatatcaagacccg<br>ggcaatcacaatgccctagattcaatacatgtccagagatctgctgggaaggagtttataatgatgcattcctaattg<br>acagaatcaattggataagcgcgggtgtattccttgacagcaatcagaccgcagaaaatcctgtttttactgtattcaa<br>agataatgaaatactttatagggcacaactggcttctgaggacaccaatgcacaaaaaacaataactaattgttttct<br>cttgaagaataagatttggtgcatatcactggtagaaatttacgatacaggagataacgtgatccggcctaaactgttt<br>gcggtcaaaatccctgaacaatgcacgtaa |
| **Ephrin B2 RBD (28-165)** | atggggatccttcccagccctgggatgcctgcgctgctctccctcgtgagccttctctccgtgctgctgatgggttgcgt<br>agctgaaaccggcatagttttagagcctatctattggaattcctcgaactccaaatttctacctggacaaggactggta<br>ctatacccacagataggagacaaattggatattatttgccccaaagtggactctaaaactgttggccagtatgaatat<br>tataaagtttatatggttgataaagaccaagcagacagatgcactattaagaaggaaaatacccctctcctcaactg<br>tgccaaaccagaccaagatatcaaattcaccatcaagtttcaagaattcagccctaacctctggggtctagaatttca<br>gaagaacaaagattattacattatatctacatcaaatgggtctttggagggcctggataaccaggagggaggggtg<br>tgccagacaagagccatgaagatcctcatgaaagttggaaaacatcaccatcaccatcactaa |

**Table S5. Coding sequences of constructs used to express Nipah $G_M$ ectodomain, Head and Ephrin B2 RBD.**

| Mutation site | Forward Primer (5'-3') | Reverse Primer (3'-5') | Annealing Temp (°C) |
|---|---|---|---|
| **T308A** | tctgaatagcgcgtactggtccg | atagggtctccaacagttg | 63 |
| **S380A** | atacaatgatgcgaattgtcccatc | ttaaactctgtcctgacc | 58 |
| **S419A** | atacaatctagcggatggggagaac | tttaatagtccagatcgaagg | 60 |
| **N481D** | caattggcgtgataacacggtaatatc | acaaccagagggttgac | 62 |
| **T483A** | gcgtaataacgcggtaatatcaag | caattgacaaccagagg | 58 |
| **T531A** | cagcaatcaggcggcagaaaatc | tcaaggaatacacccg | 59 |

**Table S6. Primers used to generate site directed mutants of Nipah $G_M$ Head.**

| Time (mins) | Solvent A (%) | Solvent B (%) | Flow (mL/min) |
|---|---|---|---|
| **0.0** | 22.0 | 78.0 | 0.50 |
| **48.5** | 54.1 | 45.9 | 0.50 |
| **49.5** | 100.0 | 0.0 | 0.25 |
| **54.5** | 100.0 | 0.0 | 0.25 |
| **56.5** | 22.0 | 78.0 | 0.50 |

**Table S7. uPLC running conditions for 2AA-labelled glycans.** Solvent A: 100 mM ammonium formate, pH 4.5; Solvent B: MeCN; Temperature 60 °C.

| Scan Range (m/z) | Scan Time (s) | Capillary voltage (kV) | Sample cone (V) | Extraction cone (V) | Source Temperature (ºC) | Polarity | ToF Operation | Gas |
|---|---|---|---|---|---|---|---|---|
| 50-2000 | 1 | 0.8-1.2 | 65 | 3.3 | 80 | Negative | Sensitivity Mode | He |

**Table S8. Ion mobility conditions for N-glycan analysis.**

| *m/z* | | Composition | | | | Ion |
| --- | --- | --- | --- | --- | --- | --- |
| Found | Calc. | Man | GlcNAc | Fuc | TFA | |
| 1347.6 | 1347.4 | 5 | 2 | 0 | 1 | $[M + TFA - H]^-$ |
| 1509.7 | 1509.5 | 6 | 2 | 0 | 1 | $[M + TFA - H]^-$ |
| 1575.8 | 1575.5 | 3 | 4 | 1 | 1 | $[M + TFA - H]^-$ |
| 1671.8 | 1671.5 | 7 | 2 | 0 | 1 | $[M + TFA - H]^-$ |
| 1833.9 | 1833.6 | 8 | 2 | 0 | 1 | $[M + TFA - H]^-$ |

**Table S9. Negative singly charged ion species identified for Nipah $G_M$ tetramer released glycans.** Ions were extracted via ion mobility prior to identification.

| *m/z* | | Composition | | | | | Ion |
|---|---|---|---|---|---|---|---|
| Found | Calc. | Man | GlcNAc | Gal | Fuc | NeuAc | |
| 1183.6 | 1183.4 | 3 | 4 | 2 | 1 | 2 | $[M - H_2]^{2-}$ |
| 1366.7 | 1366.0 | 3 | 5 | 3 | 1 | 2 | $[M - H_2]^{2-}$ |
| 1549.3 | 1548.5 | 3 | 6 | 4 | 1 | 2 | $[M - H_2]^{2-}$ |

**Table S10. Negative doubly charged ion species identified for Nipah $G_M$ tetramer released glycans.** Ions were extracted via ion mobility prior to identification.

| *m/z* | | Composition | | | | | Ion |
|---|---|---|---|---|---|---|---|
| Found | Calc. | Man | GlcNAc | Gal | Fuc | NeuAc | |
| 1007.8 | 1009.0 | 3 | 4 | 2 | 1 | 2 | $[M – H_3]^{3-}$ |
| 1129.5 | 1130.7 | 3 | 6 | 4 | 1 | 3 | $[M – H_3]^{3-}$ |
| 1226.6 | 1227.8 | 3 | 6 | 4 | 1 | 4 | $[M – H_3]^{3-}$ |

**Table S11. Negative triply charged ion species identified for Nipah $G_M$ tetramer released glycans.** Ions were extracted via ion mobility prior to identification.

| Time (mins) | Solvent A (%) | Solvent B (%) |
|---|---|---|
| **0.0** | 98.0 | 2.0 |
| **0.0** | 98.0 | 2.0 |
| **6.6** | 98.0 | 2.0 |
| **105.0** | 64.0 | 36.0 |
| **125.0** | 40.0 | 60.0 |
| **126.0** | 5.0 | 95.0 |
| **131.0** | 5.0 | 95.0 |
| **132.0** | 98.0 | 2.0 |
| **147.0** | 98.0 | 2.0 |

**Table S12. Glycoproteomics LC conditions.** Solvent A: LCMS-H2O, 0.1 % FA, Solvent B: MeCN, 0.1 % FA.

| Scan Range (m/z) | MS resolution | MS2 resolution | Isolation window (m/z) | Fragmentation | Collision Energy (%) | Polarity | Charge exclusion |
|---|---|---|---|---|---|---|---|
| 375-1500 | 70,000 | 35,000 | 2.0 | Stepped HCD | 27, 30, 33 | Positive | 1, ≥8 |

**Table S13. Glycoproteomics MS/MS conditions.** Data collected on a Q-Exactive Orbitrap MS.

| Phase | Time (s) |
|---|---|
| Baseline 1 | 60 |
| Baseline 2 | 60 |
| Baseline 3 | 60 |
| Loading | 100 |
| Baseline 4 | 60 |
| Baseline 5* | 60 |
| Association | 800-1000 |
| Dissociation | 800-1000 |

**Table S14. BLI run conditions.** Data collected on an Octet R8 at 5.0 Hz.

| Time (mins) | Solvent A (%) | Solvent B (%) |
|---|---|---|
| **0.0** | 92 | 8 |
| **7.0** | 65 | 35 |
| **7.5** | 15 | 85 |
| **8.5** | 15 | 85 |
| **9.0** | 92 | 8 |
| **12.0** | 92 | 8 |

**Table S15. HDX LC conditions.** Solvent A: LCMS-H2O, 0.23% FA , Solvent B: MeCN, 0.23% FA, flow rate 40 µL/min. Temperature 1 °C

| Scan Range (m/z) | Scan Time (s) | Trap Collision Energy (V) | Transfer Collision Energy (V) | Ramp Transfer (V) | Cone Voltage (V) | Polarity | ToF Operation | Gas |
|---|---|---|---|---|---|---|---|---|
| 50-2000 | 0.4 | 6 | 4 | 20-30 | 50 | Positive | Resolution Mode | He |

**Table S16. HDX MSe conditions.** Data collected for references only, collected on a Synapt G2Si.

| Scan Range (m/z) | Scan Time (s) | Trap Collision Energy (V) | Transfer Collision Energy (V) | Cone Voltage (V) | IMS wave velocity (m/s) | Transfer wave velocity (m/s) | Polarity | ToF Operation | Gas |
|---|---|---|---|---|---|---|---|---|---|
| 20-2000 | 0.4 | 6 | 4 | 30 | 675 | 175 | Positive | Resolution Mode, Normal sensitivity | He |

**Table S17. HDX IM/MS conditions.** Data collected for timepoints and MaxD only, collected on a Synapt G2Si.

References

(1) Jones, D. T.; Taylor, W. R.; Thornton, J. M. The rapid generation of mutation data matrices from protein sequences. *Comput Appl Biosci* **1992**, *8* (3), 275-282. DOI: 10.1093/bioinformatics/8.3.275 From PubMed.
(2) Tamura, K.; Stecher, G.; Kumar, S. MEGA11: Molecular Evolutionary Genetics Analysis Version 11. *Mol Biol Evol* **2021**, *38* (7), 3022-3027. DOI: 10.1093/molbev/msab120 From PubMed.
(3) Cao, L.; Diedrich, J. K.; Ma, Y.; Wang, N.; Pauthner, M.; Park, S.-K. R.; Delahunty, C. M.; McLellan, J. S.; Burton, D. R.; Yates, J. R.; et al. Global site-specific analysis of glycoprotein N-glycan processing. *Nat Protoc* **2018**, *13* (6), 1196-1212. DOI: 10.1038/nprot.2018.024 From PubMed.
(4) Mellacheruvu, D.; Wright, Z.; Couzens, A. L.; Lambert, J.-P.; St-Denis, N. A.; Li, T.; Miteva, Y. V.; Hauri, S.; Sardiu, M. E.; Low, T. Y.; et al. The CRAPome: a contaminant repository for affinity purification–mass spectrometry data. *Nature Methods 2013 10:8* **2013**, *10* (8). DOI: 10.1038/nmeth.2557.
(5) Lambert, T.; Gramlich, M.; Stutzke, L.; Smith, L.; Deng, D.; Kaiser, P. D.; Rothbauer, U.; Benesch, J. L. P.; Wagner, C.; Koenig, M.; et al. Development of a PNGase Rc Column for Online Deglycosylation of Complex Glycoproteins during HDX-MS. *J Am Soc Mass Spectrom* **2023**, *34* (11). DOI: 10.1021/jasms.3c00268.
(6) Houde, D.; Berkowitz, S. A.; Engen, J. R. The Utility of Hydrogen/Deuterium Exchange Mass Spectrometry in Biopharmaceutical Comparability Studies. *Journal of pharmaceutical sciences* **2010**, *100* (6). DOI: 10.1002/jps.22432.
(7) Calvaresi, V.; Wrobel, A. G.; Toporowska, J.; Hammerschmid, D.; Doores, K. J.; Bradshaw, R. T.; Parsons, R. B.; Benton, D. J.; Roustan, C.; Reading, E.; et al. Structural dynamics in the evolution of SARS-CoV-2 spike glycoprotein. *Nature Communications 2023 14:1* **2023**, *14* (1). DOI: 10.1038/s41467-023-36745-0.
