## Supporting File 2. HDX-MS uptake plots for "An evolutionary distinct Nipah G glycosylation site provides stability for receptor engagement"

**Supporting File 4 – HDX-MS Uptake Plots**


### Authors

Tia É. Hawkins[a,b,c], Valeria Calvaresi[b,d], Sean A. Burnap[b,d], Liang Wu[a,e*], Weston B. Struwe[b,d*]

### Addresses

a. The Rosalind Franklin Institute, Harwell Science & Innovation Campus, Harwell OX11 0FA, U.K.

b. Kavli Institute for Nanoscience Discovery, University of Oxford, OX1 3QU, U.K.

c. Department of Chemistry, University of Oxford, Oxford OX1 3TA, U.K.

d. Department of Biochemistry, University of Oxford, Oxford, OX1 3QU, U.K.

e. Division of Structural Biology, Nuffield Department of Medicine, University of Oxford, Oxford, OX3 7BN, U.K.

### Correspondents

*, *


Uptake plots were exported from DynamX 3.0. Pages 2-19 contain NiVG WT +/- Ephrin B2 uptake plots, pages 20-37 the NiVG WT, N481D and T483A uptake plots and page 37 is T483A unique peptides.

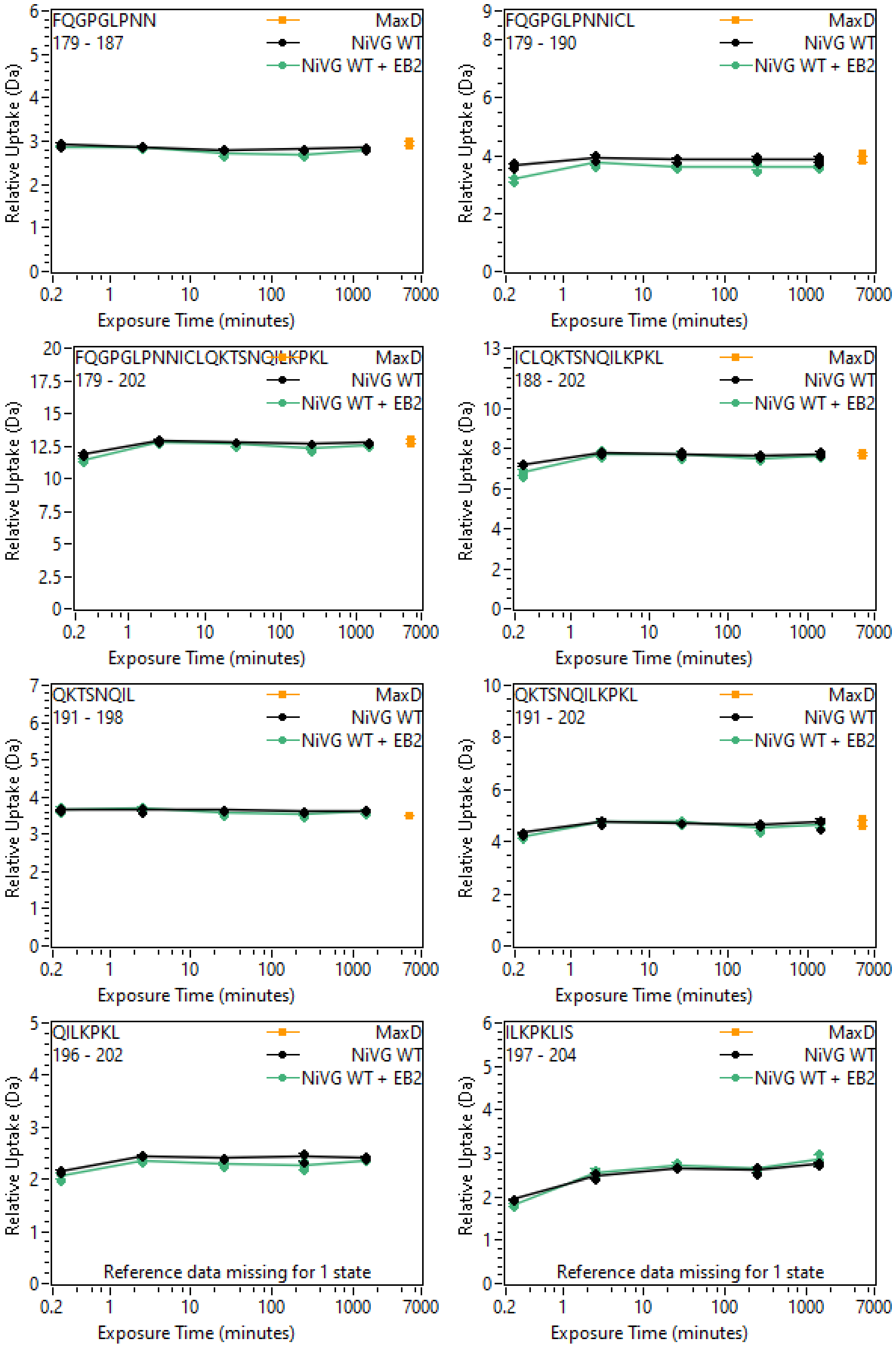
FQGPGLPNN
179 - 187
MaxD
NiVG WT
NiVG WT + EB2
Relative Uptake (Da)
Exposure Time (minutes)
FQGPGLPNNICL
179 - 190
MaxD
NiVG WT
NiVG WT + EB2
Relative Uptake (Da)
Exposure Time (minutes)
FQGPGLPNNICLQKTSNQILKPKL
179 - 202
MaxD
NiVG WT
NiVG WT + EB2
Relative Uptake (Da)
Exposure Time (minutes)
ICLQKTSNQILKPKL
188 - 202
MaxD
NiVG WT
NiVG WT + EB2
Relative Uptake (Da)
Exposure Time (minutes)
QKTSNQIL
191 - 198
MaxD
NiVG WT
NiVG WT + EB2
Relative Uptake (Da)
Exposure Time (minutes)
QKTSNQILKPKL
191 - 202
MaxD
NiVG WT
NiVG WT + EB2
Relative Uptake (Da)
Exposure Time (minutes)
QILKPKL
196 - 202
MaxD
NiVG WT
NiVG WT + EB2
Reference data missing for 1 state
Relative Uptake (Da)
Exposure Time (minutes)
ILKPKLIS
197 - 204
MaxD
NiVG WT
NiVG WT + EB2
Reference data missing for 1 state
Relative Uptake (Da)
Exposure Time (minutes)

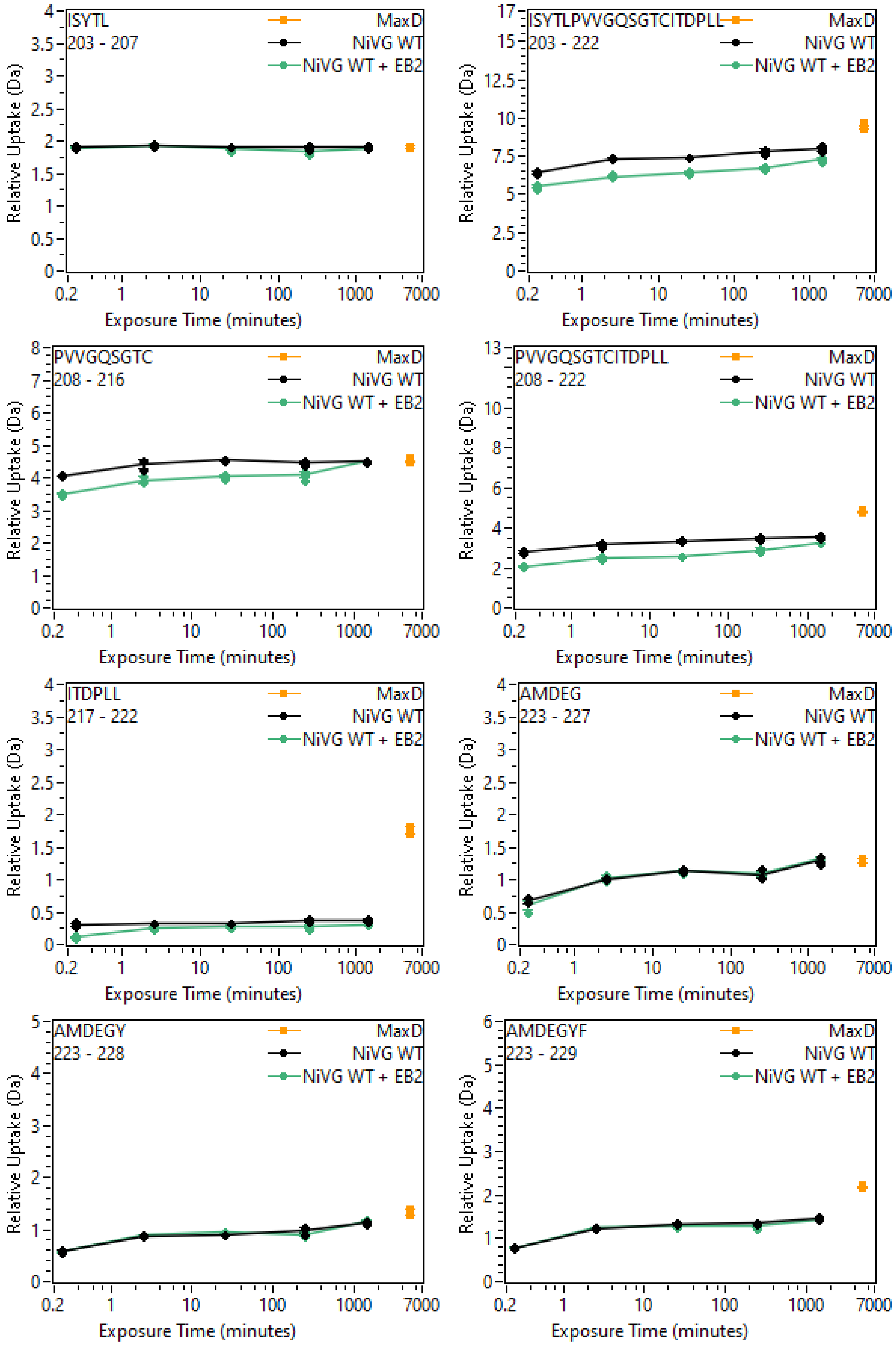
ISYTL
203 - 207
ISYTLPVVGQSGTCITDPLL
203 - 222
PVVGQSGTC
208 - 216
PVVGQSGTCITDPLL
208 - 222
ITDPLL
217 - 222
AMDEG
223 - 227
AMDEGY
223 - 228
AMDEGYF
223 - 229
MaxD
NiVG WT
NiVG WT + EB2
Relative Uptake (Da)
Exposure Time (minutes)
0.2
1
10
100
1000
7000

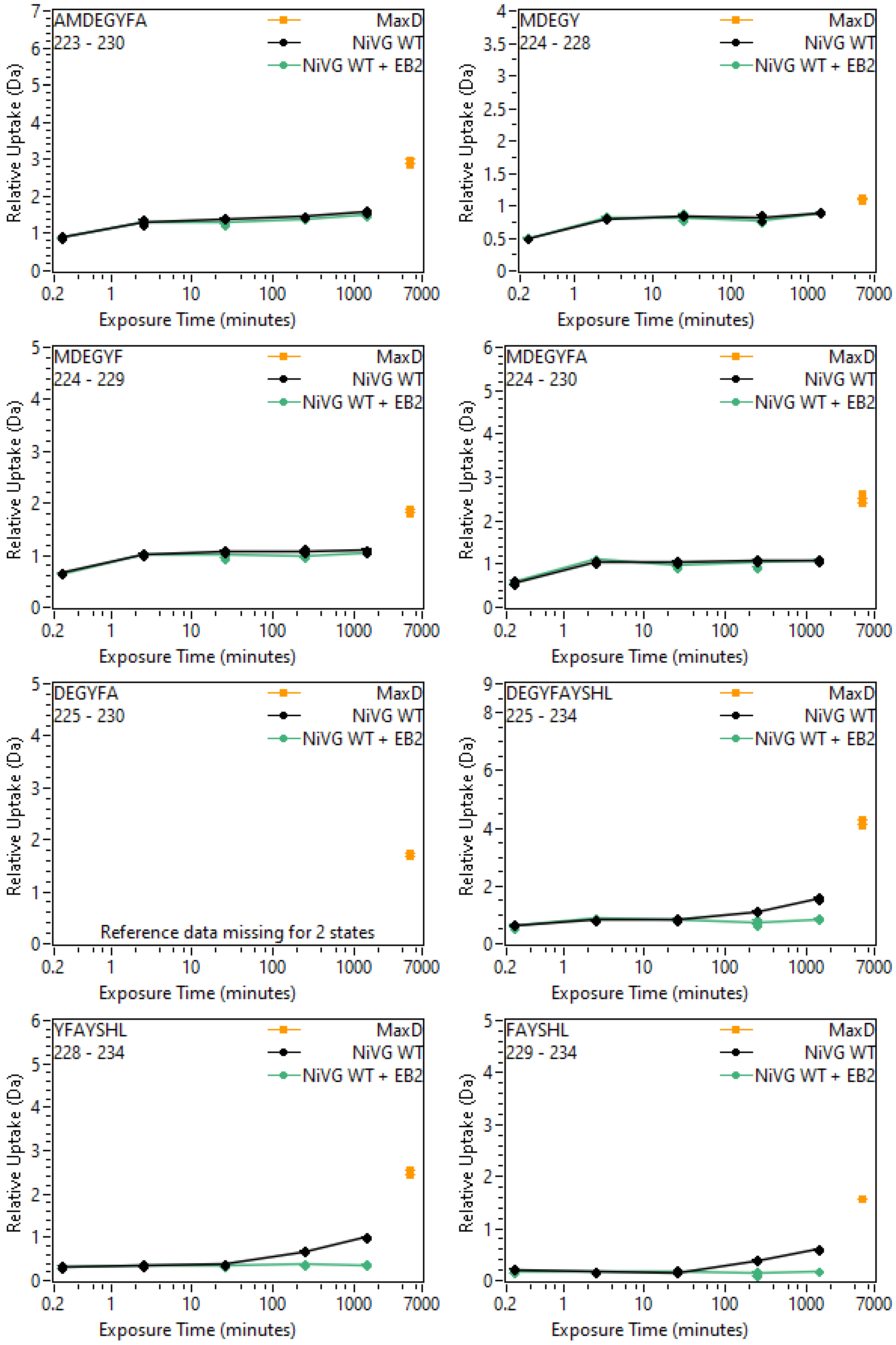
AMDEGYFA
223 - 230
MaxD
NiVG WT
NiVG WT + EB2
MDEGY
224 - 228
MDEGYF
224 - 229
MDEGYFA
224 - 230
DEGYFA
225 - 230
Reference data missing for 2 states
DEGYFAYSHL
225 - 234
YFAYSHL
228 - 234
FAYSHL
229 - 234
Relative Uptake (Da)
Exposure Time (minutes)
0.2
1
10
100
1000
7000

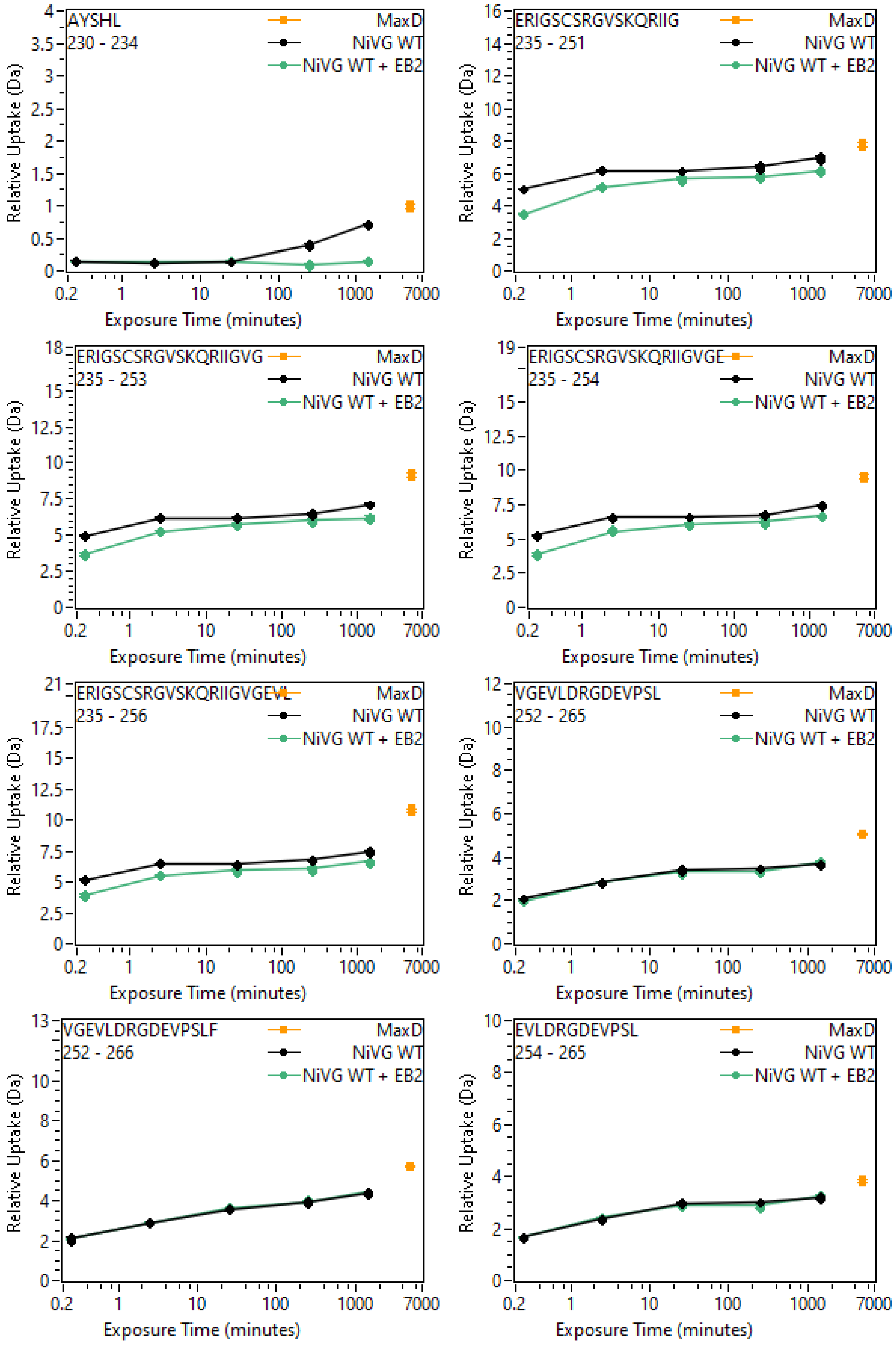
AYSHL
230 - 234
ERIGSCSRGVSKQRIIG
235 - 251
ERIGSCSRGVSKQRIIGVG
235 - 253
ERIGSCSRGVSKQRIIGVGE
235 - 254
ERIGSCSRGVSKQRIIGVGEVL
235 - 256
VGEVLDRGDEVPSL
252 - 265
VGEVLDRGDEVPSLF
252 - 266
EVLDRGDEVPSL
254 - 265
MaxD
NiVG WT
NiVG WT + EB2
Relative Uptake (Da)
Exposure Time (minutes)
0.2
1
10
100
1000
7000

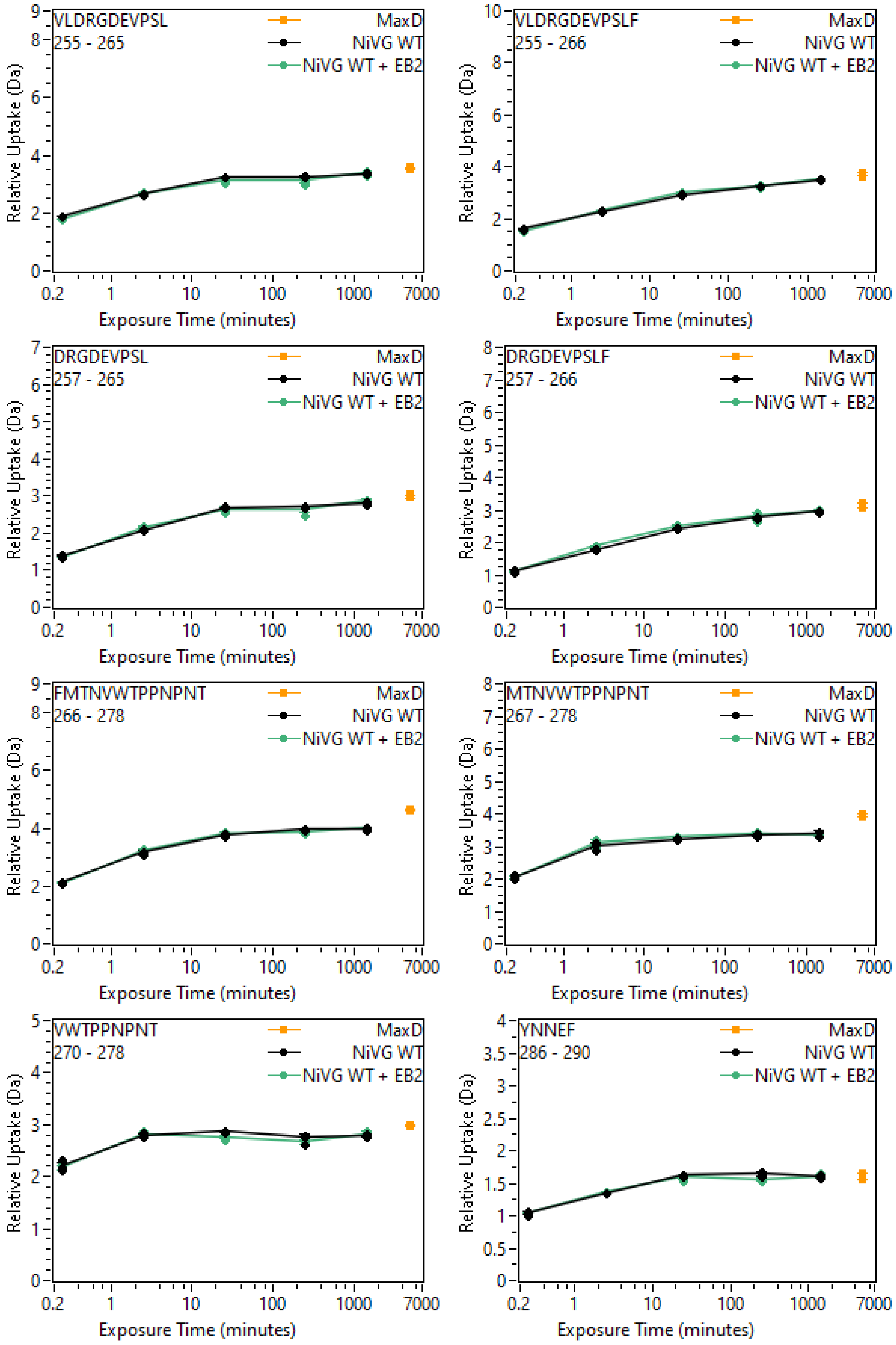
VLDRGDEVPSL
255 - 265
VLDRGDEVPSLF
255 - 266
DRGDEVPSL
257 - 265
DRGDEVPSLF
257 - 266
FMTNVWTPPNPNT
266 - 278
MTNVWTPPNPNT
267 - 278
VWTPPNPNT
270 - 278
YNNEF
286 - 290
MaxD
NiVG WT
NiVG WT + EB2
Relative Uptake (Da)
Exposure Time (minutes)
0.2
1
10
100
1000
7000

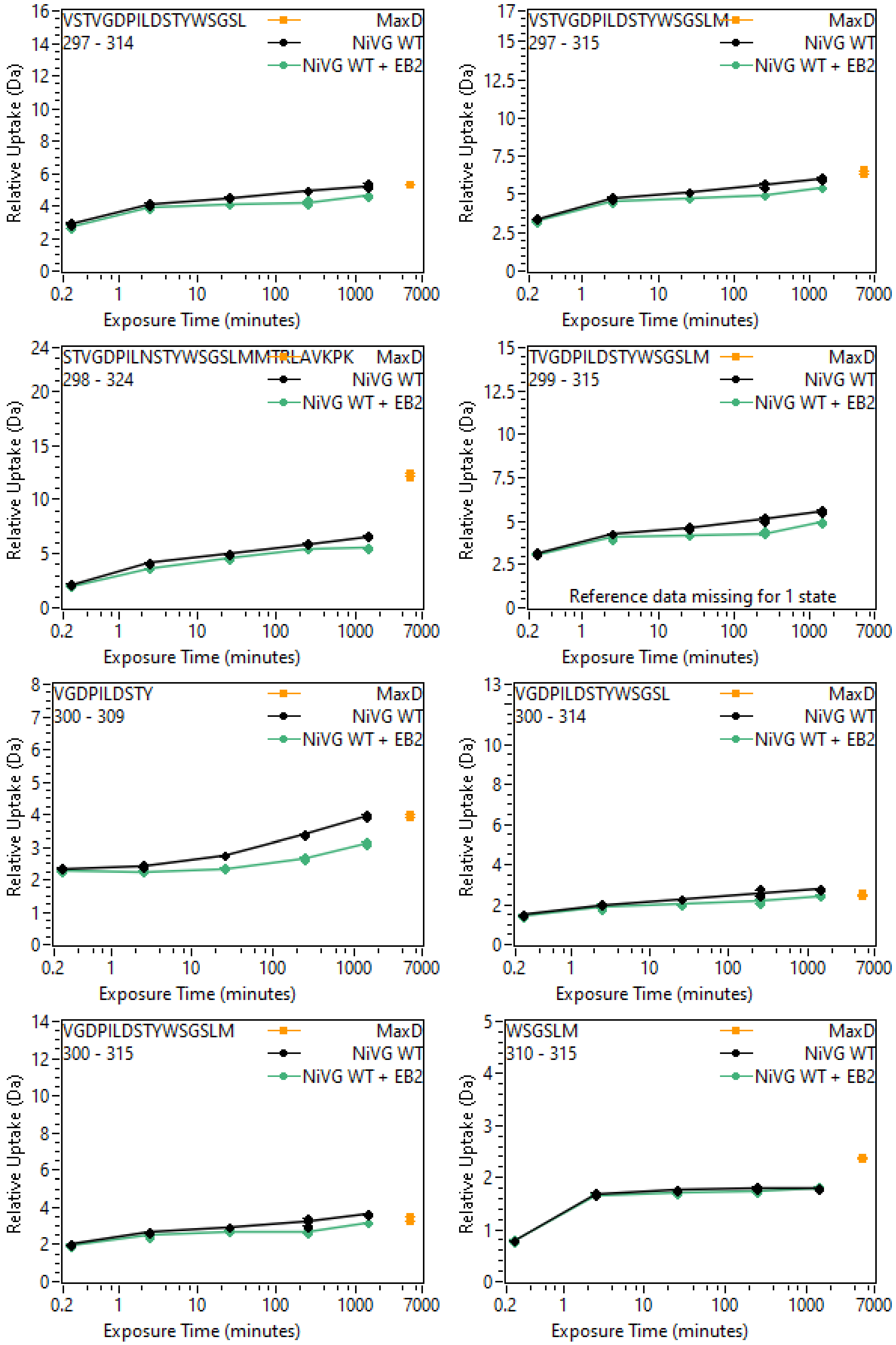
VSTVGDPILDSTYWSGSL
297 - 314
VSTVGDPILDSTYWSGSLM
297 - 315
STVGDPILNSTYWSGSLMMTRLAVKPK
298 - 324
TVGDPILDSTYWSGSLM
299 - 315
Reference data missing for 1 state
VGDPILDSTY
300 - 309
VGDPILDSTYWSGSL
300 - 314
VGDPILDSTYWSGSLM
300 - 315
WSGSLM
310 - 315
MaxD
NiVG WT
NiVG WT + EB2
Relative Uptake (Da)
Exposure Time (minutes)

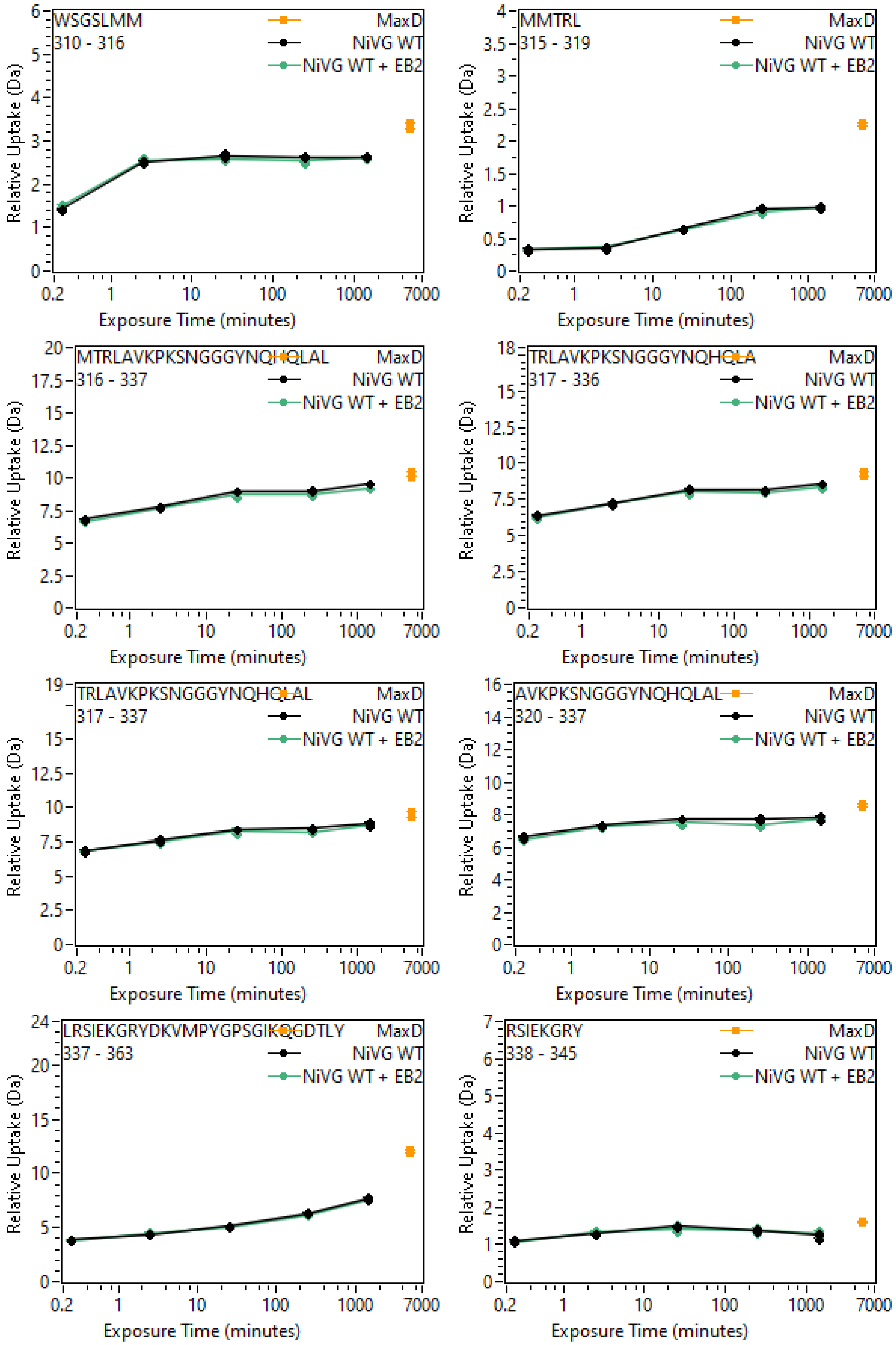
WSGSLMM
310 - 316
MMTRL
315 - 319
MTRLAVKPKSNGGGYNQHQLAL
316 - 337
TRLAVKPKSNGGGYNQHQLA
317 - 336
TRLAVKPKSNGGGYNQHQLAL
317 - 337
AVKPKSNGGGYNQHQLAL
320 - 337
LRSIEKGRYDKVMPYGPSGIKQGDTLY
337 - 363
RSIEKGRY
338 - 345
MaxD
NiVG WT
NiVG WT + EB2
Relative Uptake (Da)
Exposure Time (minutes)
0.2
1
10
100
1000
7000

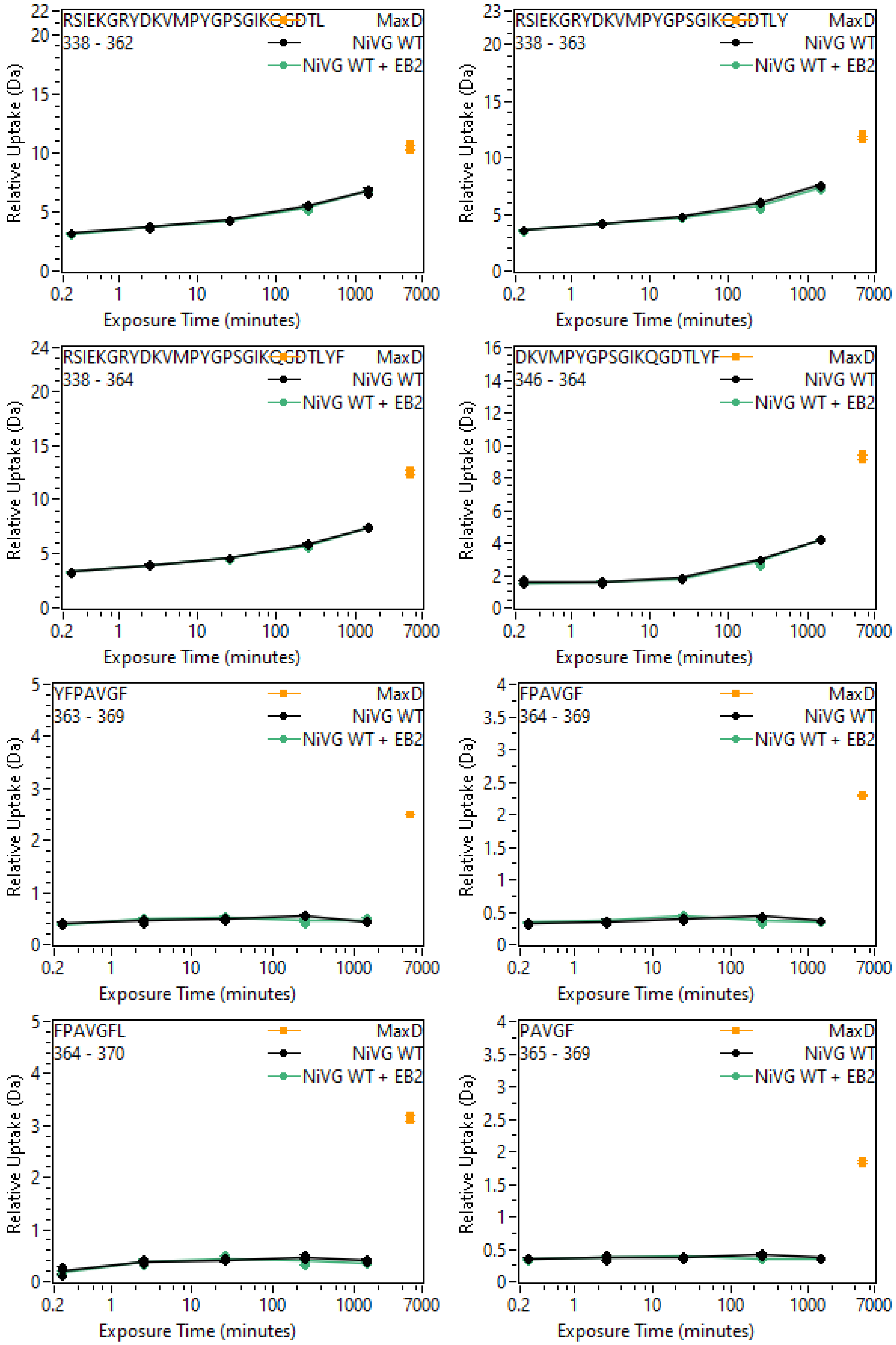
RSIEKGRYDKVMPYGPSGIKQGDTL
338 - 362
RSIEKGRYDKVMPYGPSGIKQGDTLY
338 - 363
RSIEKGRYDKVMPYGPSGIKQGDTLYF
338 - 364
DKVMPYGPSGIKQGDTLYF
346 - 364
YFPAVGF
363 - 369
FPAVGF
364 - 369
FPAVGFL
364 - 370
PAVGF
365 - 369
MaxD
NiVG WT
NiVG WT + EB2
Relative Uptake (Da)
Exposure Time (minutes)
0.2
1
10
100
1000
7000

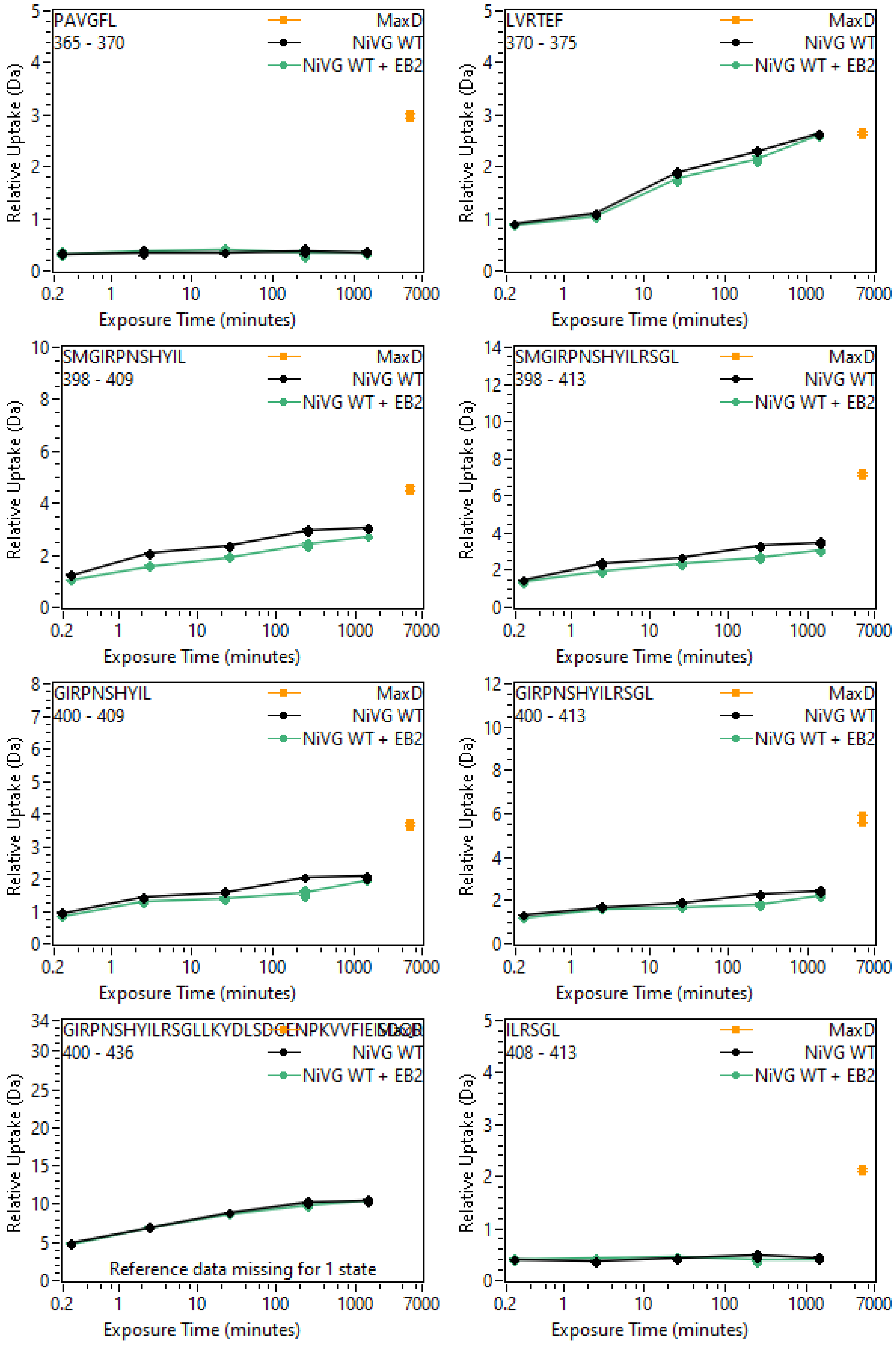
PAVGFL
365 - 370
MaxD
NiVG WT
NiVG WT + EB2
Relative Uptake (Da)
Exposure Time (minutes)
LVRTEF
370 - 375
MaxD
NiVG WT
NiVG WT + EB2
Relative Uptake (Da)
Exposure Time (minutes)
SMGIRPNSHYIL
398 - 409
MaxD
NiVG WT
NiVG WT + EB2
Relative Uptake (Da)
Exposure Time (minutes)
SMGIRPNSHYILRSGL
398 - 413
MaxD
NiVG WT
NiVG WT + EB2
Relative Uptake (Da)
Exposure Time (minutes)
GIRPNSHYIL
400 - 409
MaxD
NiVG WT
NiVG WT + EB2
Relative Uptake (Da)
Exposure Time (minutes)
GIRPNSHYILRSGL
400 - 413
MaxD
NiVG WT
NiVG WT + EB2
Relative Uptake (Da)
Exposure Time (minutes)
400 - 436
NiVG WT
NiVG WT + EB2
Reference data missing for 1 state
Relative Uptake (Da)
Exposure Time (minutes)
ILRSGL
408 - 413
MaxD
NiVG WT
NiVG WT + EB2
Relative Uptake (Da)
Exposure Time (minutes)

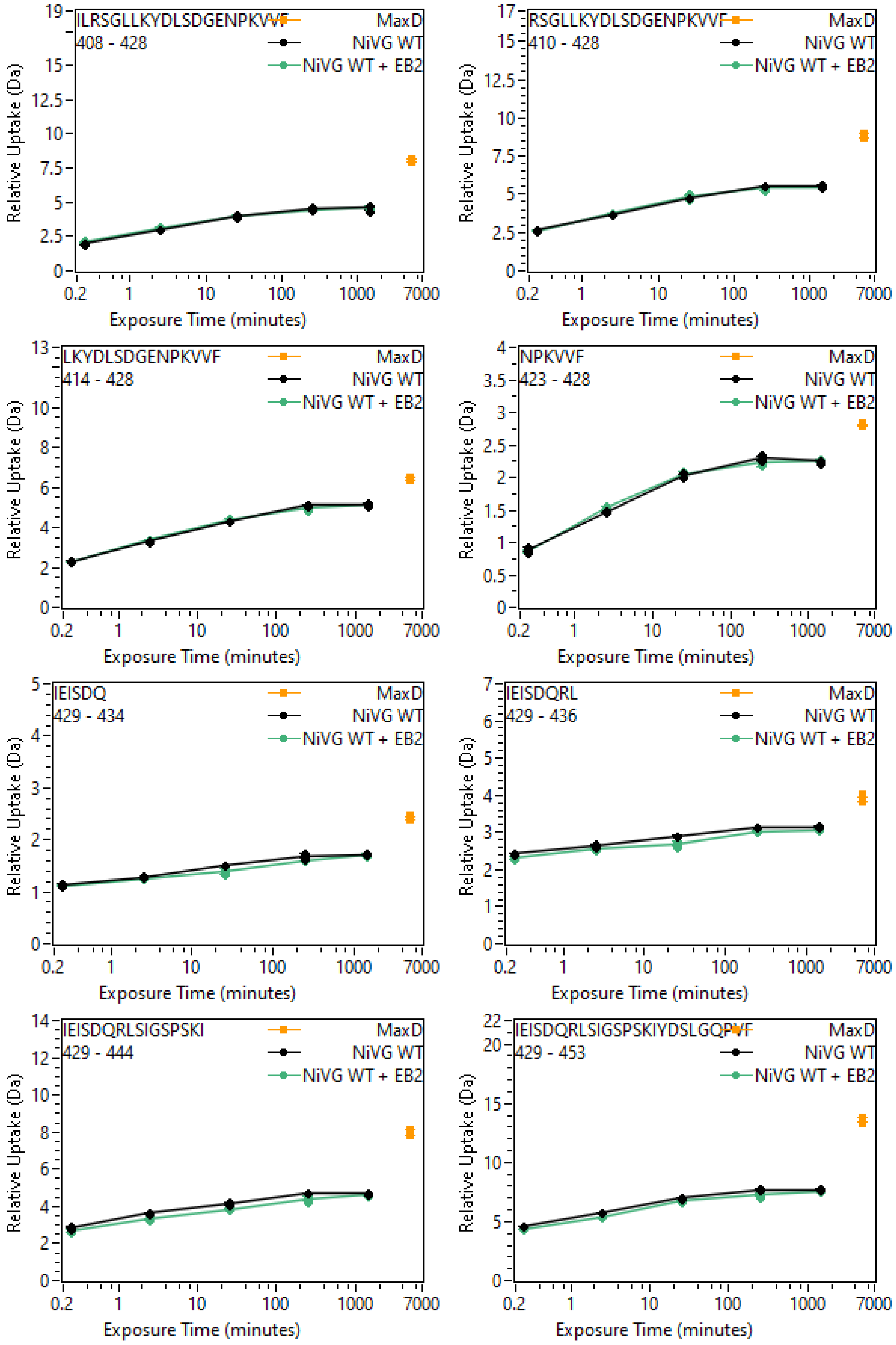
ILRSGLLKYDLSDGENPKVVF
408 - 428
RSGLLKYDLSDGENPKVVF
410 - 428
LKYDLSDGENPKVVF
414 - 428
NPKVVF
423 - 428
IEISDQ
429 - 434
IEISDQRL
429 - 436
IEISDQRLSIGSPSKI
429 - 444
IEISDQRLSIGSPSKIYDSLGQPVF
429 - 453
MaxD
NiVG WT
NiVG WT + EB2
Relative Uptake (Da)
Exposure Time (minutes)
0.2
1
10
100
1000
7000

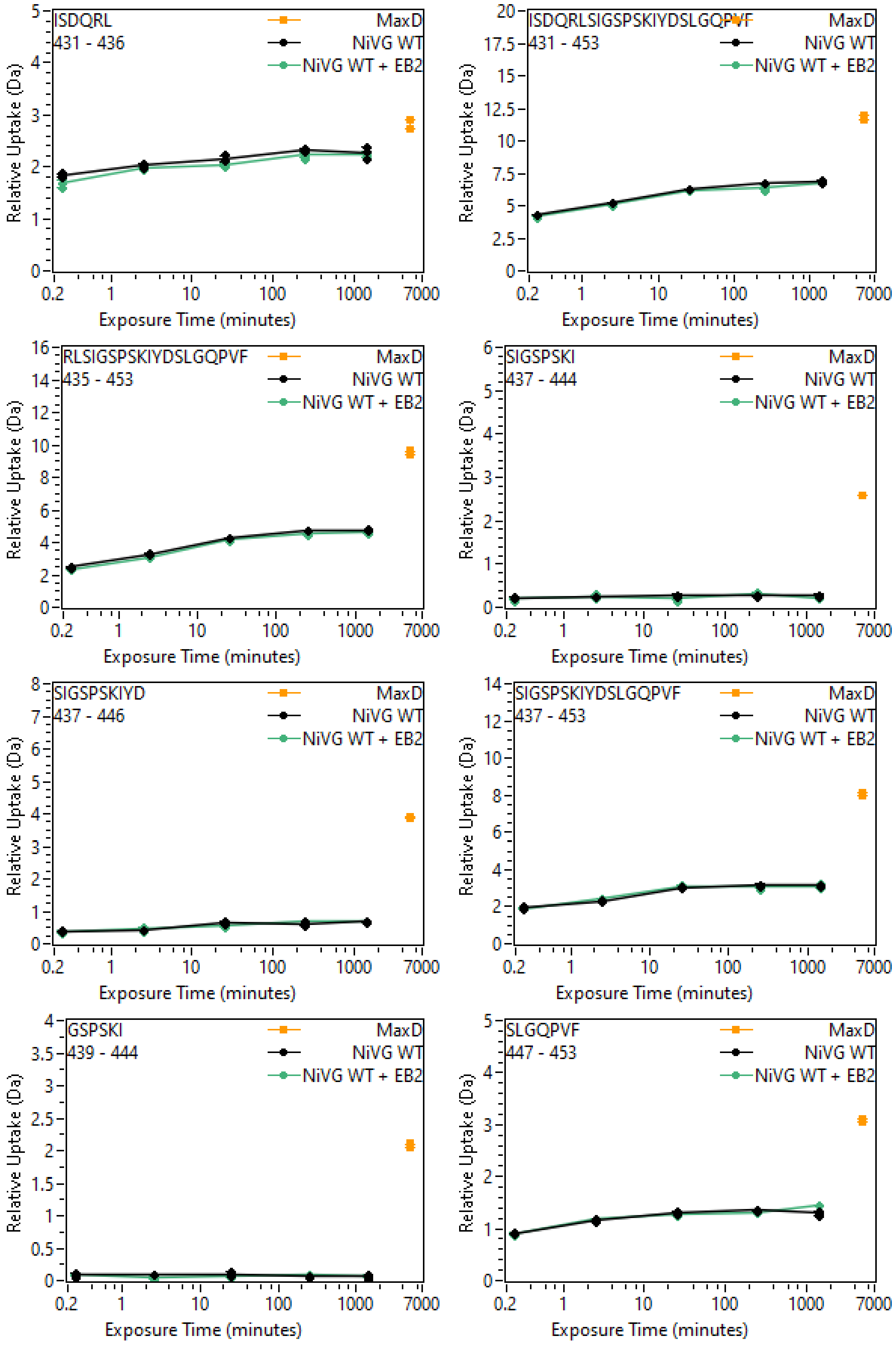
ISDQRL
431 - 436
MaxD
NiVG WT
NiVG WT + EB2
ISDQRLSIGSPSKIYDSLGQPVF
431 - 453
RLSIGSPSKIYDSLGQPVF
435 - 453
SIGSPSKI
437 - 444
SIGSPSKIYD
437 - 446
SIGSPSKIYDSLGQPVF
437 - 453
GSPSKI
439 - 444
SLGQPVF
447 - 453
Relative Uptake (Da)
Exposure Time (minutes)
0.2
1
10
100
1000
7000

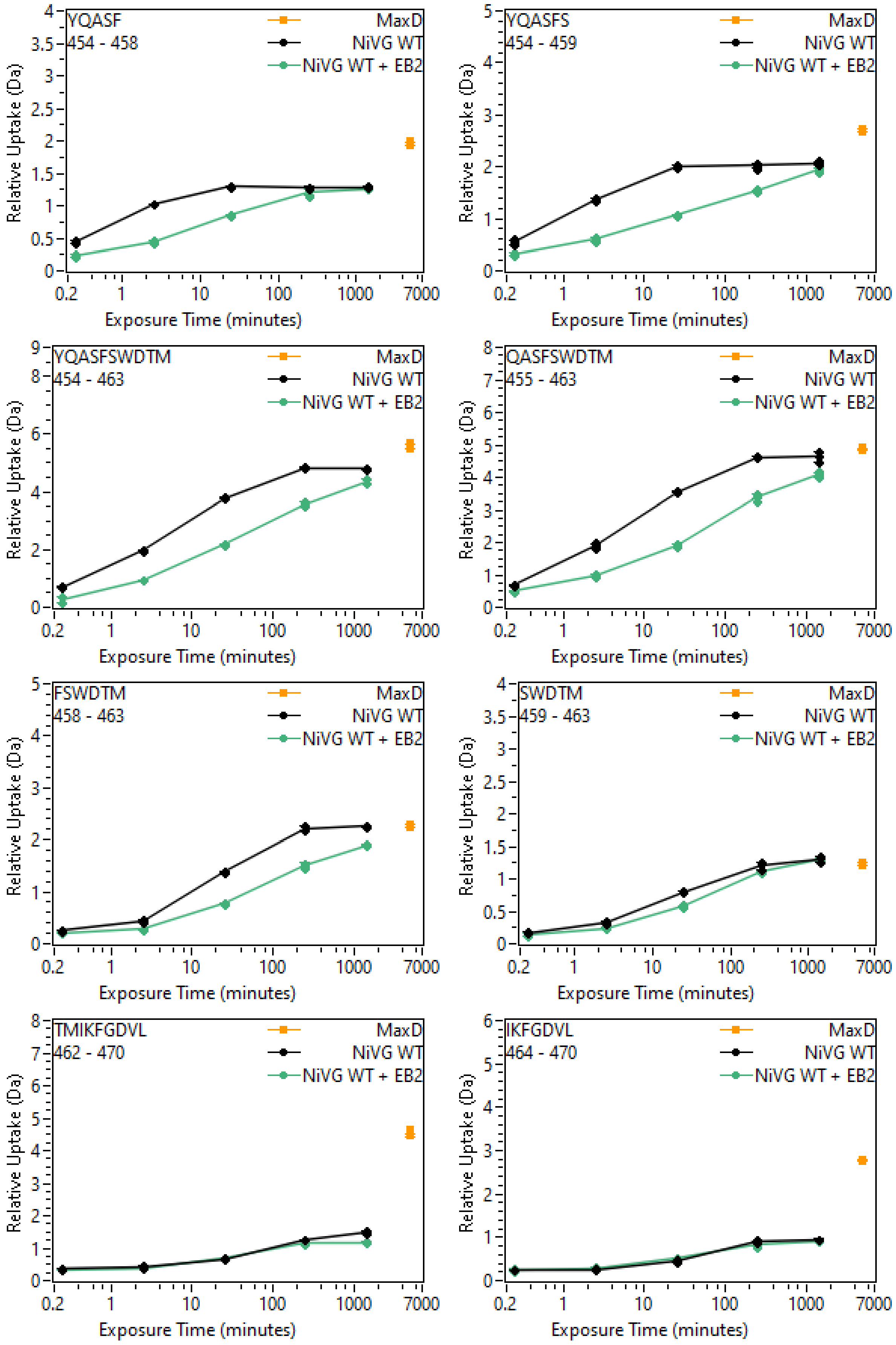
YQASF
454 - 458
YQASFS
454 - 459
YQASFSWDTM
454 - 463
QASFSWDTM
455 - 463
FSWDTM
458 - 463
SWDTM
459 - 463
TMIKFGDVL
462 - 470
IKFGDVL
464 - 470
MaxD
NiVG WT
NiVG WT + EB2
Relative Uptake (Da)
Exposure Time (minutes)
0.2
1
10
100
1000
7000

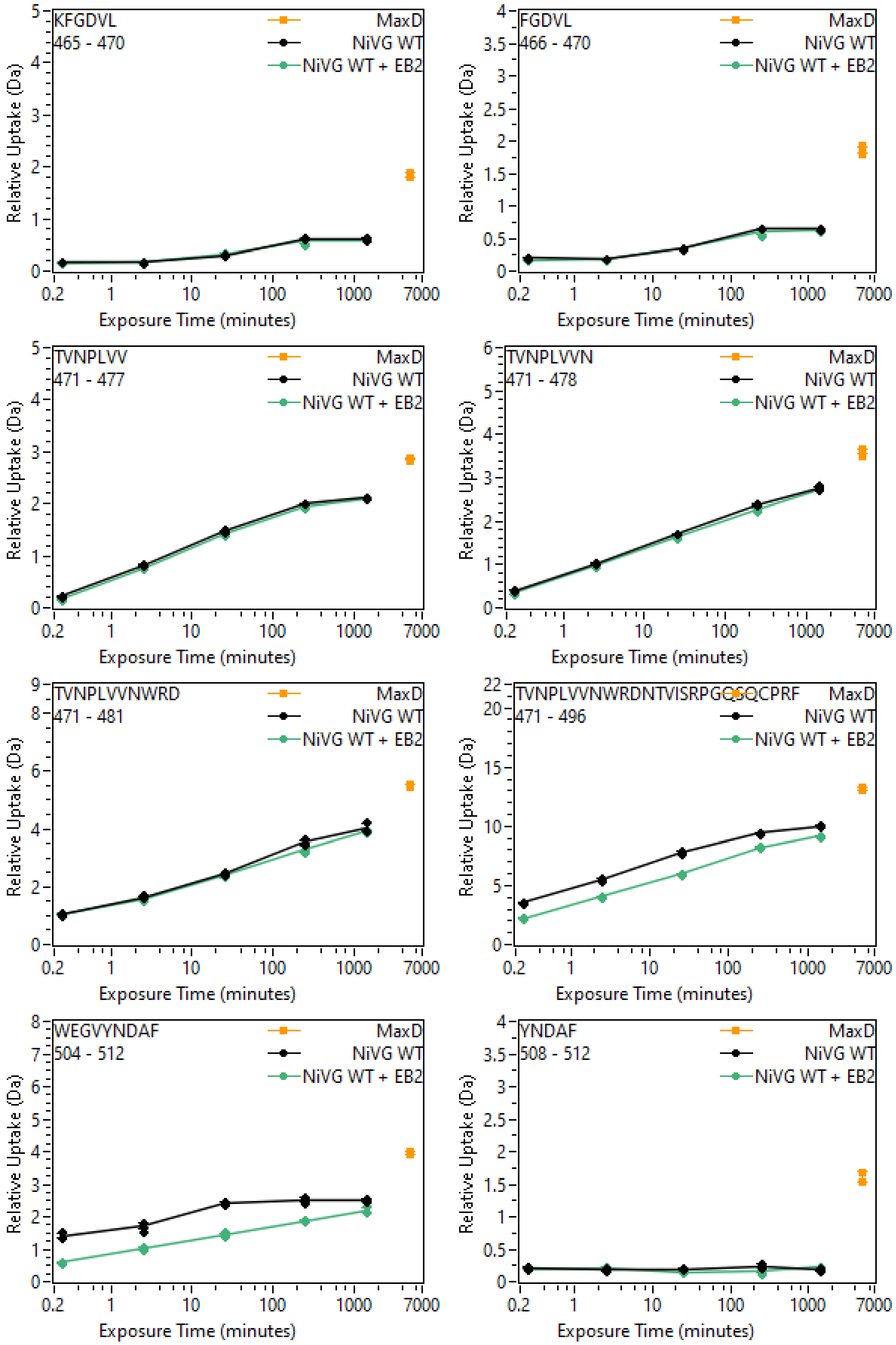
KFGDVL
465 - 470
FGDVL
466 - 470
TVNPLVV
471 - 477
TVNPLVVN
471 - 478
TVNPLVVNWRD
471 - 481
471 - 496
WEGVYNDAF
504 - 512
YNDAF
508 - 512
MaxD
NiVG WT
NiVG WT + EB2
Relative Uptake (Da)
Exposure Time (minutes)

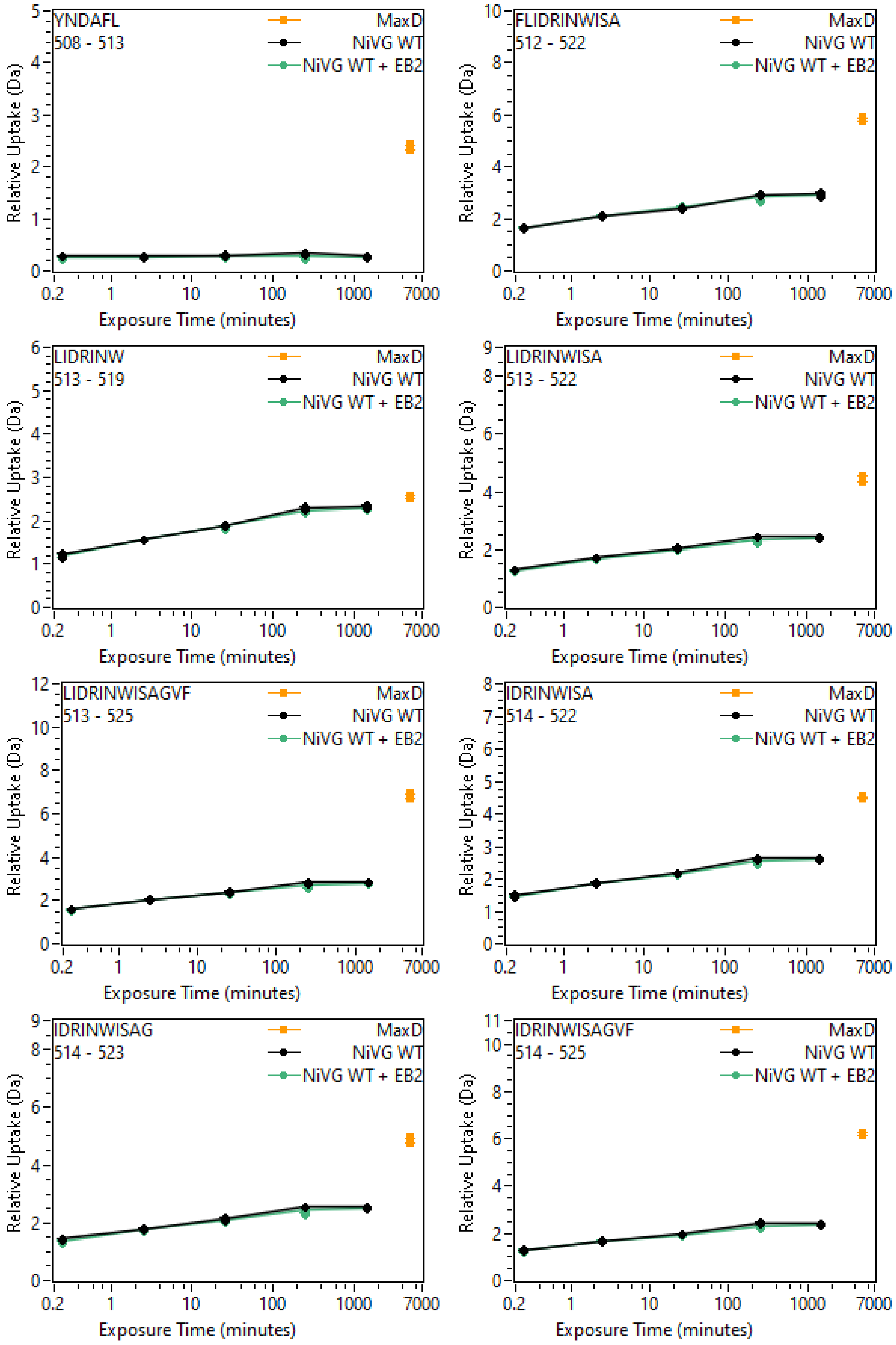
YNDAFL
508 - 513
FLIDRINWISA
512 - 522
LIDRINW
513 - 519
LIDRINWISA
513 - 522
LIDRINWISAGVF
513 - 525
IDRINWISA
514 - 522
IDRINWISAG
514 - 523
IDRINWISAGVF
514 - 525
MaxD
NiVG WT
NiVG WT + EB2
Relative Uptake (Da)
Exposure Time (minutes)
0.2
1
10
100
1000
7000

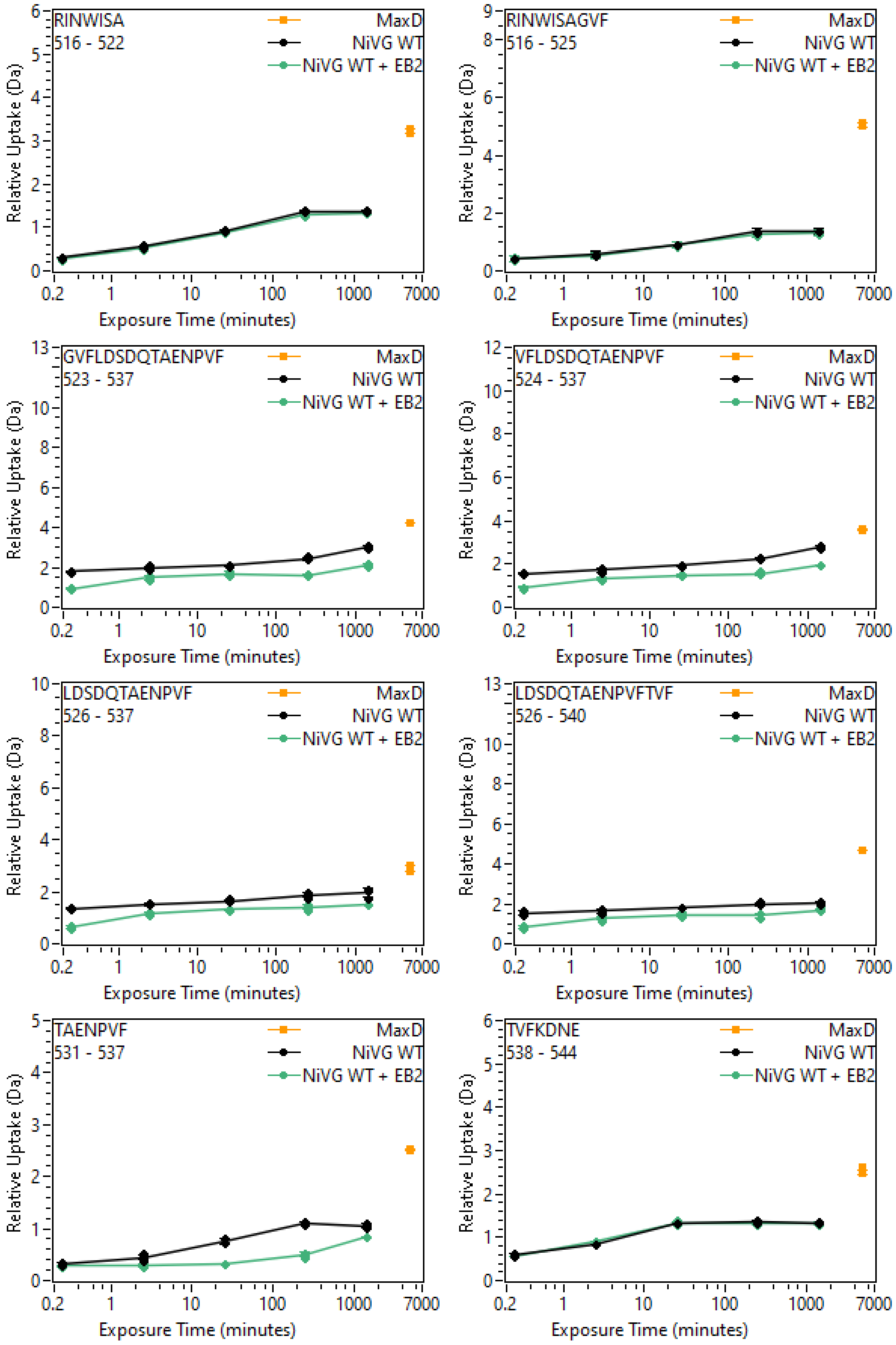
RINWISA
516 - 522
RINWISAGVF
516 - 525
GVFLDSDQTAENPVF
523 - 537
VFLDSDQTAENPVF
524 - 537
LDSDQTAENPVF
526 - 537
LDSDQTAENPVFTVF
526 - 540
TAENPVF
531 - 537
TVFKDNE
538 - 544
MaxD
NiVG WT
NiVG WT + EB2
Relative Uptake (Da)
Exposure Time (minutes)
0.2
1
10
100
1000
7000

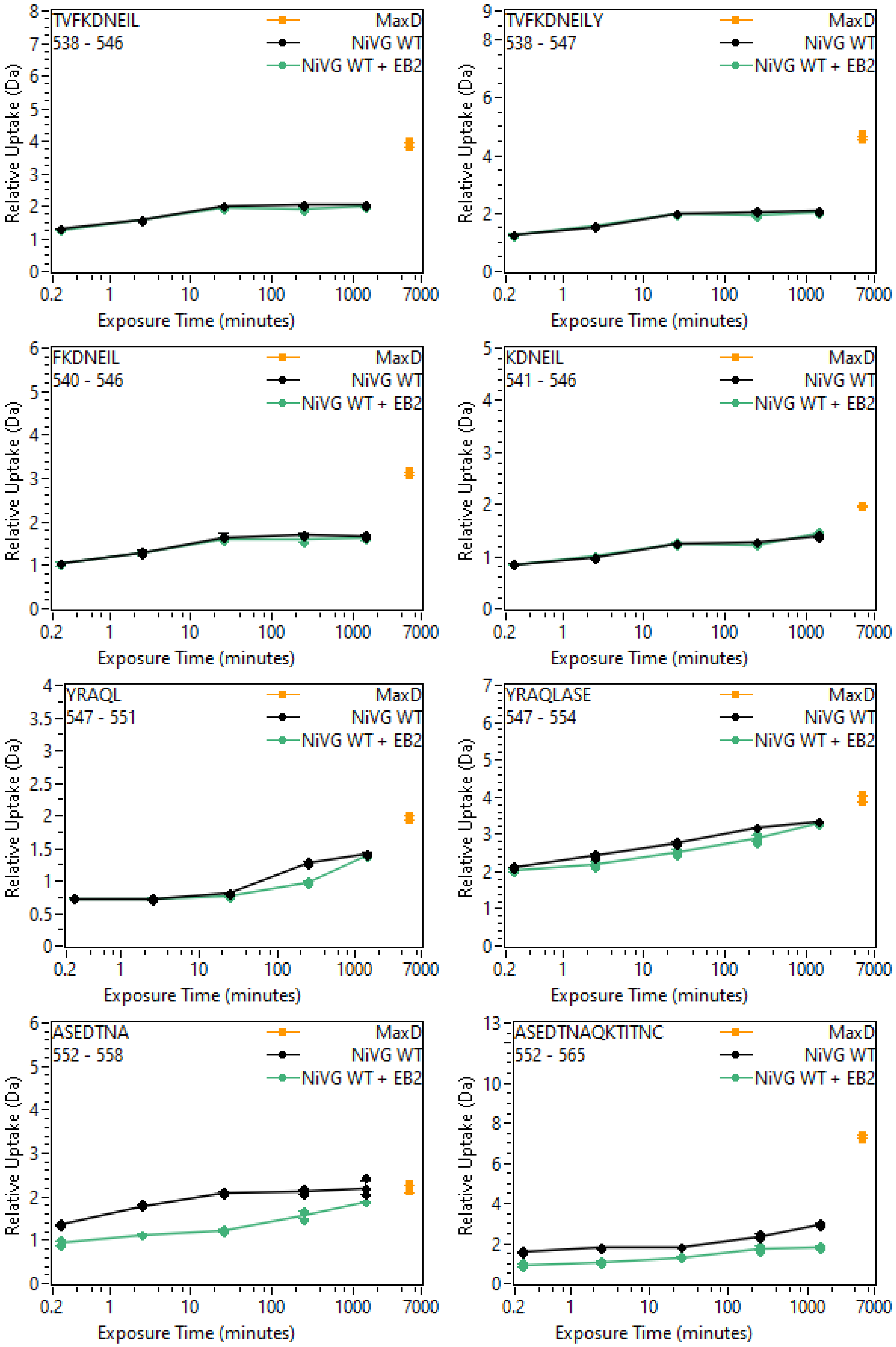
TVFKDNEIL
538 - 546
MaxD
NiVG WT
NiVG WT + EB2
TVFKDNEILY
538 - 547
FKDNEIL
540 - 546
KDNEIL
541 - 546
YRAQL
547 - 551
YRAQLASE
547 - 554
ASEDTNA
552 - 558
ASEDTNAQKTITNC
552 - 565
Relative Uptake (Da)
Exposure Time (minutes)
0.2
1
10
100
1000
7000

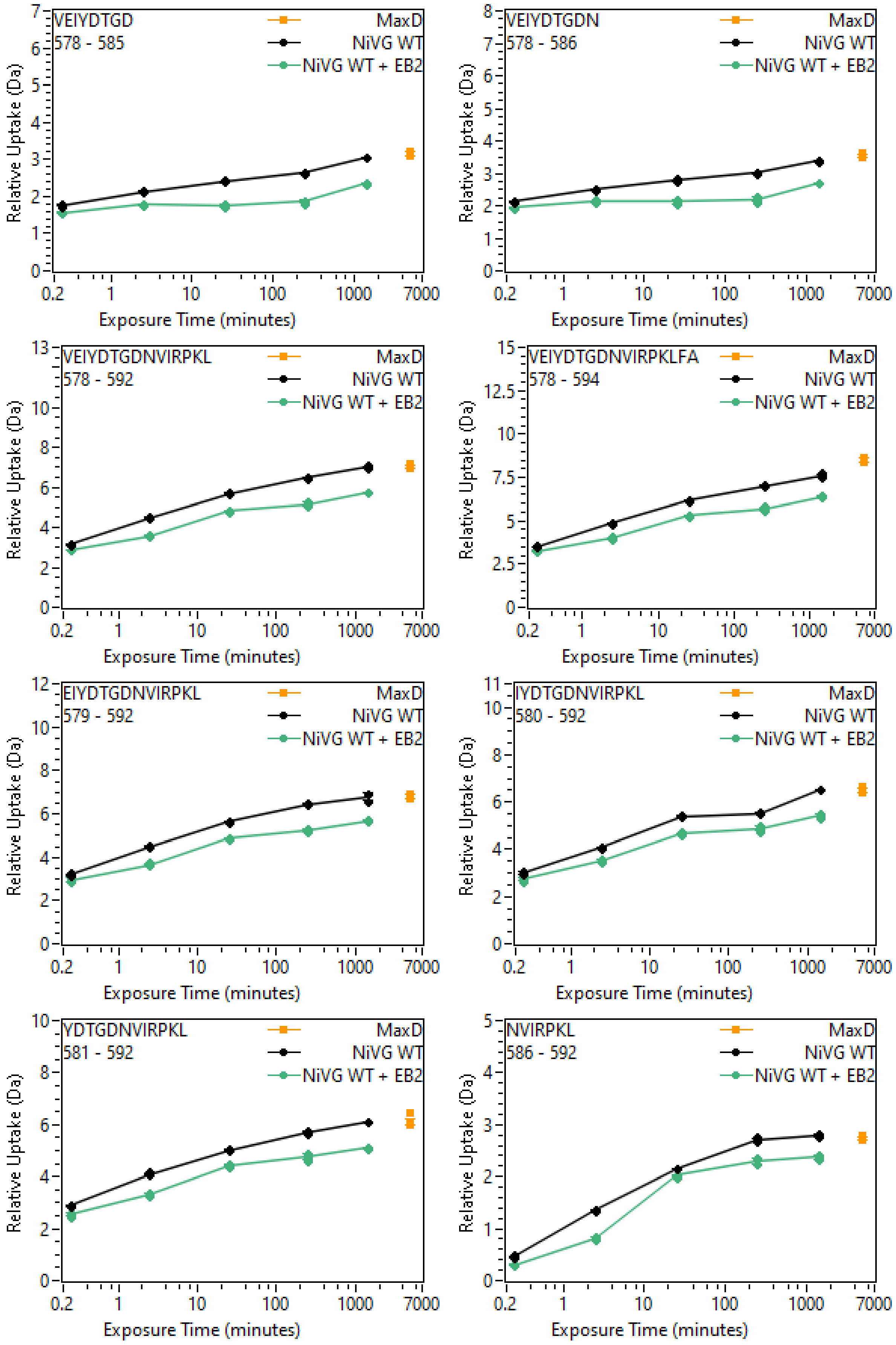
VEIYDTGD
578 - 585
VEIYDTGDN
578 - 586
VEIYDTGDNVIRPKL
578 - 592
VEIYDTGDNVIRPKLFA
578 - 594
EIYDTGDNVIRPKL
579 - 592
IYDTGDNVIRPKL
580 - 592
YDTGDNVIRPKL
581 - 592
NVIRPKL
586 - 592
MaxD
NiVG WT
NiVG WT + EB2
Relative Uptake (Da)
Exposure Time (minutes)
0.2
1
10
100
1000
7000

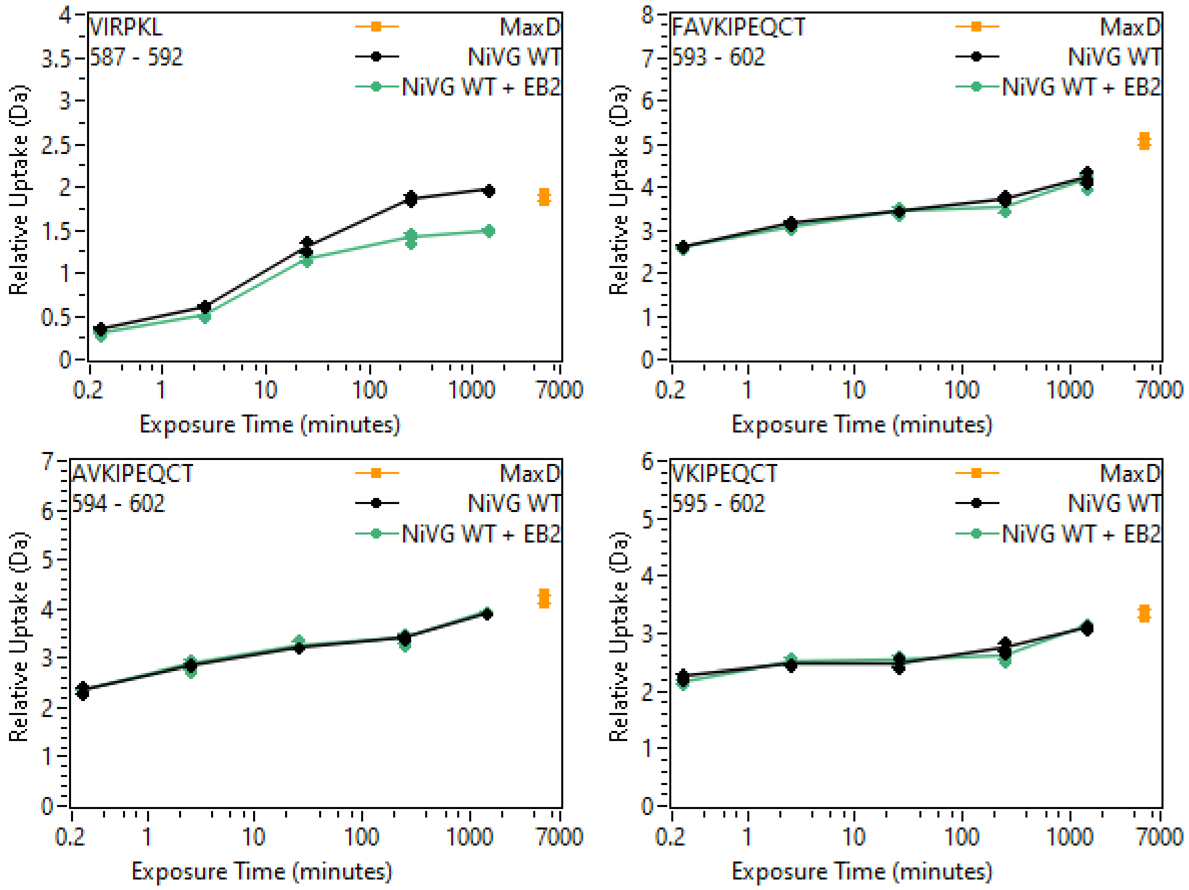
VIRPKL
587 - 592
FAVKIPEQCT
593 - 602
AVKIPEQCT
594 - 602
VKIPEQCT
595 - 602
MaxD
NiVG WT
NiVG WT + EB2
Relative Uptake (Da)
Exposure Time (minutes)
0.2
1
10
100
1000
7000

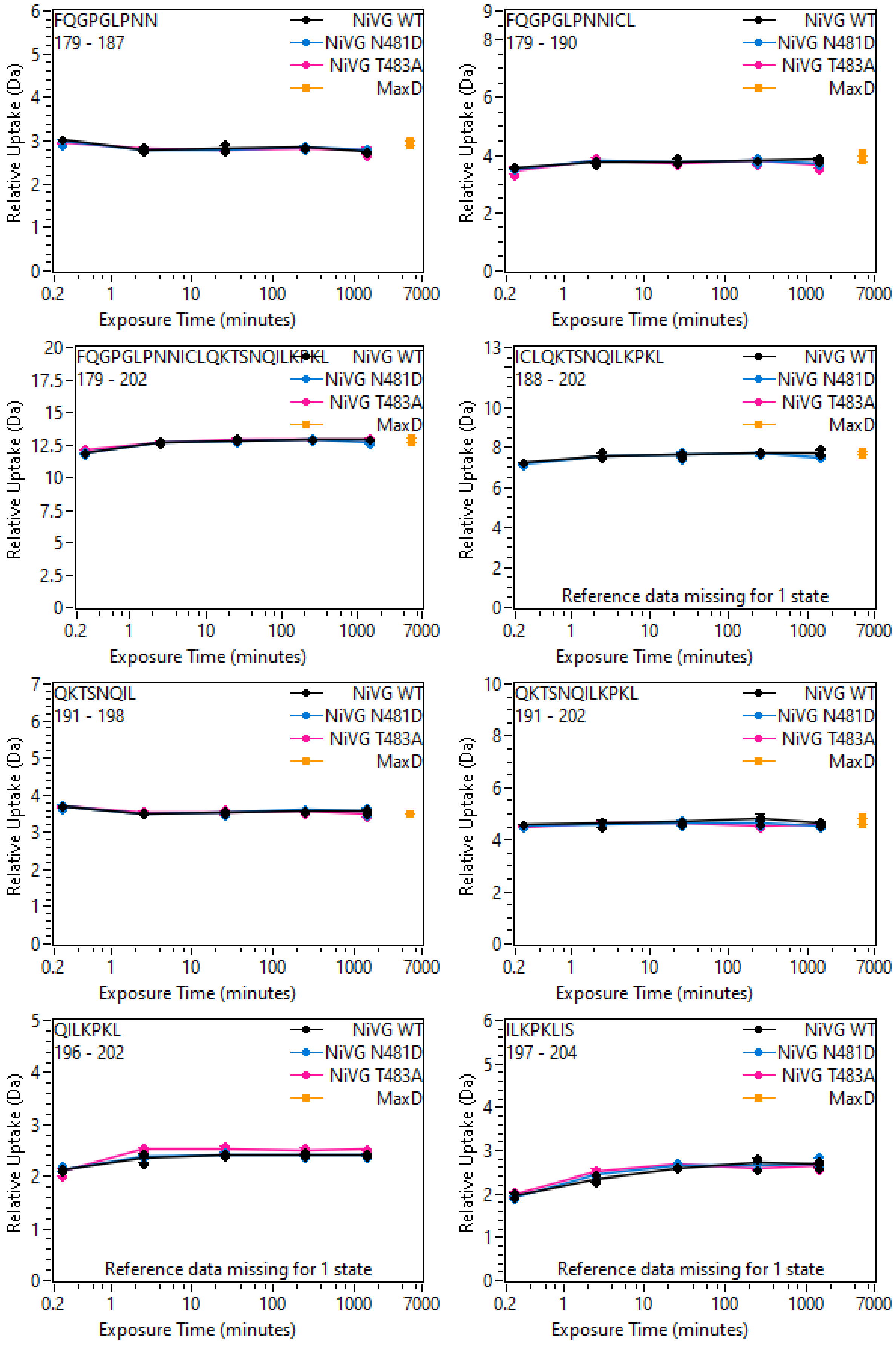
FQGPGLPNN
179 - 187
FQGPGLPNNICL
179 - 190
FQGPGLPNNICLQKTSNQILKPKL
179 - 202
ICLQKTSNQILKPKL
188 - 202
Reference data missing for 1 state
QKTSNQIL
191 - 198
QKTSNQILKPKL
191 - 202
QILKPKL
196 - 202
Reference data missing for 1 state
ILKPKLIS
197 - 204
Reference data missing for 1 state
NiVG WT
NiVG N481D
NiVG T483A
MaxD
Relative Uptake (Da)
Exposure Time (minutes)
0.2
1
10
100
1000
7000

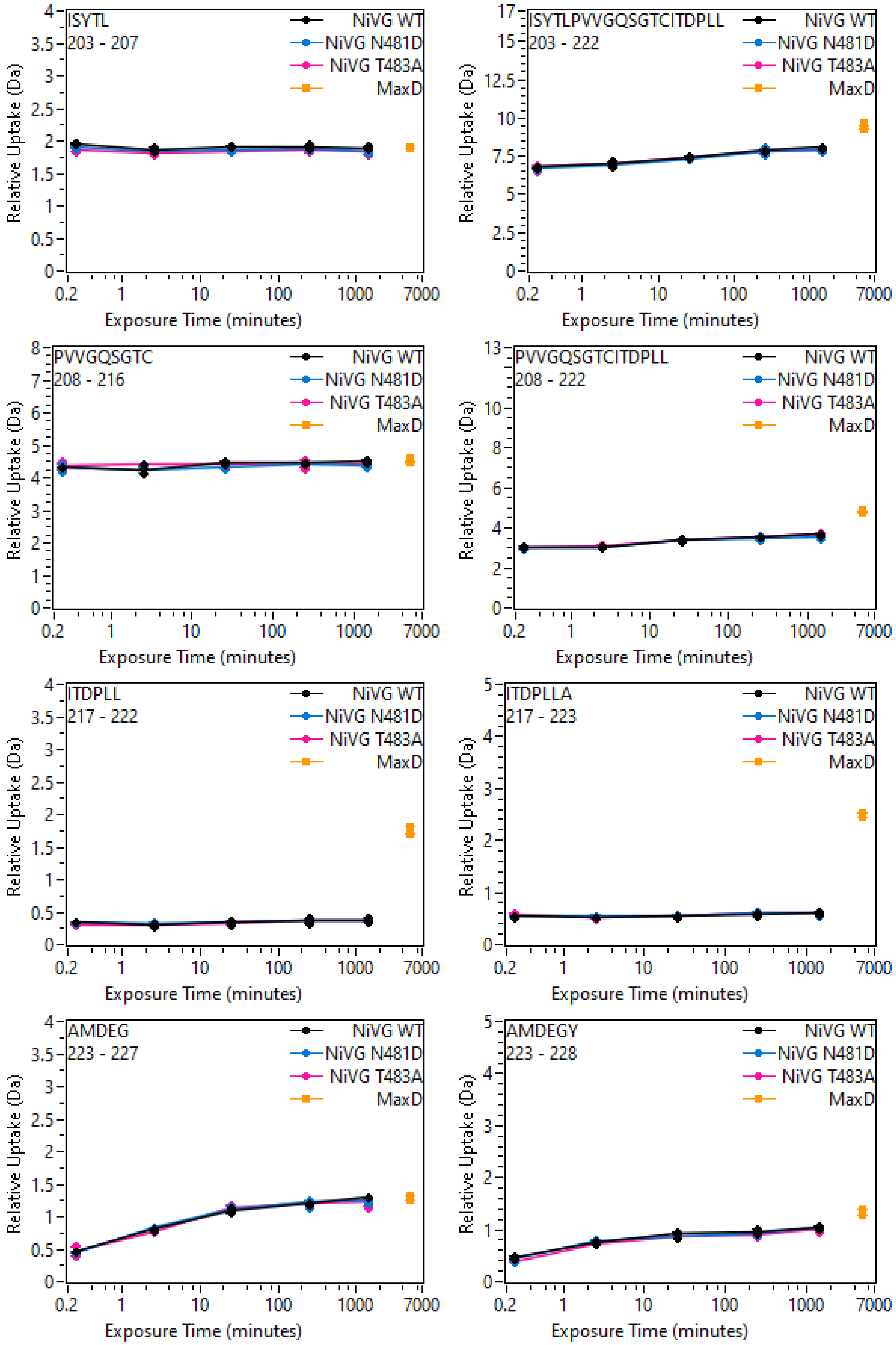
ISYTL
203 - 207
ISYTLPVVGQSGTCITDPLL
203 - 222
PVVGQSGTC
208 - 216
PVVGQSGTCITDPLL
208 - 222
ITDPLL
217 - 222
ITDPLLA
217 - 223
AMDEG
223 - 227
AMDEGY
223 - 228
NiVG WT
NiVG N481D
NiVG T483A
MaxD
Relative Uptake (Da)
Exposure Time (minutes)
0.2
1
10
100
1000
7000

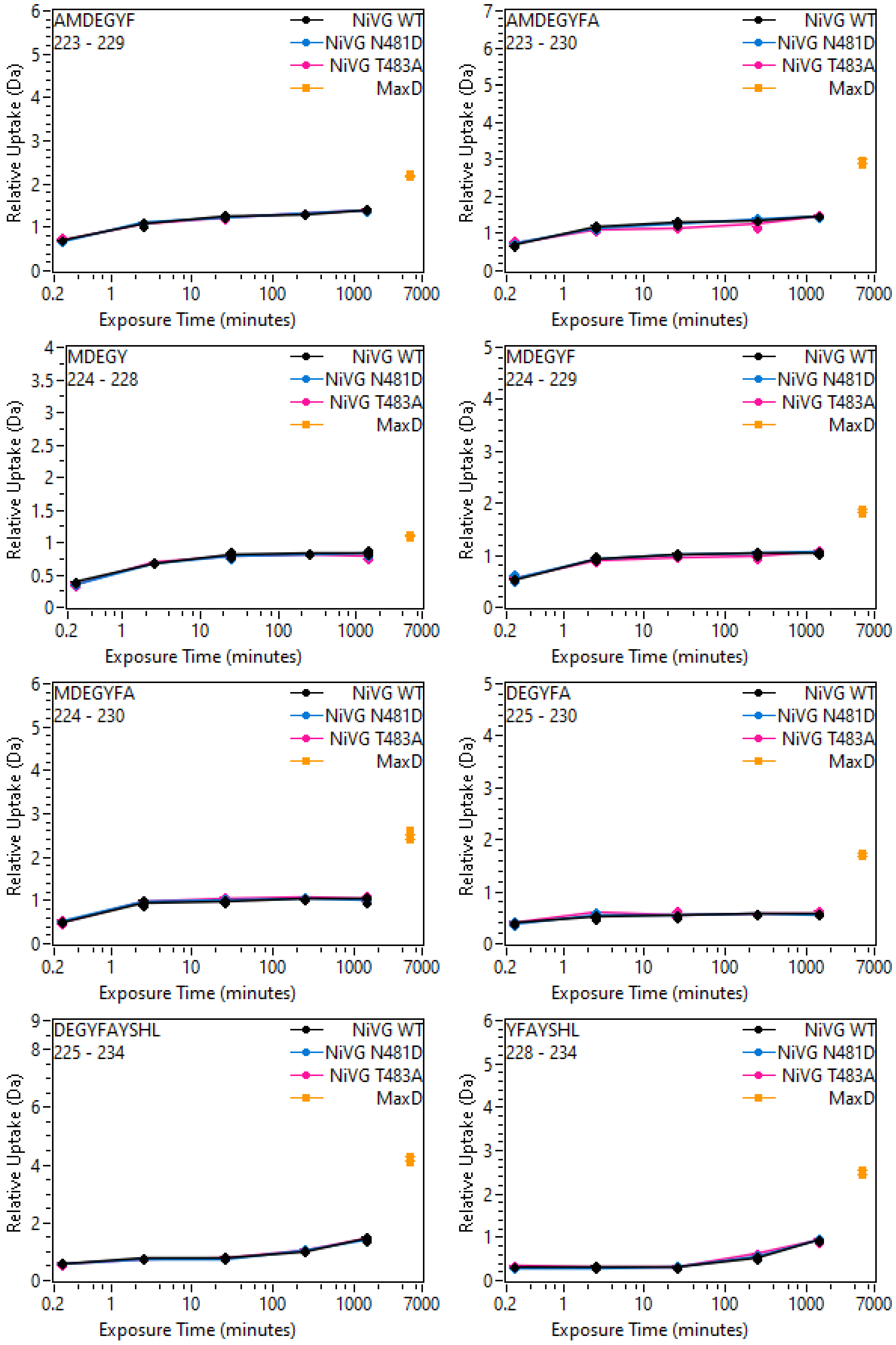
AMDEGYF
223 - 229
AMDEGYFA
223 - 230
MDEGY
224 - 228
MDEGYF
224 - 229
MDEGYFA
224 - 230
DEGYFA
225 - 230
DEGYFAYSHL
225 - 234
YFAYSHL
228 - 234
NiVG WT
NiVG N481D
NiVG T483A
MaxD
Relative Uptake (Da)
Exposure Time (minutes)
0.2
1
10
100
1000
7000

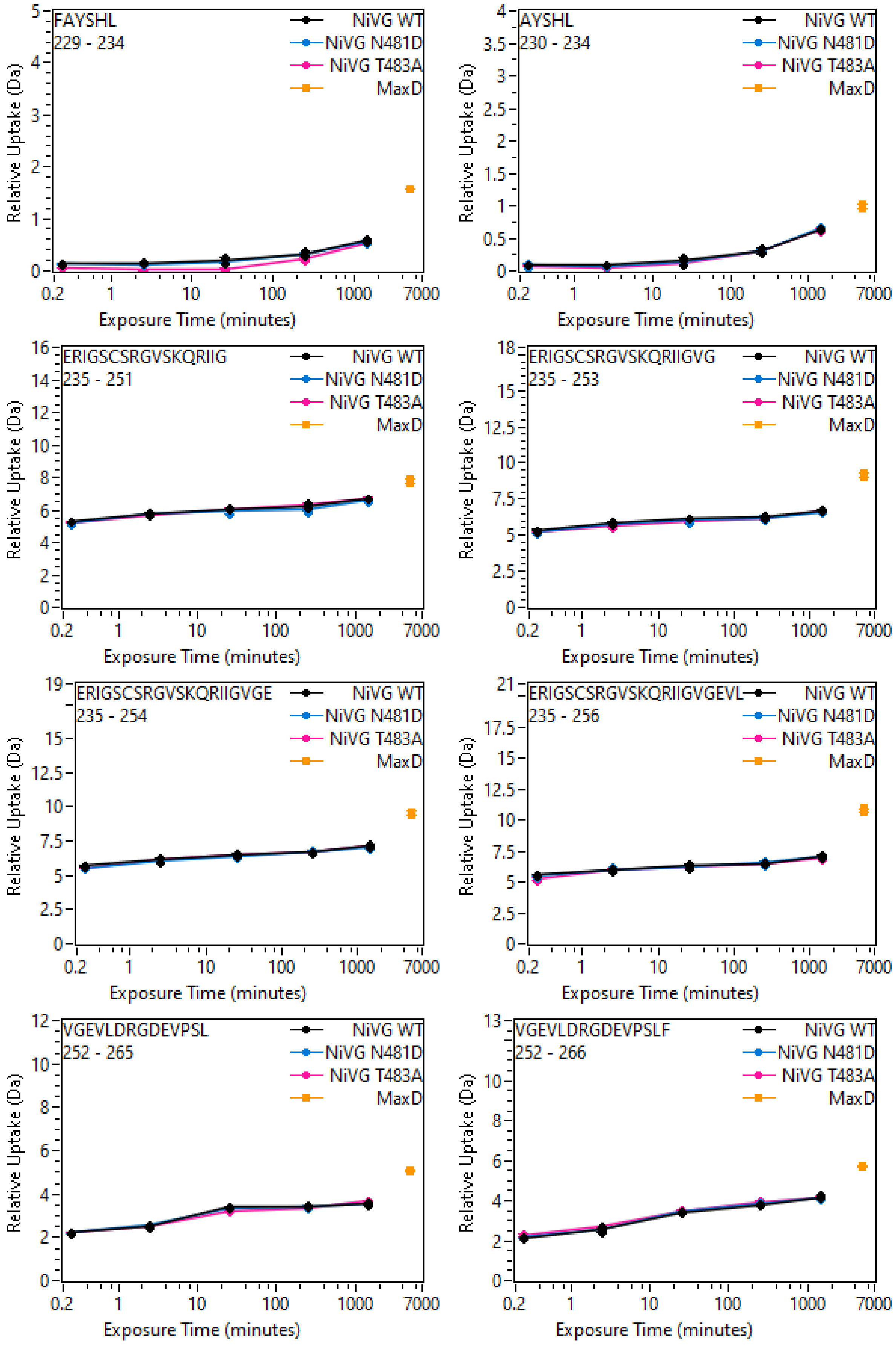
FAYSHL
229 - 234
AYSHL
230 - 234
ERIGSCSRGVSKQRIIG
235 - 251
ERIGSCSRGVSKQRIIGVG
235 - 253
ERIGSCSRGVSKQRIIGVGE
235 - 254
ERIGSCSRGVSKQRIIGVGEVL
235 - 256
VGEVLDRGDEVPSL
252 - 265
VGEVLDRGDEVPSLF
252 - 266
NiVG WT
NiVG N481D
NiVG T483A
MaxD
Relative Uptake (Da)
Exposure Time (minutes)
0.2
1
10
100
1000
7000

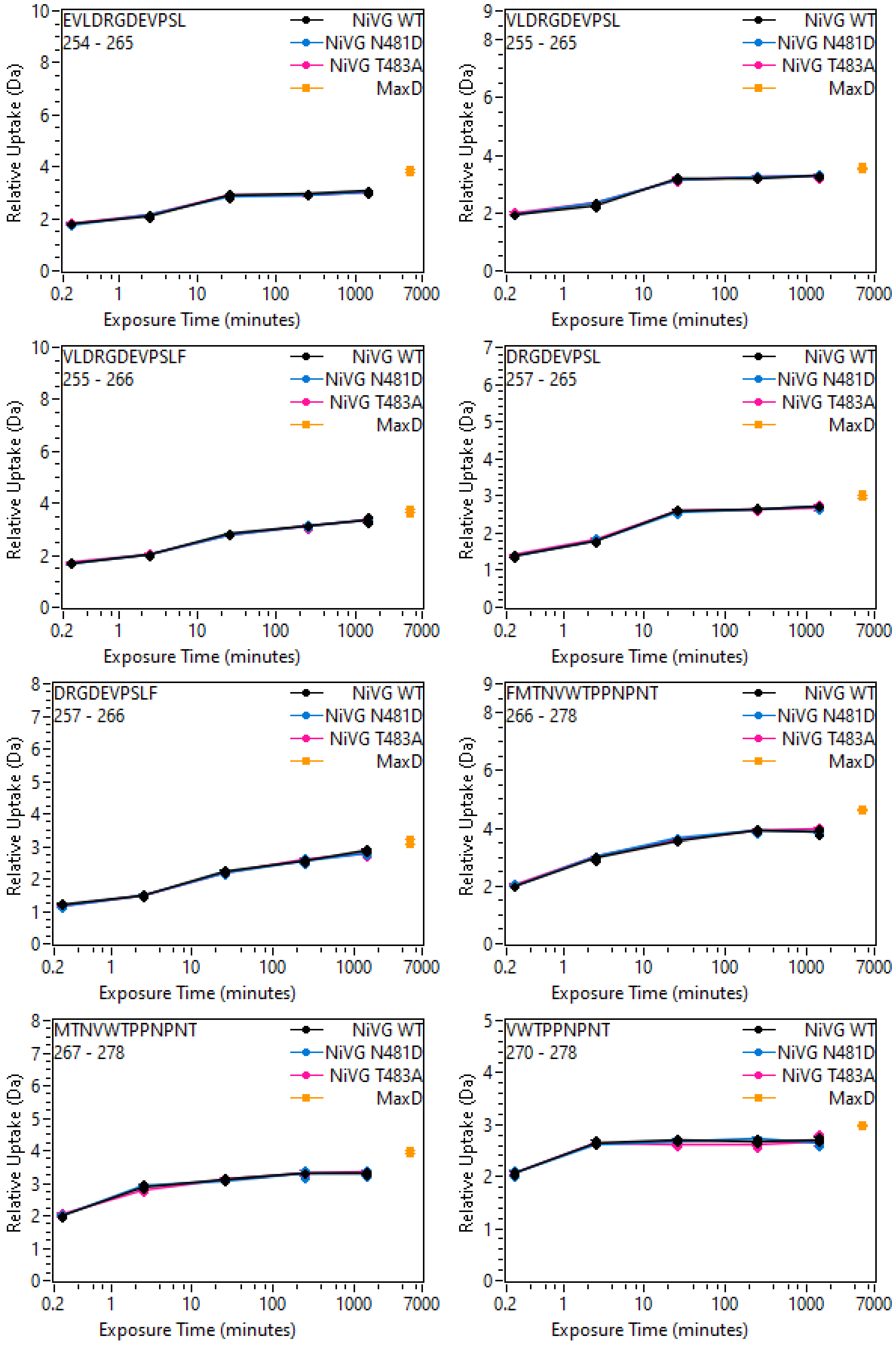
EVLDRGDEVPSL
254 - 265
VLDRGDEVPSL
255 - 265
VLDRGDEVPSLF
255 - 266
DRGDEVPSL
257 - 265
DRGDEVPSLF
257 - 266
FMTNVWTPPNPNT
266 - 278
MTNVWTPPNPNT
267 - 278
VWTPPNPNT
270 - 278
NiVG WT
NiVG N481D
NiVG T483A
MaxD
Relative Uptake (Da)
Exposure Time (minutes)
0.2
1
10
100
1000
7000

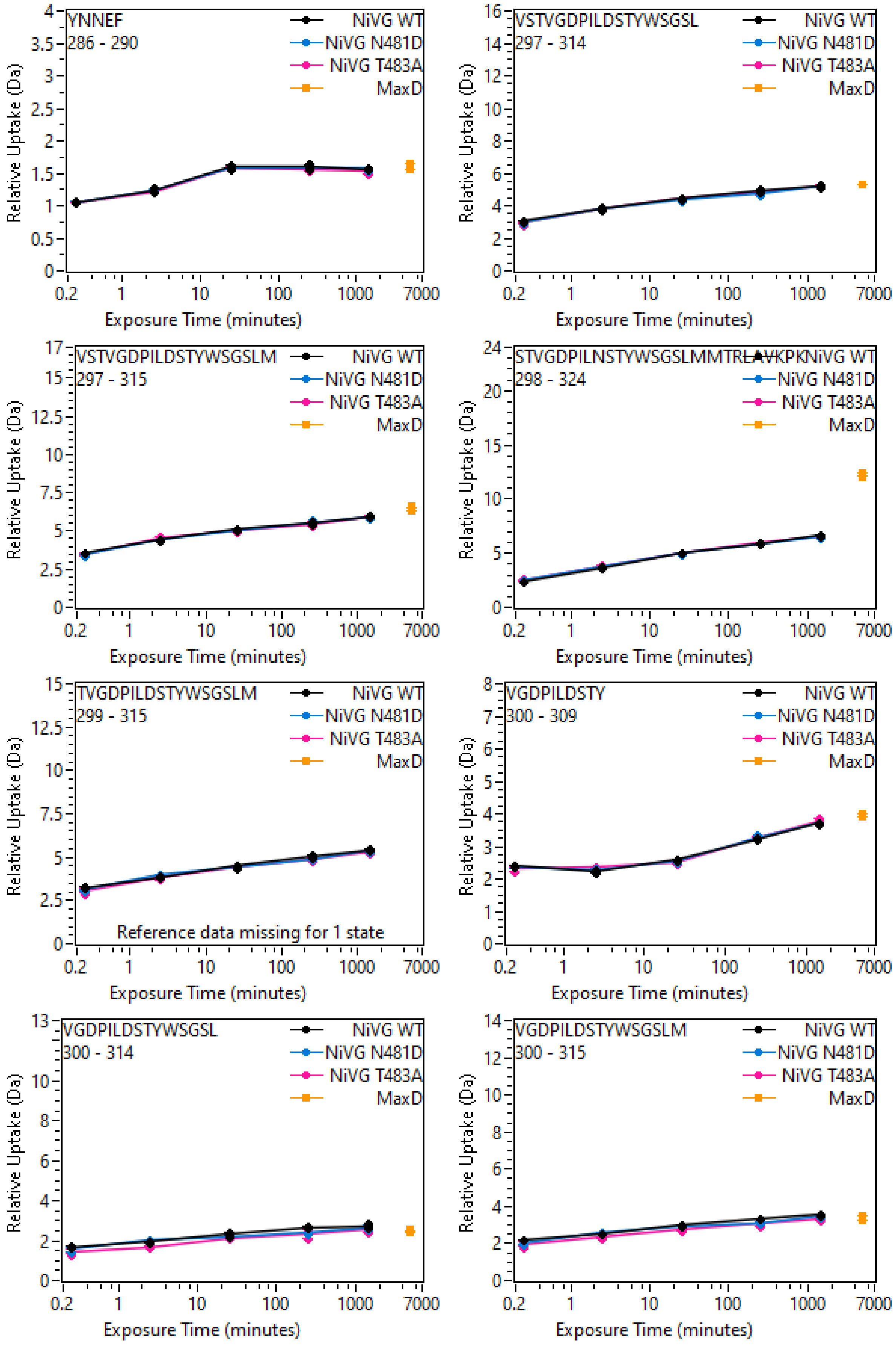
YNNEF
286 - 290
NiVG WT
NiVG N481D
NiVG T483A
MaxD
Relative Uptake (Da)
Exposure Time (minutes)
VSTVGDPILDSTYWSGSL
297 - 314
VSTVGDPILDSTYWSGSLM
297 - 315
STVGDPILNSTYWSGSLMMTRLAVKPK
298 - 324
TVGDPILDSTYWSGSLM
299 - 315
Reference data missing for 1 state
VGDPILDSTY
300 - 309
VGDPILDSTYWSGSL
300 - 314
VGDPILDSTYWSGSLM
300 - 315

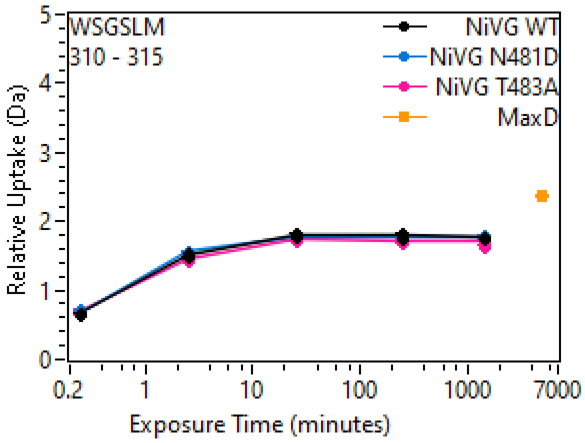
WSGSLM
310 - 315
NiVG WT
NiVG N481D
NiVG T483A
MaxD
Relative Uptake (Da)
Exposure Time (minutes)

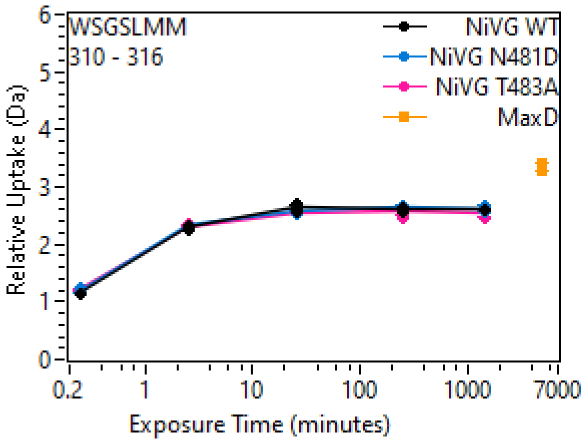
WSGSLMM
310 - 316
NiVG WT
NiVG N481D
NiVG T483A
MaxD
Relative Uptake (Da)
Exposure Time (minutes)

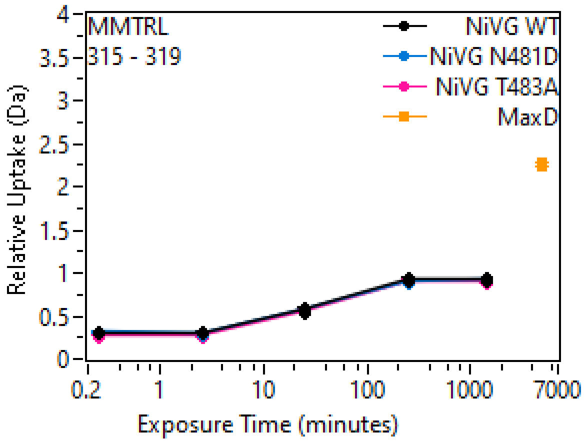
MMTRL
315 - 319
NiVG WT
NiVG N481D
NiVG T483A
MaxD
Relative Uptake (Da)
Exposure Time (minutes)

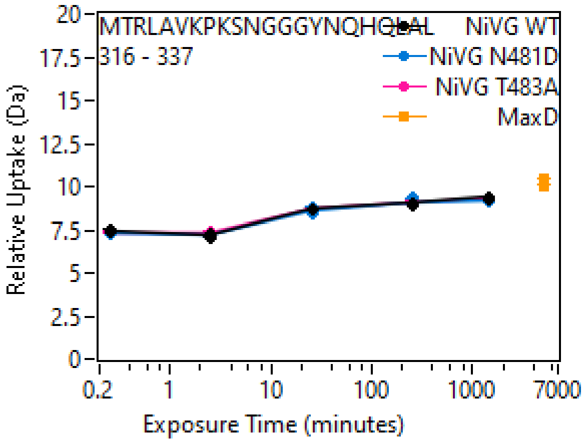
MTRLAVKPKSNGGGYNQHQLAL
316 - 337
NiVG WT
NiVG N481D
NiVG T483A
MaxD
Relative Uptake (Da)
Exposure Time (minutes)

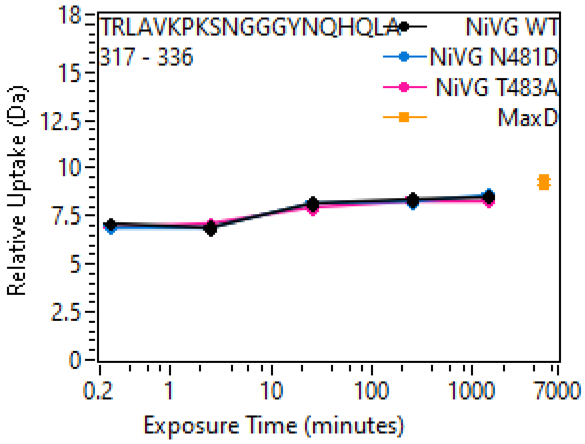
TRLAVKPKSNGGGYNQHQLA
317 - 336
NiVG WT
NiVG N481D
NiVG T483A
MaxD
Relative Uptake (Da)
Exposure Time (minutes)

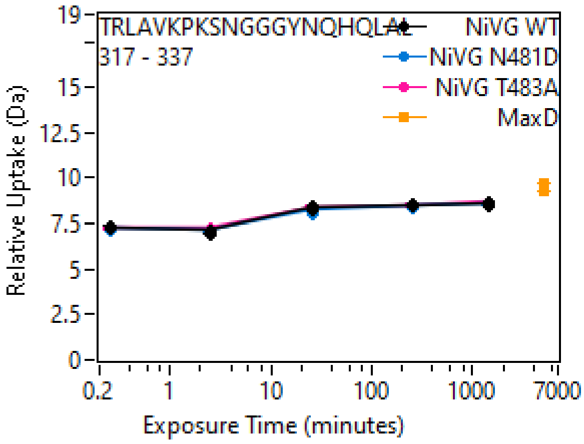
TRLAVKPKSNGGGYNQHQLAL
317 - 337
NiVG WT
NiVG N481D
NiVG T483A
MaxD
Relative Uptake (Da)
Exposure Time (minutes)

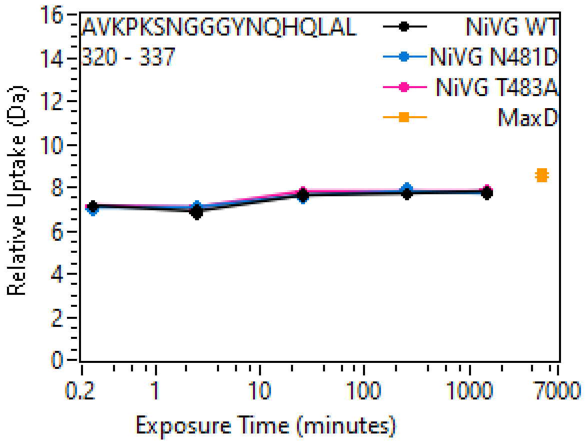
AVKPKSNGGGYNQHQLAL
320 - 337
NiVG WT
NiVG N481D
NiVG T483A
MaxD
Relative Uptake (Da)
Exposure Time (minutes)

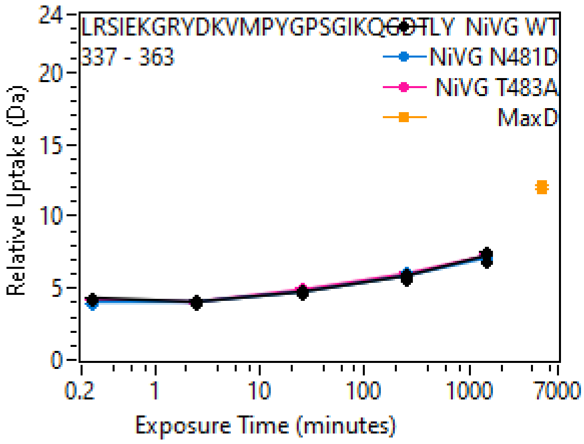
LRSIEKGRYDKVMPYGPSGIKQGDTLY
337 - 363
NiVG WT
NiVG N481D
NiVG T483A
MaxD
Relative Uptake (Da)
Exposure Time (minutes)

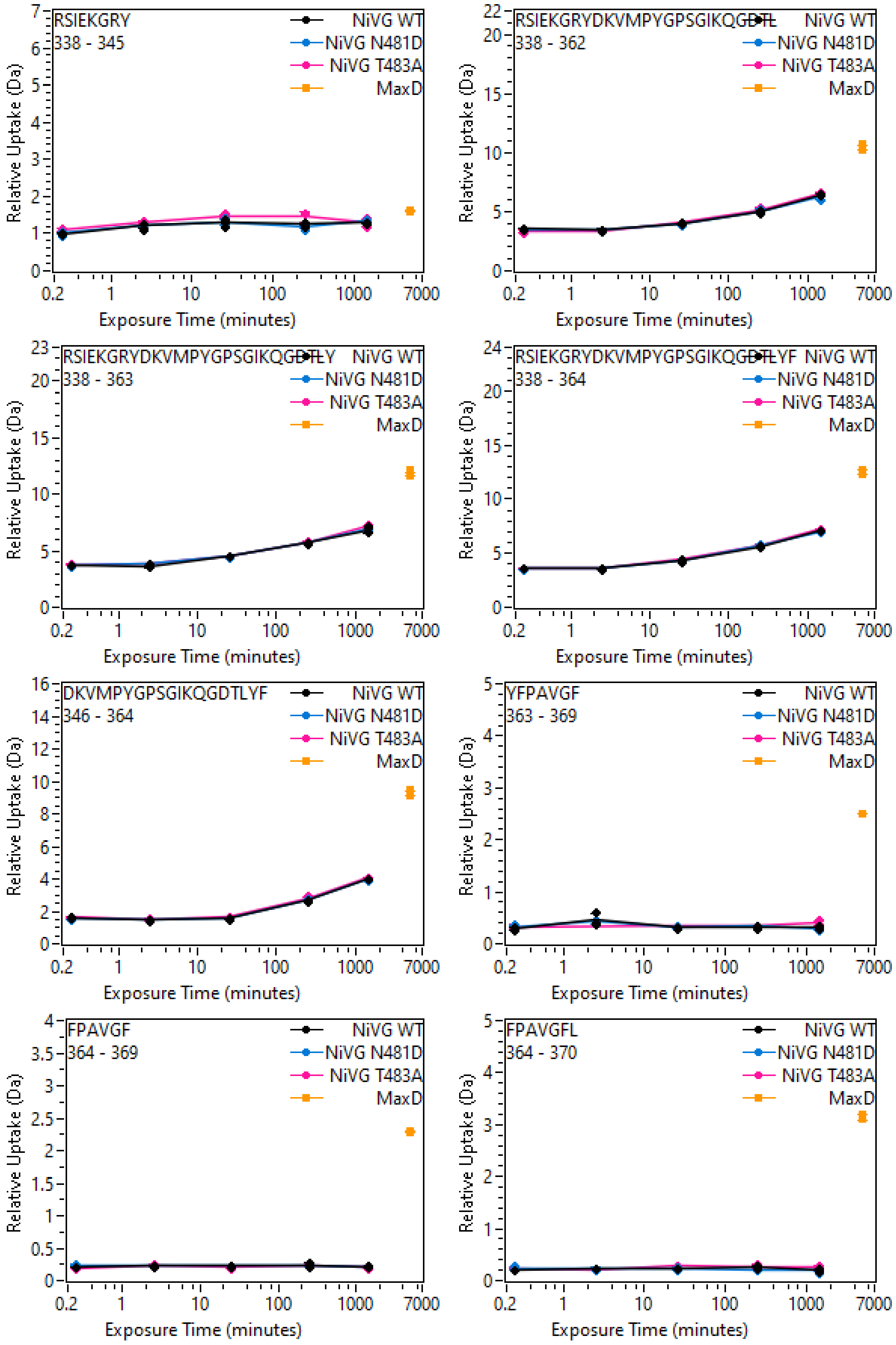
RSIEKGRY
338 - 345
NiVG WT
NiVG N481D
NiVG T483A
MaxD
Relative Uptake (Da)
Exposure Time (minutes)
RSIEKGRYDKVMPYGPSGIKQGDTL
338 - 362
RSIEKGRYDKVMPYGPSGIKQGDTLY
338 - 363
RSIEKGRYDKVMPYGPSGIKQGDTLYF
338 - 364
DKVMPYGPSGIKQGDTLYF
346 - 364
YFPAVGF
363 - 369
FPAVGF
364 - 369
FPAVGFL
364 - 370

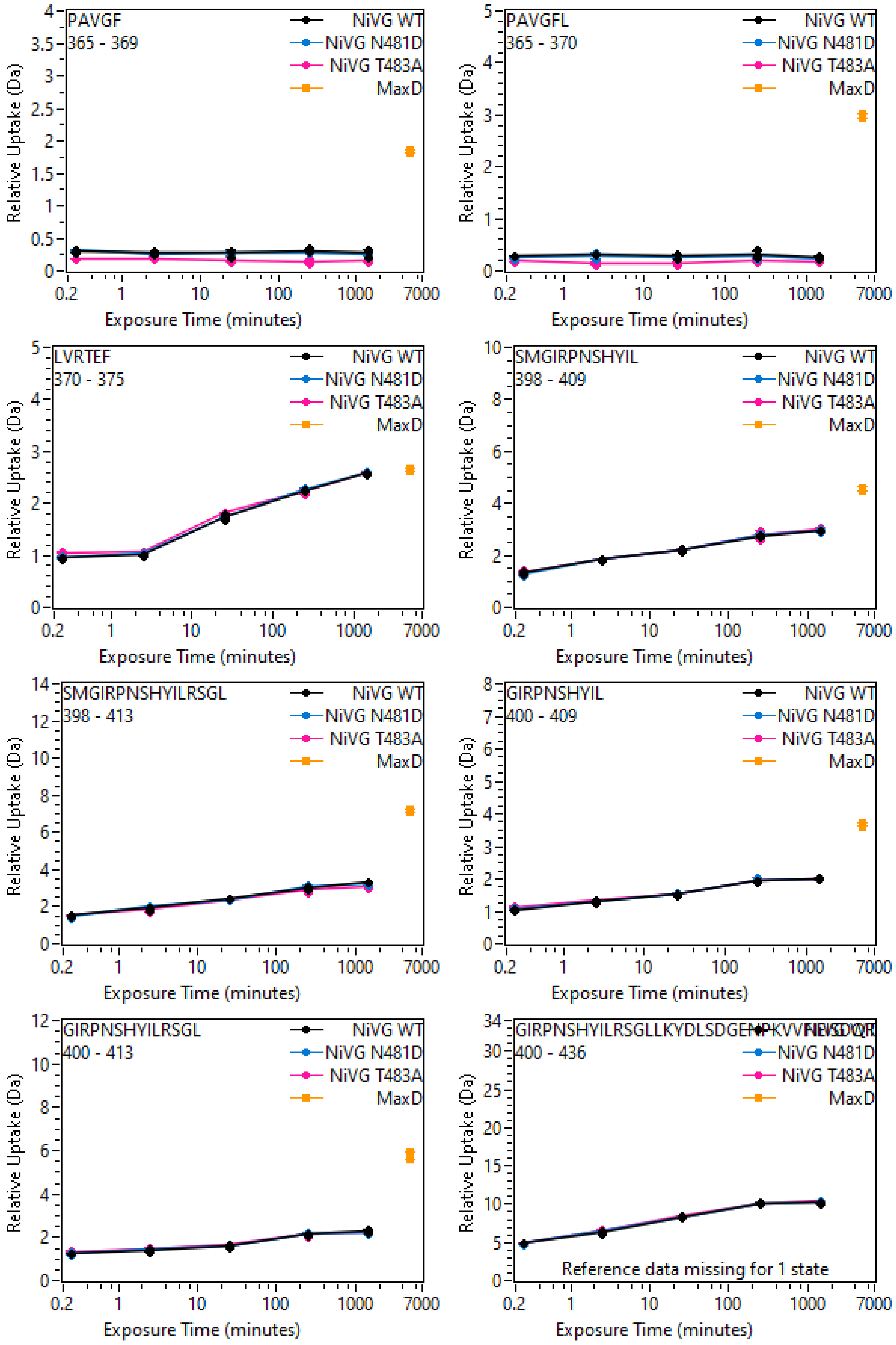
PAVGF
365 - 369
PAVGFL
365 - 370
LVRTEF
370 - 375
SMGIRPNSHYIL
398 - 409
SMGIRPNSHYILRSGL
398 - 413
GIRPNSHYIL
400 - 409
GIRPNSHYILRSGL
400 - 413
400 - 436
NiVG WT
NiVG N481D
NiVG T483A
MaxD
Relative Uptake (Da)
Exposure Time (minutes)
Reference data missing for 1 state

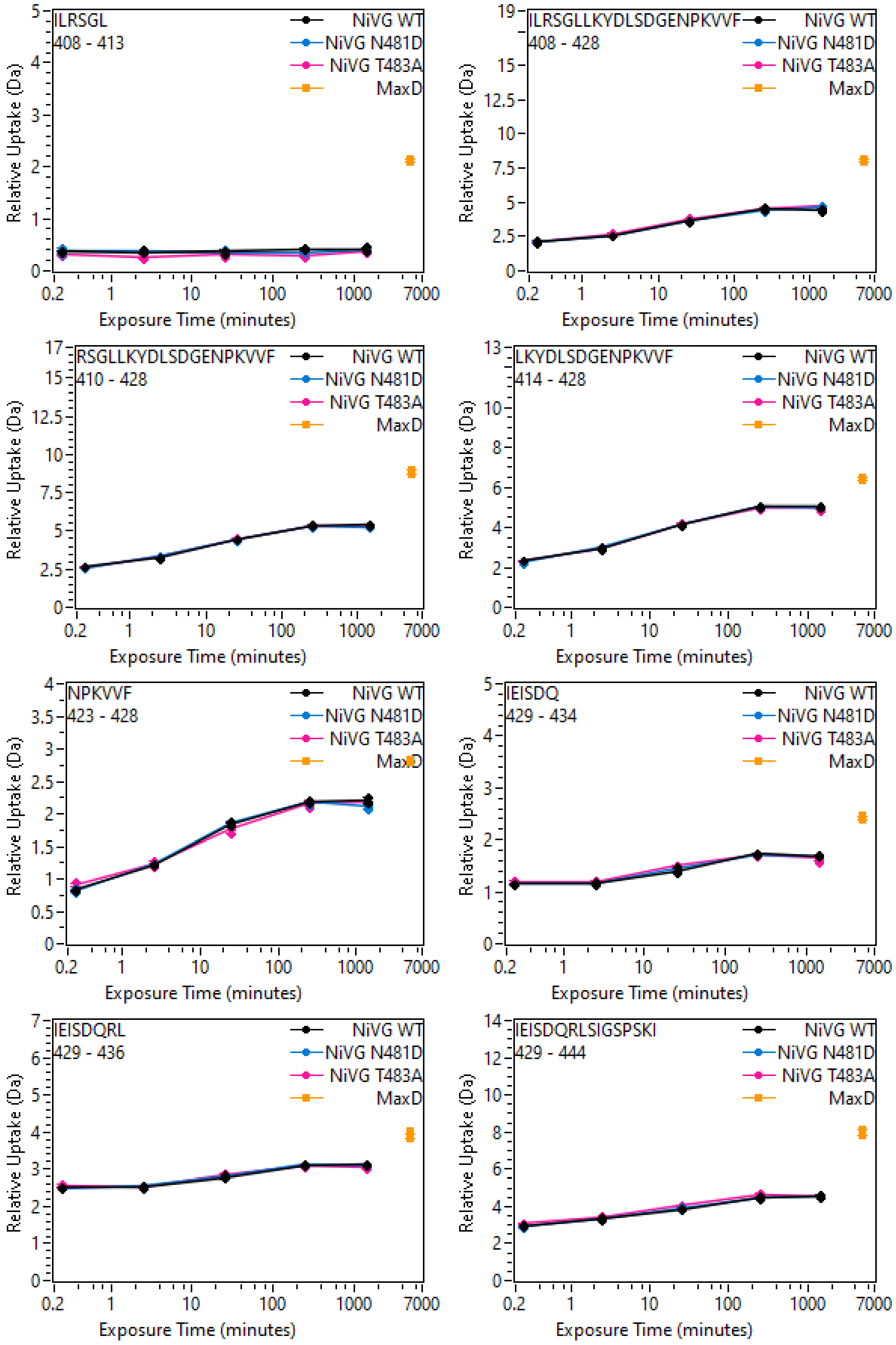
ILRSGL
408 - 413
RSGLLKYDLSDGENPKVVF
410 - 428
NPKVVF
423 - 428
IEISDQRL
429 - 436
ILRSGLLKYDLSDGENPKVVF
408 - 428
LKYDLSDGENPKVVF
414 - 428
IEISDQ
429 - 434
IEISDQRLSIGSPSKI
429 - 444
NiVG WT
NiVG N481D
NiVG T483A
MaxD
Relative Uptake (Da)
Exposure Time (minutes)
0.2
1
10
100
1000
7000

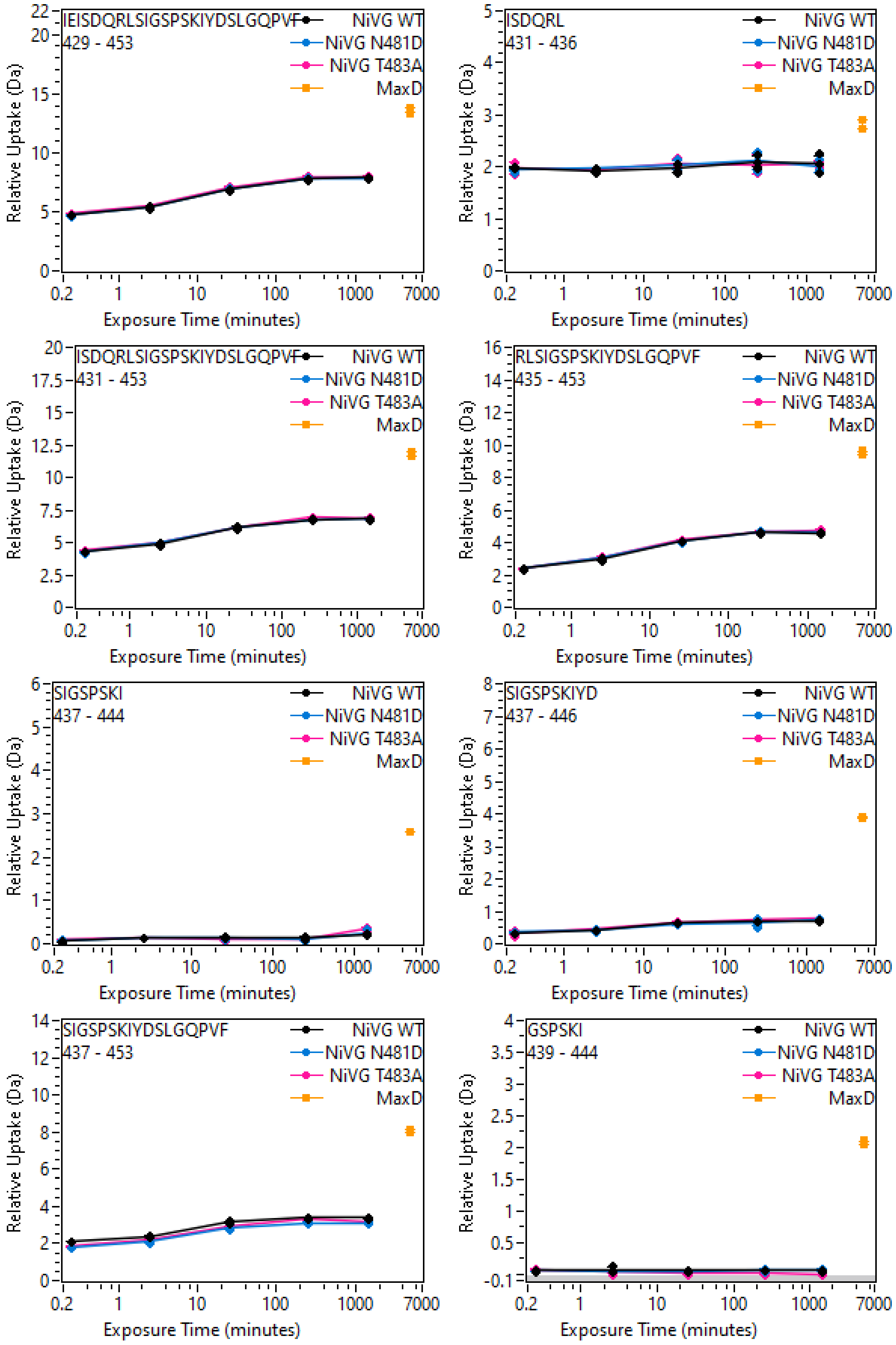
IEISDQRLSIGSPSKIYDSLGQPVF
429 - 453
NiVG WT
NiVG N481D
NiVG T483A
MaxD
Relative Uptake (Da)
Exposure Time (minutes)
ISDQRL
431 - 436
NiVG WT
NiVG N481D
NiVG T483A
MaxD
Relative Uptake (Da)
Exposure Time (minutes)
ISDQRLSIGSPSKIYDSLGQPVF
431 - 453
NiVG WT
NiVG N481D
NiVG T483A
MaxD
Relative Uptake (Da)
Exposure Time (minutes)
RLSIGSPSKIYDSLGQPVF
435 - 453
NiVG WT
NiVG N481D
NiVG T483A
MaxD
Relative Uptake (Da)
Exposure Time (minutes)
SIGSPSKI
437 - 444
NiVG WT
NiVG N481D
NiVG T483A
MaxD
Relative Uptake (Da)
Exposure Time (minutes)
SIGSPSKIYD
437 - 446
NiVG WT
NiVG N481D
NiVG T483A
MaxD
Relative Uptake (Da)
Exposure Time (minutes)
SIGSPSKIYDSLGQPVF
437 - 453
NiVG WT
NiVG N481D
NiVG T483A
MaxD
Relative Uptake (Da)
Exposure Time (minutes)
GSPSKI
439 - 444
NiVG WT
NiVG N481D
NiVG T483A
MaxD
Relative Uptake (Da)
Exposure Time (minutes)

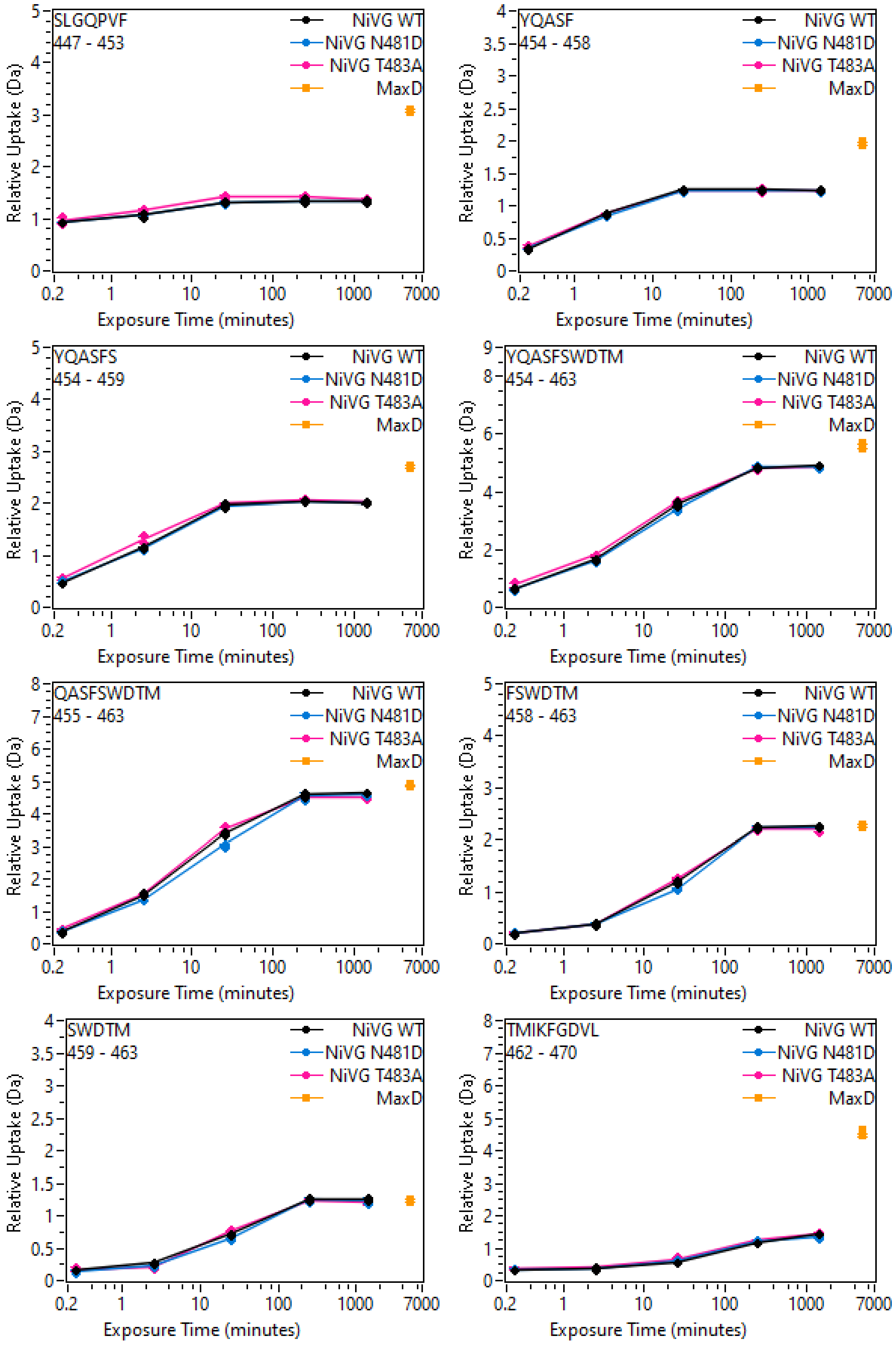
SLGQPVF
447 - 453
YQASF
454 - 458
YQASFS
454 - 459
YQASFSWDTM
454 - 463
QASFSWDTM
455 - 463
FSWDTM
458 - 463
SWDTM
459 - 463
TMIKFGDVL
462 - 470
NiVG WT
NiVG N481D
NiVG T483A
MaxD
Relative Uptake (Da)
Exposure Time (minutes)
0.2
1
10
100
1000
7000

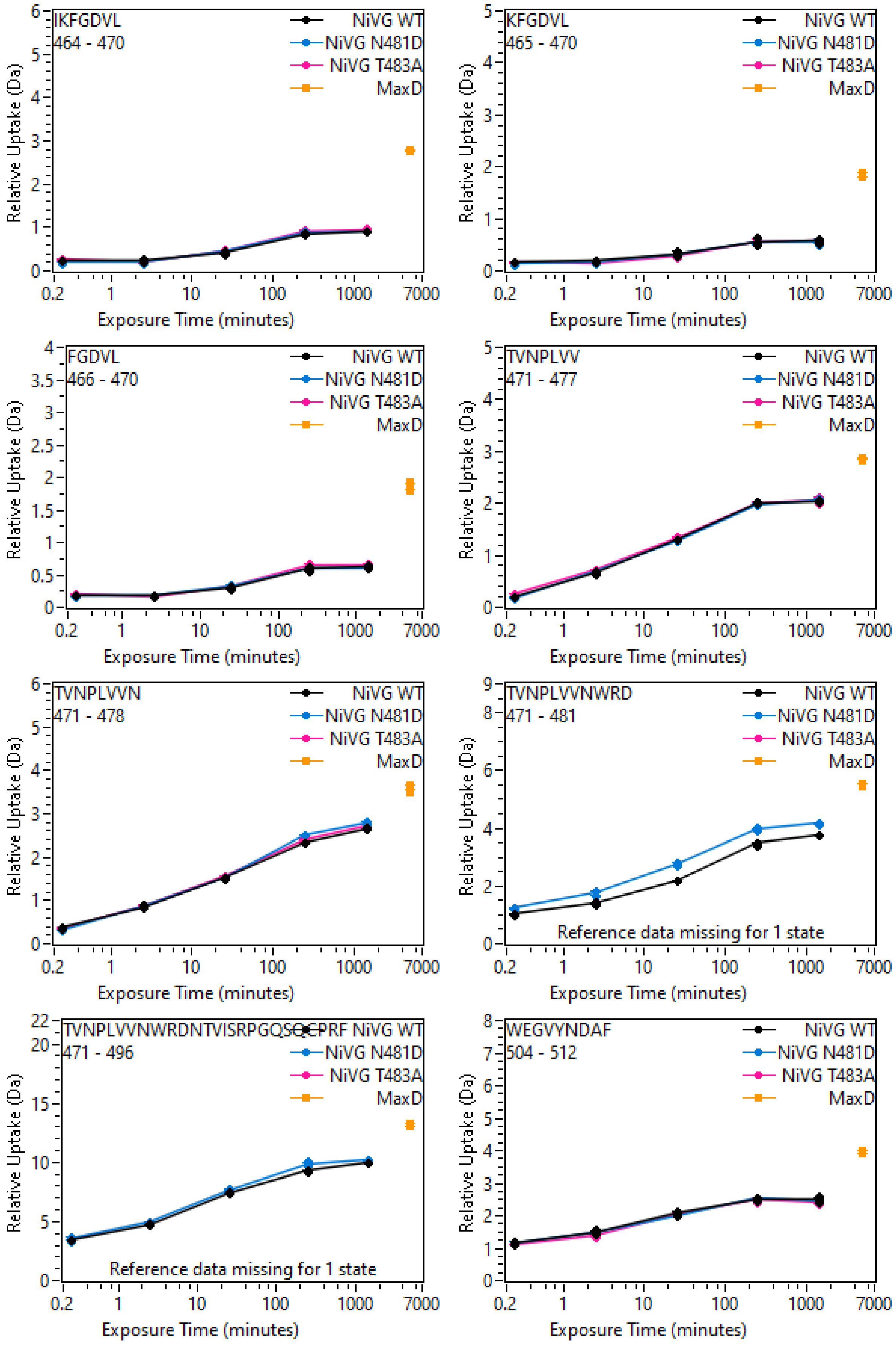
IKFGDVL
464 - 470
KFGDVL
465 - 470
FGDVL
466 - 470
TVNPLVV
471 - 477
TVNPLVVN
471 - 478
TVNPLVVNWRD
471 - 481
Reference data missing for 1 state
TVNPLVVNWRDNTVISRPGQSGCPRF
471 - 496
Reference data missing for 1 state
WEGVYNDAF
504 - 512
NiVG WT
NiVG N481D
NiVG T483A
MaxD
Relative Uptake (Da)
Exposure Time (minutes)
0.2
1
10
100
1000
7000

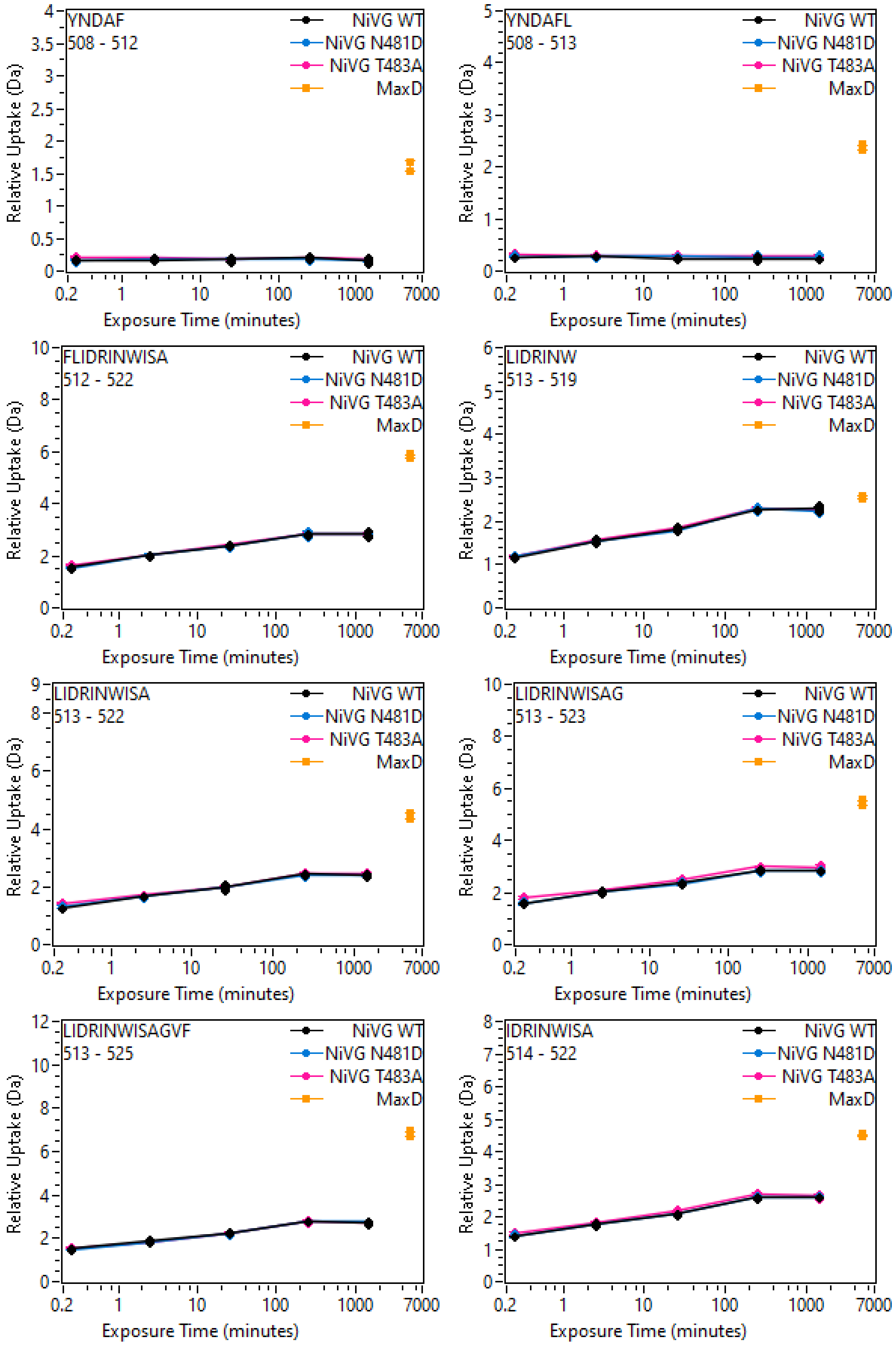
YNDAF
508 - 512
YNDAFL
508 - 513
FLIDRINWISA
512 - 522
LIDRINW
513 - 519
LIDRINWISA
513 - 522
LIDRINWISAG
513 - 523
LIDRINWISAGVF
513 - 525
IDRINWISA
514 - 522
NiVG WT
NiVG N481D
NiVG T483A
MaxD
Relative Uptake (Da)
Exposure Time (minutes)
0.2
1
10
100
1000
7000

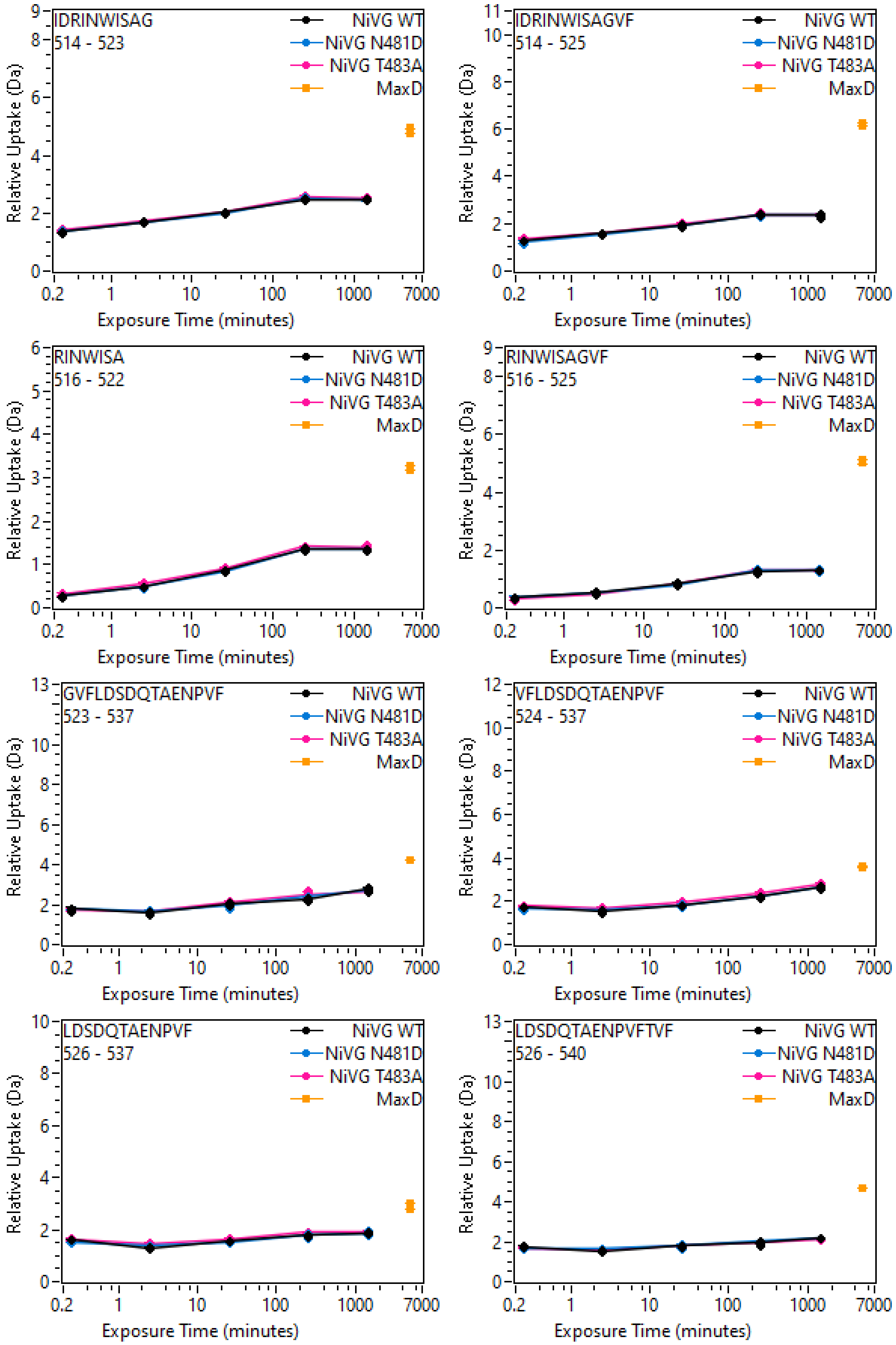

IDRINWISAG
514 - 523
IDRINWISAGVF
514 - 525
RINWISA
516 - 522
RINWISAGVF
516 - 525
GVFLDSDQTAENPVF
523 - 537
VFLDSDQTAENPVF
524 - 537
LDSDQTAENPVF
526 - 537
LDSDQTAENPVFTVF
526 - 540
NiVG WT
NiVG N481D
NiVG T483A
MaxD
Relative Uptake (Da)
Exposure Time (minutes)
0.2
1
10
100
1000
7000

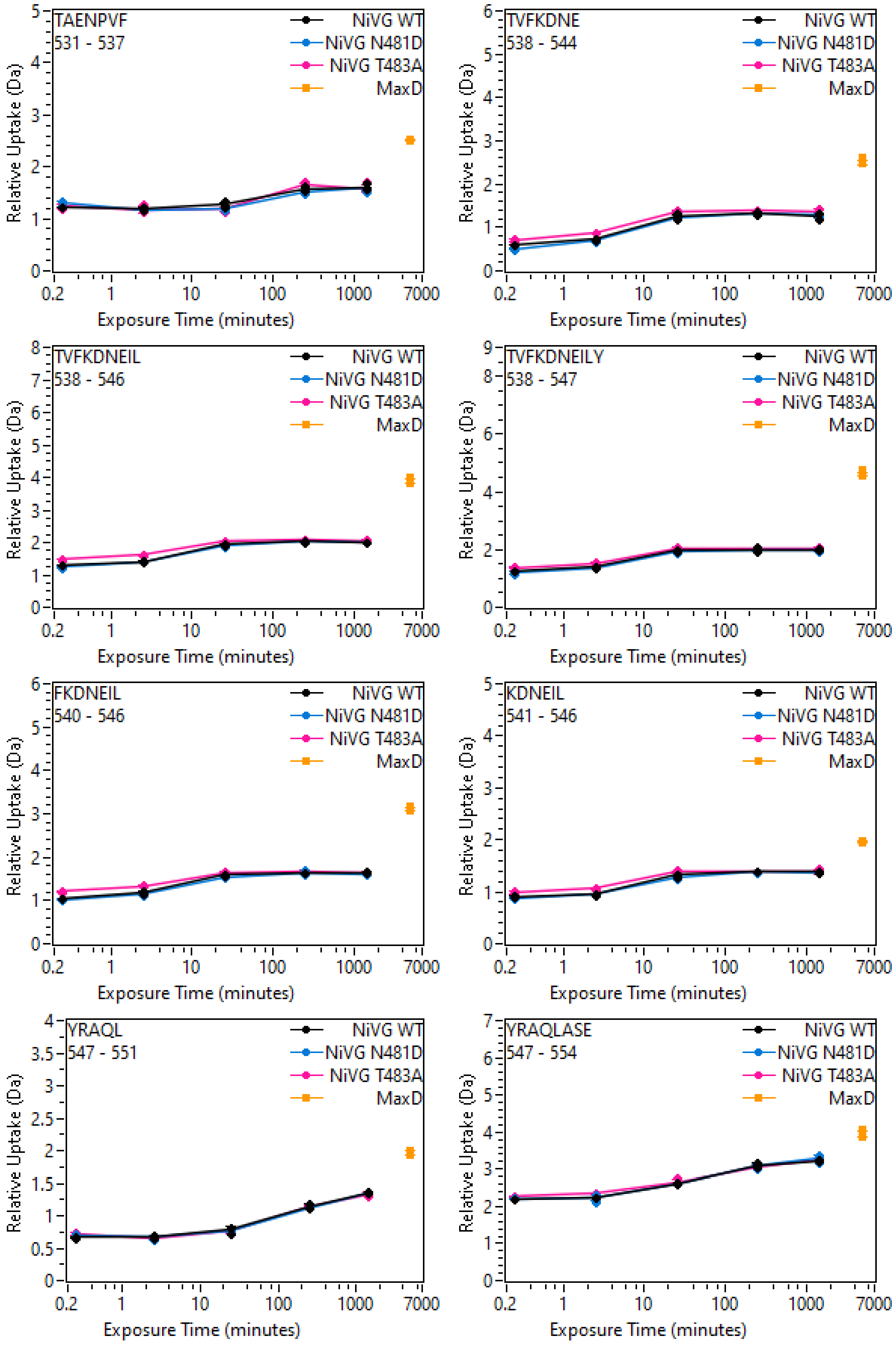
TAENPVF
531 - 537
TVFKDNE
538 - 544
TVFKDNEIL
538 - 546
TVFKDNEILY
538 - 547
FKDNEIL
540 - 546
KDNEIL
541 - 546
YRAQL
547 - 551
YRAQLASE
547 - 554
NiVG WT
NiVG N481D
NiVG T483A
MaxD
Relative Uptake (Da)
Exposure Time (minutes)
0.2
1
10
100
1000
7000

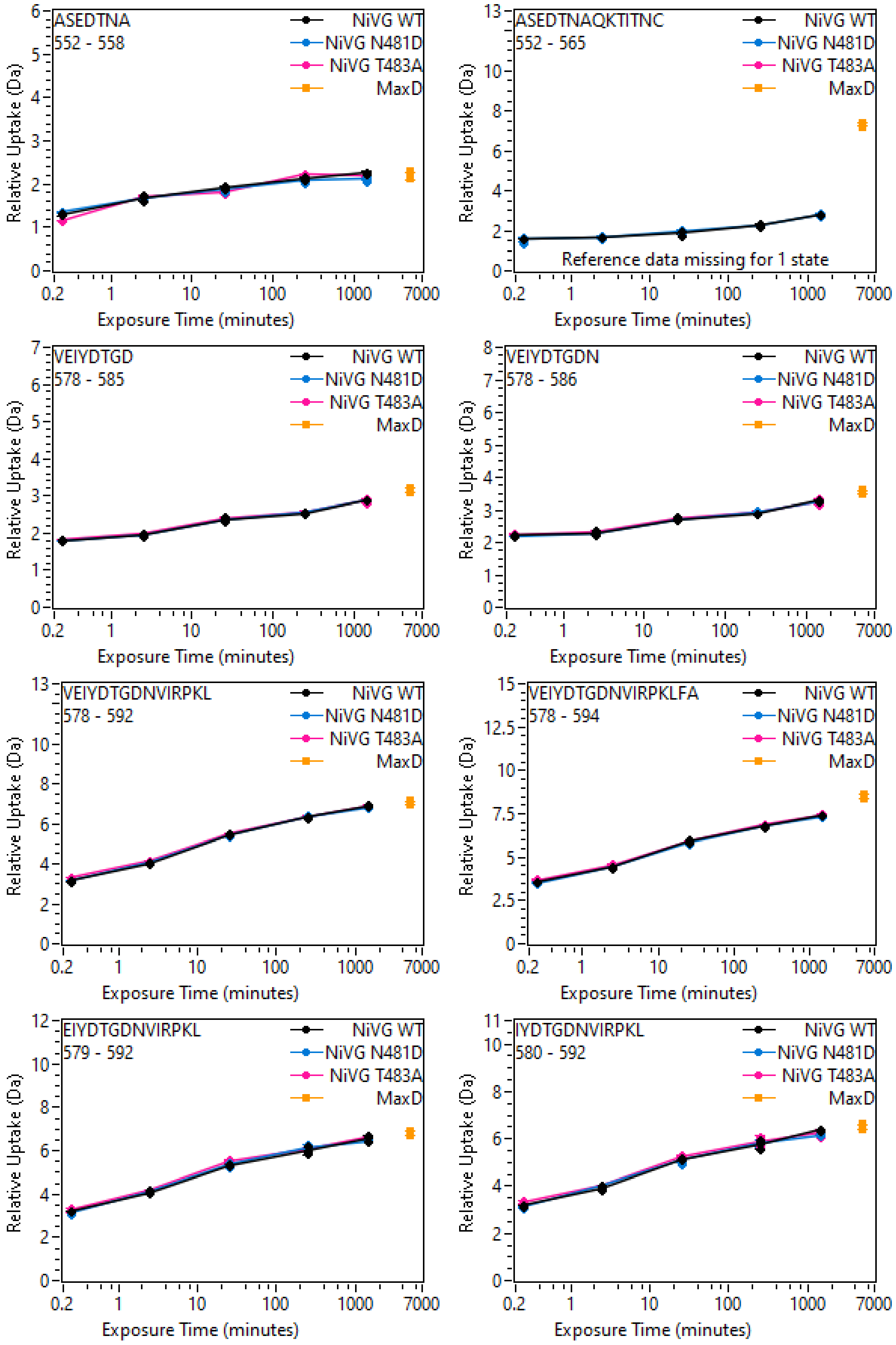
ASEDTNA
552 - 558
ASEDTNAQKTITNC
552 - 565
Reference data missing for 1 state
VEIYDTGD
578 - 585
VEIYDTGDN
578 - 586
VEIYDTGDNVIRPKL
578 - 592
VEIYDTGDNVIRPKLFA
578 - 594
EIYDTGDNVIRPKL
579 - 592
IYDTGDNVIRPKL
580 - 592
NiVG WT
NiVG N481D
NiVG T483A
MaxD
Relative Uptake (Da)
Exposure Time (minutes)
0.2
1
10
100
1000
7000

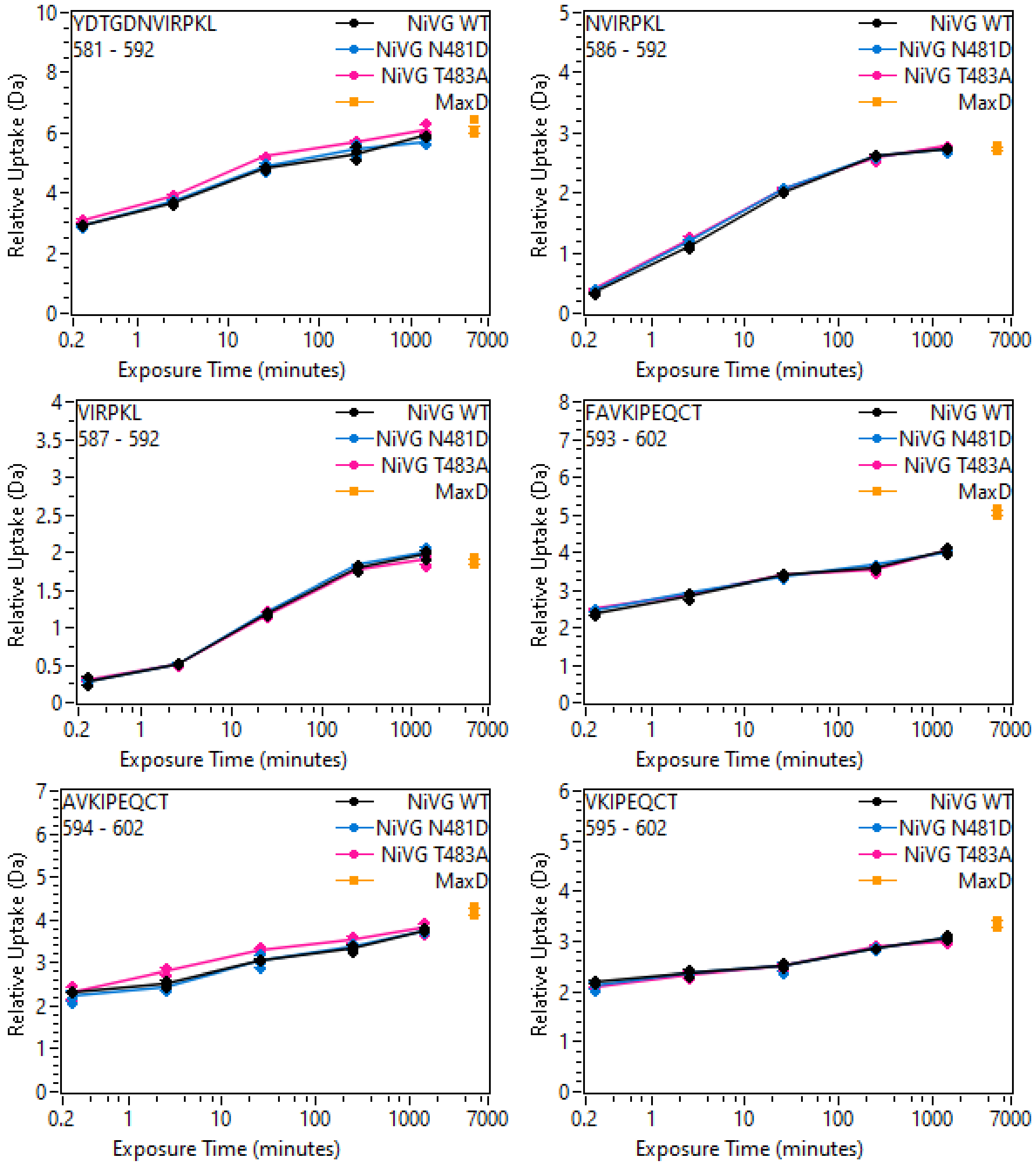
YDTGDNVIRPKL
581 - 592
NVIRPKL
586 - 592
VIRPKL
587 - 592
FAVKIPEQCT
593 - 602
AVKIPEQCT
594 - 602
VKIPEQCT
595 - 602
NiVG WT
NiVG N481D
NiVG T483A
MaxD
Relative Uptake (Da)
Exposure Time (minutes)
0.2
1
10
100
1000
7000

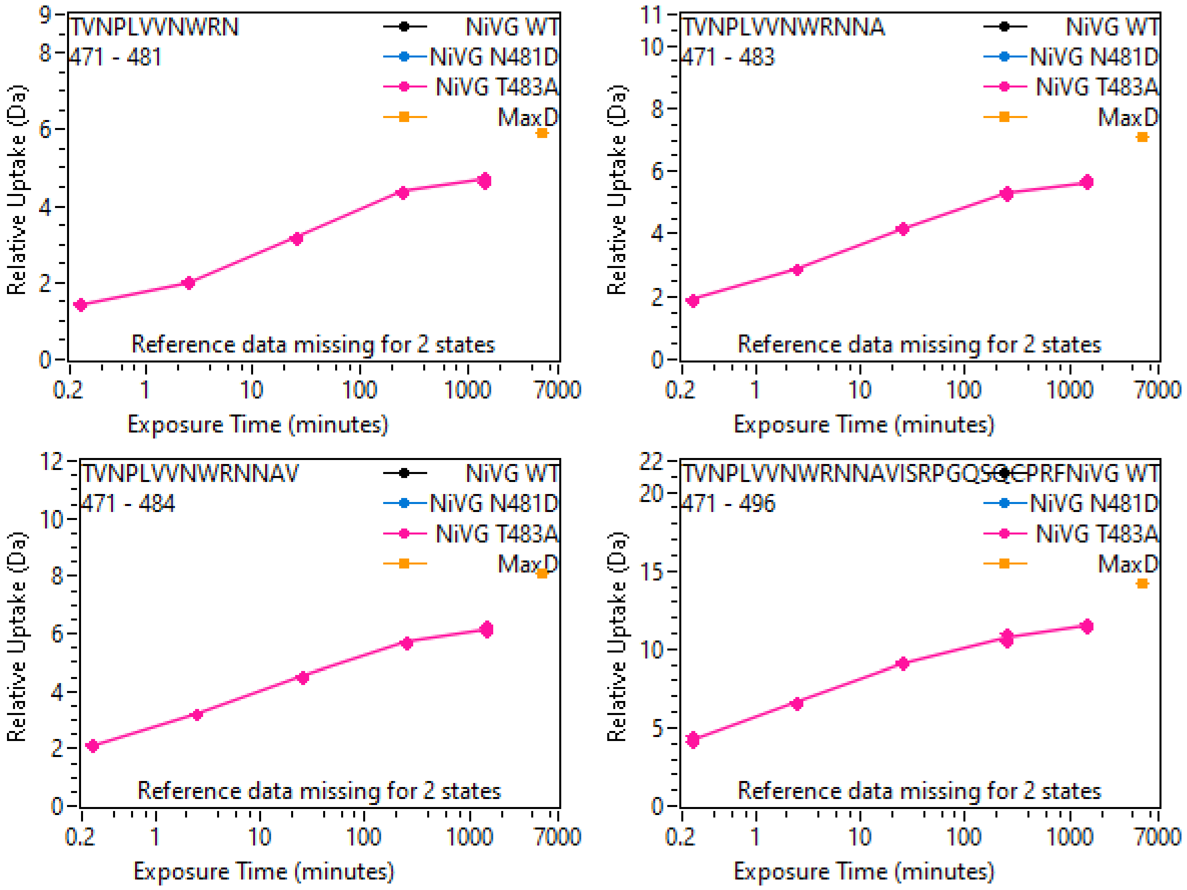
TVNPLVVNWRN
471 - 481
TVNPLVVNWRNNA
471 - 483
TVNPLVVNWRNNAV
471 - 484
TVNPLVVNWRNNAVISRPGQSQCPRF
471 - 496
NiVG WT
NiVG N481D
NiVG T483A
MaxD
Reference data missing for 2 states
Relative Uptake (Da)
Exposure Time (minutes)
0.2
1
10
100
1000
7000
